# Shedding Light on the Black Box of a Neural Network Used to Detect Prostate Cancer in Whole Slide Images by Occlusion-Based Explainability

**DOI:** 10.1101/2022.03.31.486599

**Authors:** Vojtěch Krajňanský, Matej Gallo, Rudolf Nenutil, Michal Němeček, Petr Holub, Tomáš Brázdil

## Abstract

Diagnostic histopathology is facing increasing demands due to aging populations and expanding healthcare programs. Semi-automated diagnostic systems employing deep learning methods are one approach to alleviate this pressure, with promising results for many routine diagnostic procedures. However, one major issue with deep learning approaches is their lack of interpretability—after adequate training they perform their assigned tasks admirably, but do not explain how they reach their conclusions. Knowledge of how a given method performs its task with high sensitivity and specificity would be advantageous to understand the key features responsible for diagnosis, and should in turn allow fine-tuning of deep learning approaches.

This paper presents a deep learning-based system for carcinoma detection in whole slide images of prostate core biopsies, achieving state-of-the-art performance; 100% area under curve and sensitivity of 0.978 for 8 detected false positives on average per slide.

Furthermore, we investigated various methods to extract the key features used by the neural network for classification. Of these, the technique called occlusion, adapted to whole slide images, analyzes the sensitivity of the detection system to changes in the input images. This technique produces heatmaps indicating which parts of the image have the strongest impact on the system’s output that a histopathologist can examine to identify the network’s reasoning behind a given classification. Reassuringly, the heatmaps identified several prevailing histomorphological features characterizing carcinoma, e.g. single-layered epithelium, presence of small lumina, and hyperchromatic nuclei with halos.

A convincing finding was the recognition of their mimickers in non-neoplastic tissue. The results show that the neural network approach to recognize prostatic cancer is similar to that taken by a human pathologist at medium optical resolution. The use of explainability heatmaps provides added value for automated digital pathology to analyze and fine-tune deep learning systems, and improves trust in computer-based decisions.

## 1 Introduction

The increasing lifespan in developed countries inevitably leads to higher incidences of cancer due to the aging population. Together with the expansion of cancer screening programs and personalized medicine, this results in an increased workload for healthcare systems, including diagnostic specialties such as radiology and histopathology. This effect is in part compensated by progress in digitization, providing more efficient processing, archiving and retrieval of medical records. The use of digitized medical images represents the next evolutionary step. Radiology is more advanced in this respect, already routinely utilizing picture archiving and communicating systems, while comparable pathology systems utilizing whole slide images (WSI) are currently being approved for diagnostic use [1] and introduced into routine workflows [2].

Extensive improvements of digitized WSI provided the base for developing more advanced approaches, including the use of neural networks for image analysis to facilitate diagnostics and prognostics, or simply to assist pathologists to reduce their workload in routine tasks [3– 5]. The detection of prostate cancer in core biopsies is one example of such an application. This represents a typical and often tedious part of a pathologist’s daily routine, where samples from patients identified as prostate specific antigen positive during screening are mostly negative, but all slides, often containing multiple tissue sections, must be carefully evaluated for small cancer foci. Unsurprisingly, some of the first deep learning applications in WSI analysis were dedicated to this problem, providing impressive results [6–8].

The deep learning approach is sometimes criticized for not providing insights into the mechanism of classification, which raises issues of trust in the method by the pathologist as well as hampering improvements of the algorithm and a lack of transparency may hinder acceptance by regulatory authorities [9, 10]. In addition, unravelling the precise features used by the system could provide additional information that would be useful for pathologists and for further training of the machine learning process itself. Although some successful attempts have been shown, e.g. the use of probability heatmaps to identify the histopathological features associated with adverse prognosis in glioma [11], most reports largely ignore this aspect and focus on the application of the method rather than identifying how it operates in its decision making processes. In contrast, by its nature, deep learning may reveal new information not directly attributable to known histomorphological characteristics [12]. To unravel such information from the complex “black box” of artificial intelligence, we need the tools to explain what is important for the deep learning decision-making in histopathological images.

Core biopsies of prostate represent a showcase of deep learning network’s performance to recognize cancer. From the pathologist’s point of view, the morphological criteria of malignancy in prostate are well established and relatively reproducible. In this paper we tested various approaches to this problem and present the application of occlusion-based explainability to identify the key morphological features important for the deep learning decision of prostate core biopsies.

## 2 Material and Methods

### 2.1 Material

The WSI stained with hematoxylin/eosin (containing 3 to 5 tissue core sections each) are part of the digital archive at the Department of Pathology, Masaryk Memorial Cancer Institute, Brno. They were scanned using a Pannoramic® MIDI (3DHistech, Budapest, Hungary) scanner with 20× objective lens at a resolution of 0.172 μm / pixel. The WSI were stored in MIRAX format, as uncompressed PNG images. The dataset consisted of:

1. learning WSI, obtained from 156 consecutive core biopsies (103 cases with carcinoma, 53 negative), in total 262 WSI with cancer, 436 without cancer;
2. test WSI, obtained from 10 cases with cancer, selected from additional consecutive scans to represent different Gleason patterns and types of infiltration, in total 37 WSI with cancer, 50 without.

Details and Gleason pattern distribution are given in supplementary data (Supplementum Table S1.1).

The WSI were checked in the automated slide analysis platform ASAP^1^ [13] and all biopsy cores containing carcinoma were manually annotated in ASAP for further analysis. Annotations were performed as polygons, containing carcinoma areas. The whole process is presented as a flowchart in Figure 1.

**Figure 1:**
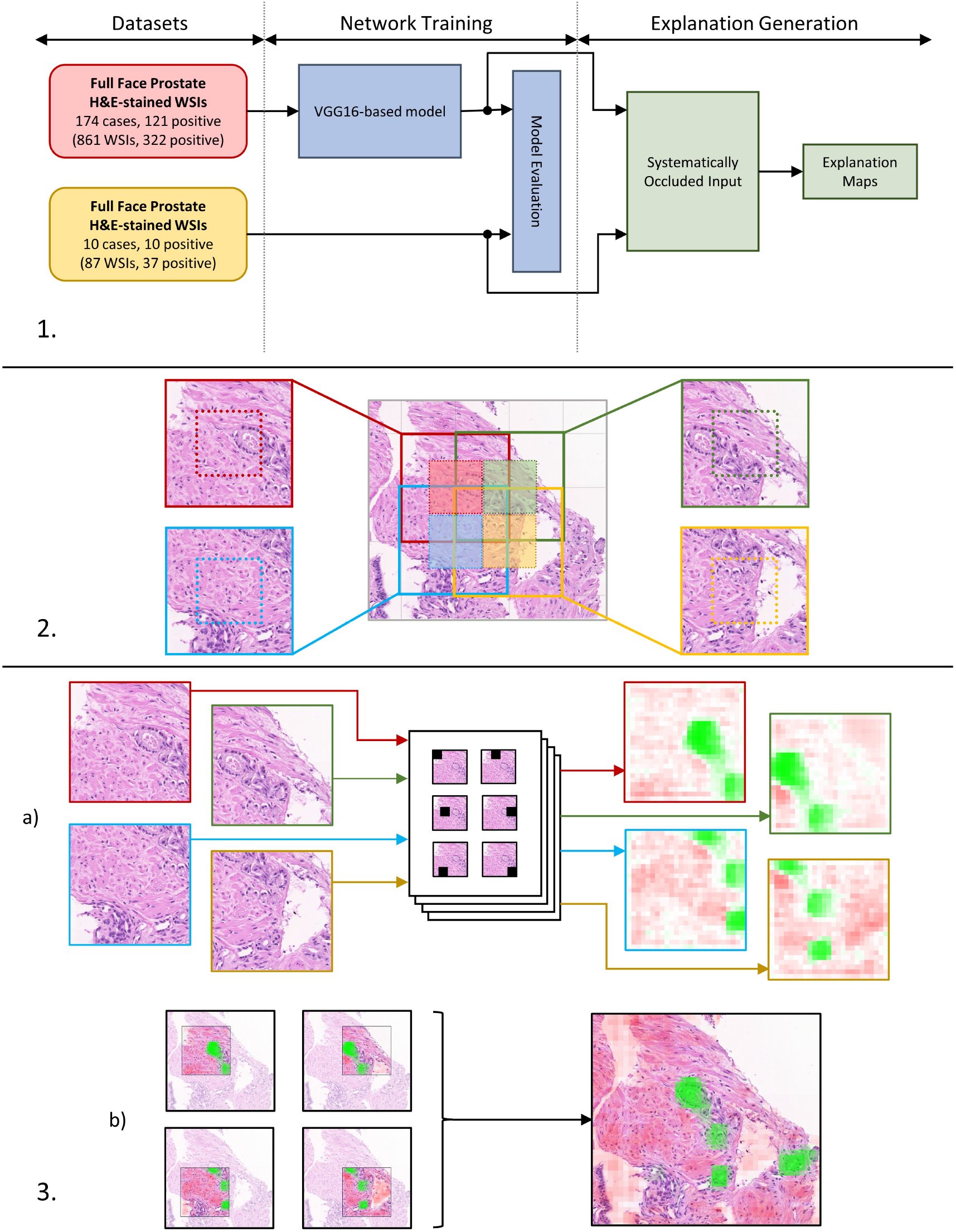
Network training and explanation generation. Section **1** shows the overall training and generation process; Section **2** illustrates how input patches overlap as a consequence of including surrounding contextual information (the whole patch) around the label-relevant central region (denoted by dotted squares); Section **3a)** illustrates the systematic occlusion of individual patches and resulting explanation maps; Section **3b)** shows how a WSI overlay is obtained by averaging the overlapping explanation maps.

### 2.2 Data set access

The data set is available as raw files stored in Mirax MRXS format^2^ compatible with the OpenSlide library [14]. Annotations used for the evaluation stage are available as XML files compatible with ASAP [13]. The data set is pseudonymized and access can be requested via BBMRI-ERIC European Research Infrastructure by following its access policy^3^; the request should be placed via BBMRI-ERIC Negotiator platform^4^ to Masaryk Memorial Cancer Institute.

### 2.3 Network training

In this section we describe our training of neural networks on WSI, which mostly follows the well-established approach used in several deep learning on WSI papers, e.g. [15].

Our dataset comes from 785 WSI of 166 patients. We used 698 WSI of 156 patients for training, leaving out 87 WSI of 10 patients for the blind test set. Due to their large size (105, 185 × 221, 772 px), the slides were cut into smaller patches of size 512 × 512 px. Thus, our training set consists of 7,878,675 patches, with 193,235 patches for testing.

#### Slide Preparation

First, we restricted the sampling process only to tissue areas. All patches whose tissue coverage did not exceed a threshold of 75% were ignored. To assign labels to the remaining patches, we used the annotation binary mask provided by pathologists. A patch was assigned a positive label if its center area of size 256 × 256 px contained at least a single pixel belonging to the cancer category, similar to [15, 16] (see Figure 1).

#### Input Preprocessing

The dataset is highly imbalanced. The training set contains over 8 million negative patches and over 1.5 million positive patches. To combat this imbalance and to improve generalization, we applied image augmentation techniques to artificially enlarge the dataset, such as random vertical and horizontal flips with 50 % probability, random brightness, hue, saturation and contrast perturbations, etc. [15–17]. We used the *imaug* package to augment the patches. Finally, all patches were scaled to [−1,1] range.

#### Patch Sampling

During training, a three-step sampling process is used to deal with the imbalanced dataset [15]. In the first step, a label *L* is selected at random. In the next step, a single slide *S* is picked at random with uniform probability from all the slides containing at least one patch with label *L*. In the last step, a patch with label *L* is selected uniformly at random from the slide *S*. This procedure also ensures that patches are drawn from all slides equally, regardless of how much tissue area or cancer area is present in the slide.

#### Network Architecture and Training

We used a VGG16 network pretrained on the *imagenet* dataset for feature extraction. The model was trained using an RMSprop optimizer with the following parameters: momentum = 0.9, *ϵ* = 1.0, *ρ* = 0.9, initial learning rate of 5 × 10^−5^ and batch size of 1. Learning rate was halved and training was terminated if no improvement on the validation data was made for 5 and 10 consecutive epochs, respectively.

### 2.4 Explainability using occlusion sensitivity analysis

In our research, we tried several approaches to interpretability such as occlusion sensitivity analysis [18], saliency maps [19], and LIME [20]. To our surprise, the simplest of these methods, occlusion, showed the highest fidelity, and was the most accurate for tumor detection.

#### Occlusion Mask Generation

Occlusion sensitivity analysis is based on the idea of systematically perturbing the input and observing the corresponding changes in the model’s output. Specifically, we systematically covered (occluded) square regions of the input by setting their values to zero, and recorded the output in a matrix, whose value positions correspond to the relative positions of the covered region in the input. This matrix can then be upsampled to obtain a mask covering the whole original input. The process is summarized in Figure 1.

After computing the occlusion matrix, it may not immediately be clear how to visualize it. An obvious approach is to scale the values to some desired range (e.g. [0, 255] for pixel values) and directly visualize the prediction values on the corresponding regions for the perturbed input. However, using this method we would lose information of whether occluding a specific region leads to an increase or decrease in the model’s response. To mitigate this, one might instead visualize the (scaled) differences between the original class score and the score obtained on a perturbed image. When the difference is positive (i.e., occluding a patch results in a lower response from the network), this signifies a region of positive influence towards the class score, and vice versa. However, this approach poses another problem specific for classification tasks. The sigmoidal output of the model may be saturated (very close to one or zero), which means that even if the network itself is strongly sensitive to an input perturbation, the change in the output value may be very small. To mitigate this problem, we do not record the differences of the sigmoid predictions themselves, instead taking the differences of logits, i.e., values that are fed into the output sigmoid.

This leaves us with a problem of values unrestricted in range. This means that there might exist wildly different ranges of values for different patches. As our goal was to obtain a single occlusion map for the whole WSI, we needed to normalize the differences into the same range across all patches. We solved this problem by feeding the differences of logits into a logistic sigmoid, thus obtaining relative differences in network reactions normalized into the interval [0, 1].

Formally, assume that *σ*(*f*(*x*)) is the function computed by our network, where *σ* is the output activation function (the logistic sigmoid) and *f*(*x*) computes the logit value from the input *x*. Now given an image *I* and its perturbation *I*′ obtained by occluding a region, our *occlusion score* is:

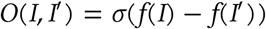

We applied occlusion sensitivity to our 512 × 512 px patches, with the occluded region size set to 55 ×55 px. With a stride of 25 px, this gives masks of size 20 ×20 px, that are subsequently upsampled to 512 × 512 px masks using nearest-neighbor upsampling. Note that this means 20 × 20 perturbed inputs are generated to obtain an occlusion mask for a single input image. As such, occlusion sensitivity analysis is quite inefficient terms of performance (see supplementary data; Section S1.3 for more information). The resulting masks have been stitched together to form a continuous occlusion mask over the whole WSI (see Figure 2).

**Figure 2:**
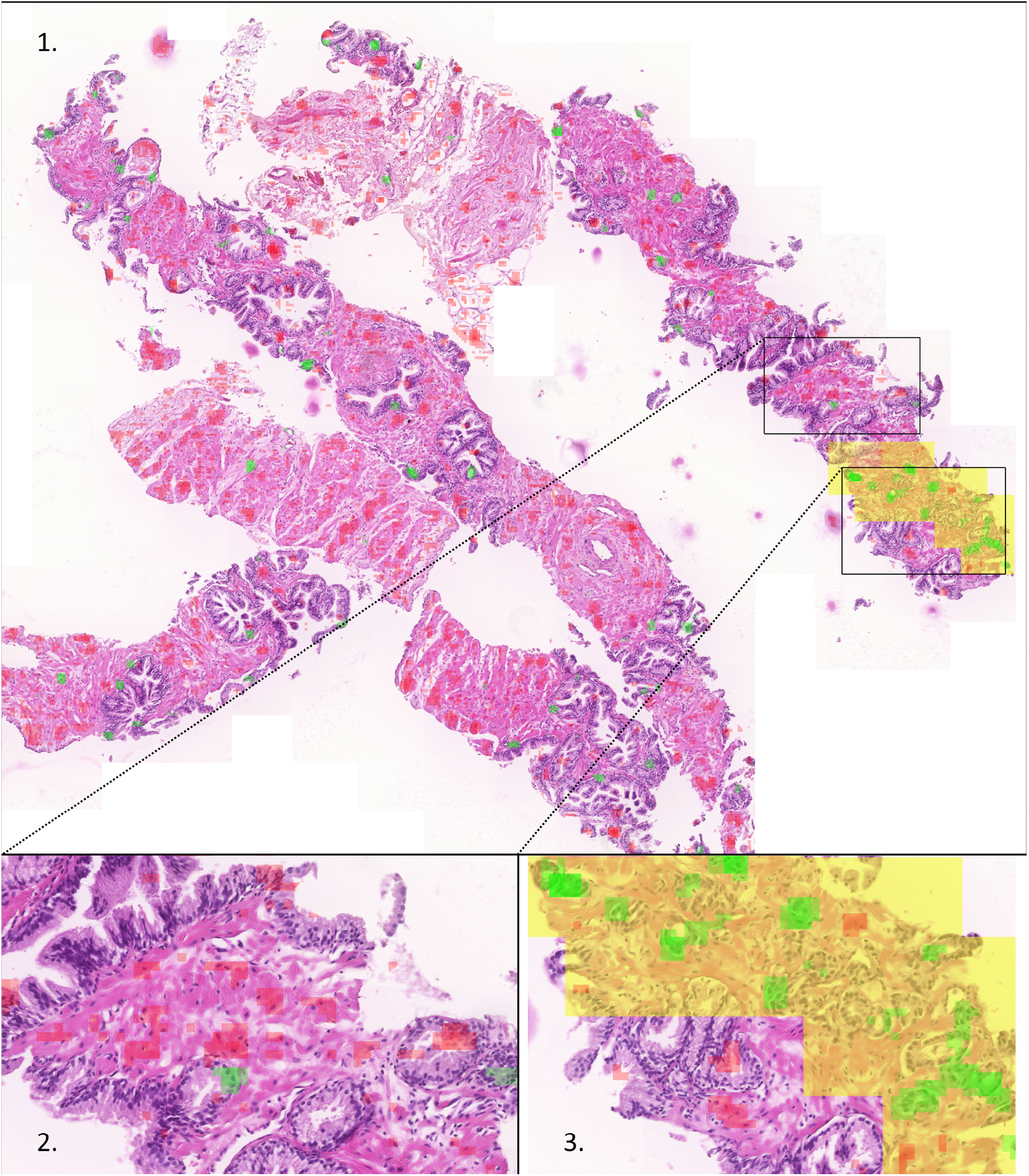
Network predictions and occlusion explanation overlays. The prediction layer (yellow) labels the focus of Gleason pattern 3 carcinoma. The explanatory layer (green and red) labels those parts of image critical for estimation of carcinoma probability using the occlusion method. Regions supporting the classification of a tile as malignant are green, while those suppressing such a classification are red. The details below show parts of a WSI with carcinoma (right) and without carcinoma (left).

Given the nature of WSI preprocessing, in which the output label only refers to the input’s center area, with the surrounding region provided as contextual information, each pixel of a single input WSI is present as part of 4 overlapping inputs, appearing once as a center pixel, 3 times contextually. It follows that each pixel is also assigned 4 distinct *occlusion scores* during the sensitivity analysis. For ease of visualization as well as a means of regularization, the overlapping *occlusion scores* are blended using the arithmetic mean. Generally, given *n* overlapping explanation masks, the final mask may be computed as:

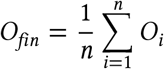

#### Sampling Regions for Explanation Evaluation

In order to evaluate the semantic strength of the generated explanatory maps, they were evaluated on 647 randomly selected samples of tissue regions from 86 test WSI (37 positive and 49 negative). We used a square grid of side length 280 px, and chose an intersection as a sample point if the absolute difference between the mean positive and negative scores in the surrounding 15×15 px square was greater than 70. This way we ensured that only unambiguously positively or negatively contributing regions were evaluated. We also limited the maximum number of samples from a single WSI to 20.

The mean sample count for a single slide was approximately 7.5, with extremes being 20 samples from 1 slide, and 1 sample in the case of two slides.

## 3 Results

### 3.1 Results for classification

We have adopted the evaluation metrics used in the CAMELYON16 [21] competition, namely, tumor-level area under the free-response receiver operating characteristic (FROC) score [22] and slide-level area under curve (AUC) score. This is because patch-level metrics such as precision and recall cannot adequately represent the quality of a trained model in this case due to two factors: firstly, not all patches need to be classified correctly for a tumor to be found; secondly, due to the complex nature of the annotation process, the slides might not be perfectly annotated to the pixel. Figure 3 shows two cases in which the network response on carcinoma tissue is relatively low.

**Figure 3:**
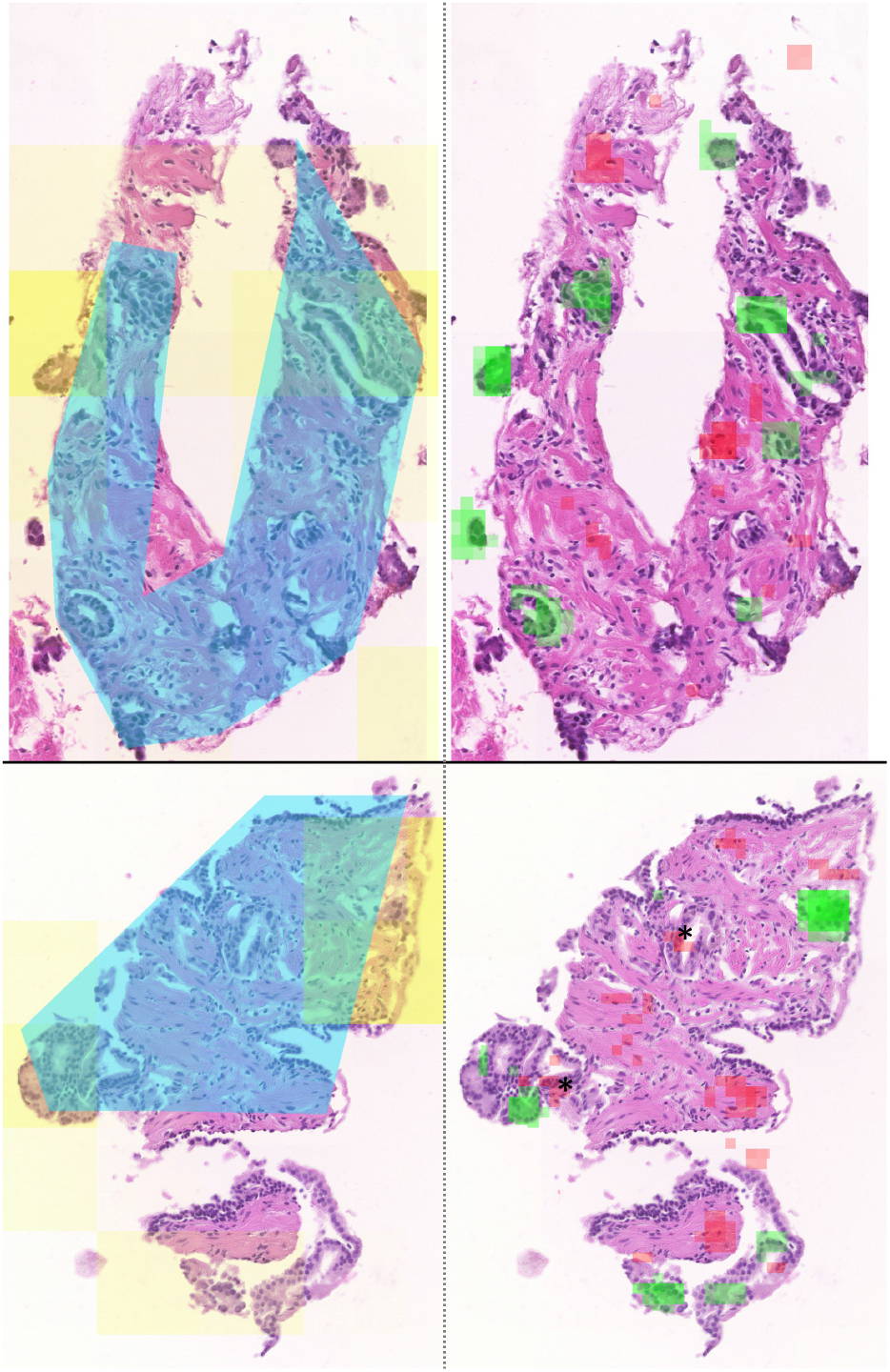
Examples of FN misclassification (low network response). Top: Part of carcinoma with sparse tumor glands infiltrating fibromuscular stroma. Although carcinoma glands are labeled correctly, the strong and frequent negative labels in stroma seem to push the decision to negativity. Bottom: A similar problem (few tumor glands and negative stroma dominates). Moreover, two points of explanation type FN1 (two layered epithelium in carcinoma) are present, denoted by asterisks. (Yellow: cancer prediction, blue: manual annotation, green and red: positively and negatively contributing regions, respectively.)

The FROC score helps to assess the ability of a model to identify tumors. On the other hand, the AUC score measures the model’s ability to identify patients with cancer. Similar to ROC, the FROC score is a plot of sensitivity versus the average number of false positives in negative slides. Sensitivity is taken for 6 predefined false positive rates: 1/4, 1/2, 1, 2, 4 and 8 average false positives per slide [15, 21].

To calculate FROC scores, we used the non-maxima suppression method [15, 17]. Because our patches have a center size of 256 px, we accordingly selected the radius *r* to 256 px. Figure 4 shows the FROC curve and the associated score, and Table 1 shows sensitivity for the 6 predefined points. We applied a max-pooling strategy on the patches to select the slide scores. We achieved 100% AUC with this method.

**Table 1:** FROC score for the 6 predefined FP rates.

| FPS/WSI | 1/4 | 1/2 | 1 | 2 | 4 | 8 |
| --- | --- | --- | --- | --- | --- | --- |
| Sensitivity | 0.854 | 0.879 | 0.916 | 0.934 | 0.954 | 0.978 |

**Figure 4:**
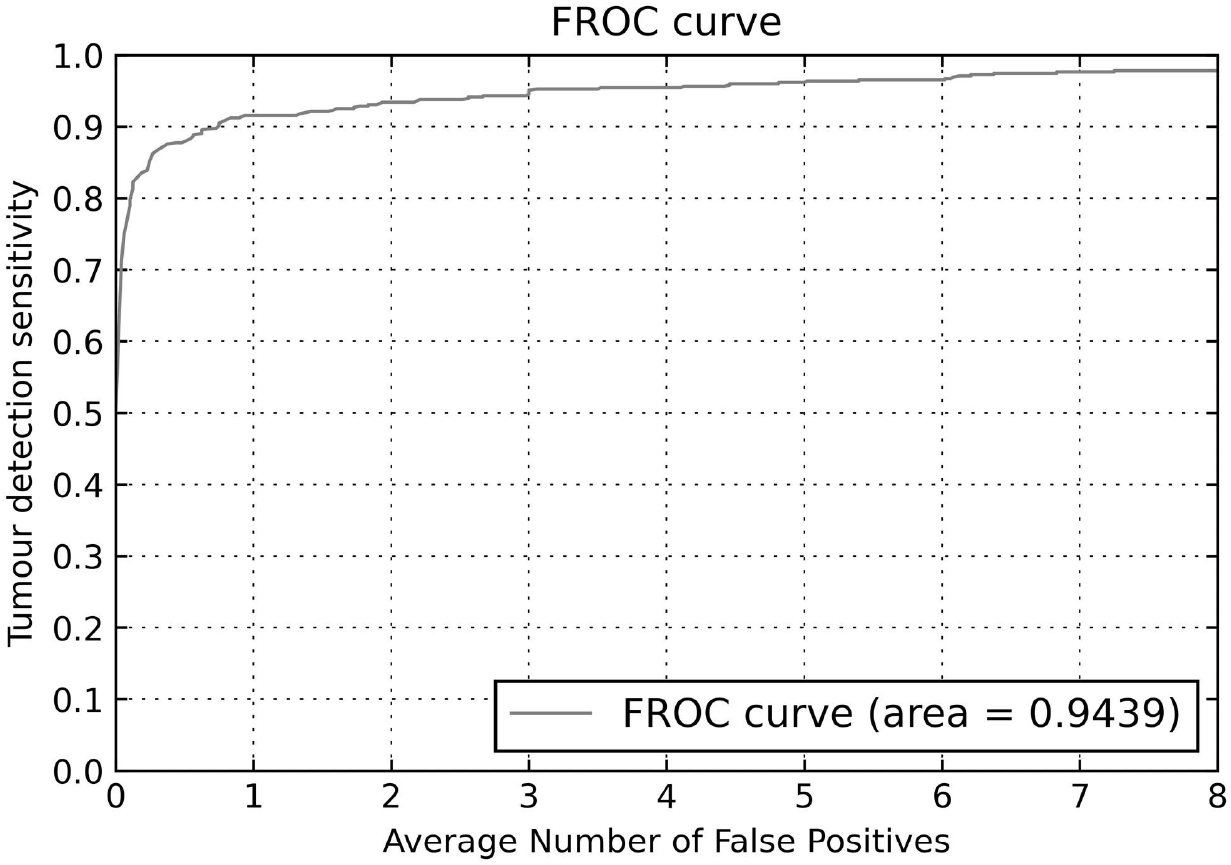
Plot of FROC curve evaluated on test set.

### 3.2 Results for explainability

The explainability results were presented to a pathologist in a web-based environment, which shows the results of classification and explainability in the form of transparent overlays at the presented WSI (Figure 2, Supplementary Figure S2.3 in supplementary data). First, these results were checked to seek for possible repeating patterns forcing the classification of the tile as suspicious of cancer or negative of cancer in a wider context. Seven such patterns were found (Table 2, Figure 5, Supplementary Figure S2.2).

**Table 2:** Distribution of sampled explainability patterns in the test set of WSI. Single layered epithelium represents the predominant true positive pattern (52.2% of true positive sample points). Note the change of pattern distribution related to the Gleason score in carcinoma tissue (underlined, in italics), which represents some internal control of reading validity. In false positive patterns, there is no obvious predominance. In true negatives, the low cellular density, in most of cases related to the stromal component predominates (25% of total sample points), followed by two layered epithelium (11.1% of total sample points). The undefined patterns represent 2% of total sample points.

| Pattern<br>Description | WSI w/ carcinoma |  |  |  |  |  |  | WSI w/o<br>carc.<br>N=49 | Total | Tot. % |
| --- | --- | --- | --- | --- | --- | --- | --- | --- | --- | --- |
|  | Gleason |  |  |  |  | Total<br>(N=37) | TP% |  |  |  |
|  | 3+3<br>(N=14) | 3+4<br>(N=3) | 4+3<br>(N=11) | 4+4<br>(N=5) | 4+5<br>(N=4) |  |  |  |  |  |
| Single layered epithe-<br>lium (TP1) | <u>52</u> | <u>7</u> | <u>65</u> | <u>6</u> | <u>2</u> | 132 | 52.2% | - | 132 | 20.4% |
| Small lumina (TP2) | <u>12</u> | <u>2</u> | <u>24</u> | <u>11</u> | <u>8</u> | 57 | 22.5% | - | 57 | 8.8% |
| High cellular density<br>(TP3) | <u>2</u> | <u>1</u> | <u>17</u> | <u>17</u> | <u>11</u> | 48 | 19.0% | - | 48 | 7.4% |
| Hyperchromatic nu-<br>clei with halo (TP4) | <u>1</u> | <u>0</u> | <u>6</u> | <u>1</u> | <u>8</u> | 16 | 6.3% | - | 16 | 2.5% |
| Small blood vessel w/<br>activated endothelia<br>(FP1) | 1 | 0 | 0 | 0 | 0 | 1 | - | 10 | 11 | 1.7% |
| Single layered epithe-<br>lium (FP2) | 5 | 0 | 0 | 1 | 0 | 6 | - | 29 | 35 | 5.4% |
| High cellular density<br>(FP3) | 5 | 3 | 0 | 2 | 0 | 10 | - | 8 | 18 | 2.8% |
| Small lumina (FP4) | 3 | 0 | 0 | 0 | 0 | 3 | - | 25 | 28 | 4.3% |
| Hyperchromatic nu-<br>clei with halo (FP5) | 0 | 1 | 2 | 0 | 0 | 3 | - | 12 | 15 | 2.3% |
| Two layered epithe-<br>lium (TN1) | 13 | 2 | 11 | 11 | 6 | 43 | - | 29 | 72 | 11.1% |
| Low cellular density<br>(TN2) | 23 | 6 | 4 | 2 | 2 | 37 | - | 125 | 162 | 25.0% |
| Highly polarised cells<br>(TN3) | 5 | 0 | 1 | 1 | 2 | 9 | - | 30 | 39 | 6.0% |
| Two layered epithe-<br>lium (FN1) | 0 | 0 | 0 | 1 | 0 | 1 | - | 0 | 1 | 0.2% |
| Undefined | 6 | 1 | 0 | 2 | 0 | 9 | - | 4 | 13 | 2.0% |

**Figure 5:**
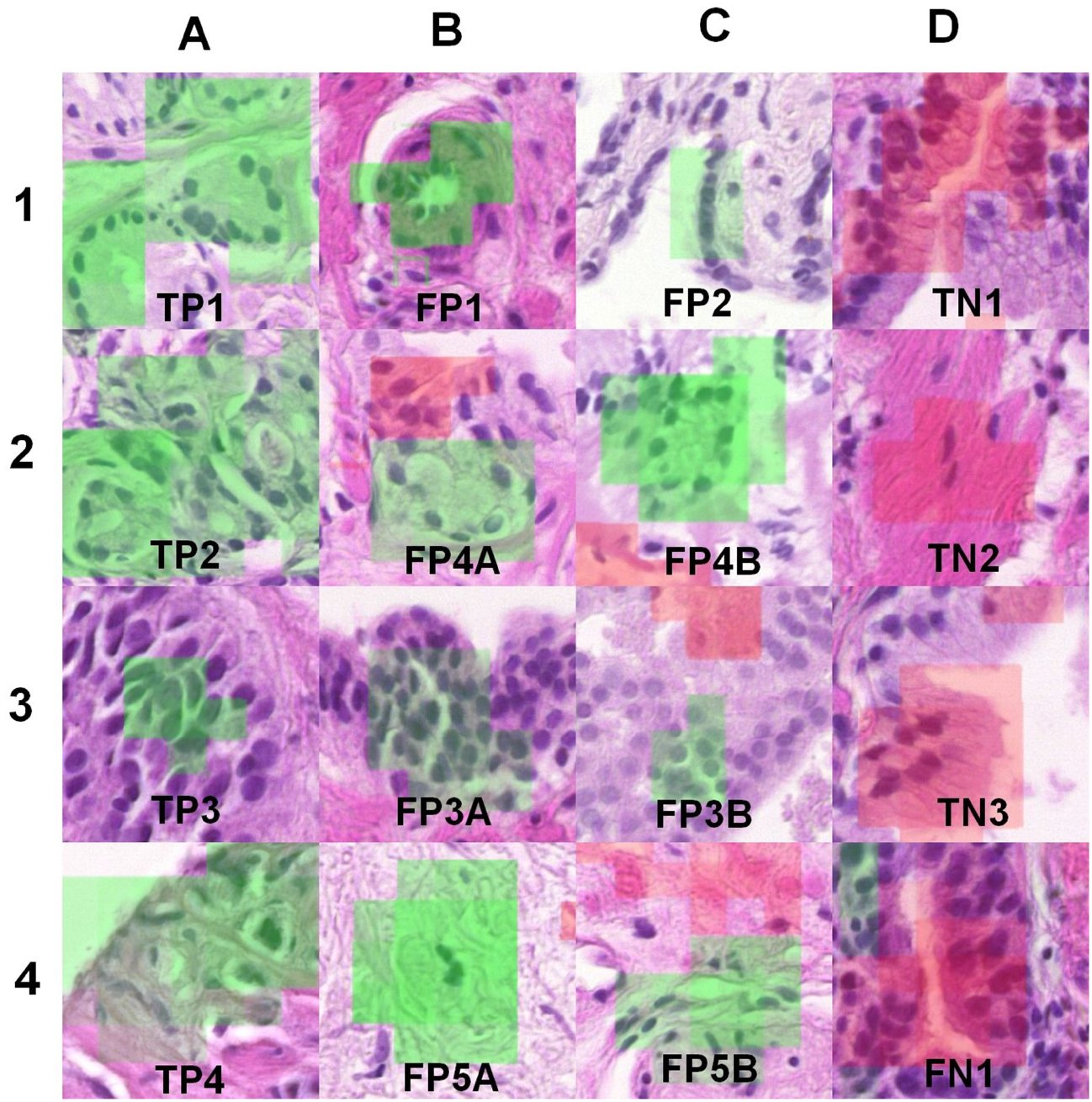
Explainability patterns overview. In column A, patterns supporting the classification of a tile as malignant (green overlay) in carcinoma cases are provided. These are single layered epithelium (TP1, fig. A1), small lumina (TP2, fig. A2), high cellular density (TP3, fig. A3) and hyperchromatic nuclei with halo (TP4, fig. A4). Columns B and C demonstrate the mimickers of these malignant patterns in non-malignant prostate tissue: small vessel (FP1, fig. B1), small lumina (FP4, figs B2, C2), single layered epithelium (FP2, fig. C1), high cellular density (FP3, figs B3, C3) and hyperchromatic nuclei with halo (FP5, figs B4, C4). Column D shows typical patterns acting against classification of a tile as malignant (red overlay). One is two layered epithelium demonstrated in normal epithelium (TN1, fig. D1) or in carcinoma (FN1, fig. D4). The other two predominant benign patterns are low cellular density (TN2, fig. D2) and highly polarized epithelium (TN3, fig. D3). Additional examples of these patterns in a wider tissue context can be found in Supplementary Figure S2.2.

Patterns supporting cancer positivity were single layered epithelium (TP1), small lumina (TP2), high cellular density (TP3) and hyperchromatic nuclei with halo (TP4), while the patterns supporting a decision of cancer negativity were two layered epithelium (TN1), low cellular density (TN2) and highly polarized cells (TN3). As these patterns may not be very informative in the sense of explainability (they can simply reflect the most prevalent pattern in cancerous or non-cancerous tissue), the overlays were checked also for false positive and false negative results.

Five typical patterns for false positivity and one for false negativity were found (Table 2, Figure 5, Supplementary Figure S2.2). False positivity patterns reflected the patterns of true positivity: small blood vessels with activated endothelium (FP1) probably include two other false positive patterns of single layered epithelium (FP2) and small lumina (FP4). The other false positive patterns are high cellular density in tangential sections of non-cancerous epithelium (FP3) and larger hyperchromatic stromal nuclei with halo (FP5). The single false negative pattern was two layered epithelium (FN1).

The results of evaluation are presented in Table 2. Overall, of 647 sample points, 253 (39.1%) were evaluated as true positive, 107 (16.5%) as false positive, 273 (42.2%) as true negative, and 1 (0.2%) as false negative. The 13 (2%) samples where it was not clear what information was highlighted by the explanation map were labeled as undefined. Single layered epithelium represents the predominant true positive pattern (20.4% of total, 52.2% of true positive sample points), followed by small lumina (22.5% of true positive samples) and high cellular density (19% of true positive samples).

Note the change in pattern distribution related to the Gleason score in carcinoma tissue (Table 2, underlined, in italics), showing a shift from single layered epithelium and small lumina, predominant in pattern 3, to high cellular density and hyperchromatic nuclei with halo, predominant in patterns 4 and 5, which represents internal controls of reading validity, but unfortunately provides limited explanation, following the predominant tissue morphology. Among the false positive patterns, there is no obvious predominance, but due to the similarity to true positives they provide much more information regarding explainability. In the true negatives, low cellular density, in most cases related to the fibromuscular stromal component, predominates (25% of total sample points), followed by two layered epithelium (11.1% of total sample points), again providing less explanation. The false negative samples are too rare to provide any reasonable explainability information.

## 4 Discussion

The histopathological diagnosis of prostate carcinoma is based on a relatively consistent set of morphological features. At low magnification (20×–50×) this is mainly the abnormal density and shape of glands. Cancers of Gleason patterns 4 (predominantly areas of fused glands with high epithelial density) and 5 (complete loss of glandular pattern with dissociated or medullary growth, sometimes cells with hyperchromatic nuclei and vacuoles) can be usually diagnosed at this resolution level. Pattern 3 (tubular growth) can be diagnosed at lower magnification if infiltration is extensive enough to noticeably change gland architecture. In case of doubt, suspect areas can be checked at medium magnification (10×–20×) for the absence of basal cells (i.e. the presence of single layered epithelium), which represents another hallmark of prostate cancer. This is especially useful to recognize the small foci of pattern 3 carcinomas. High magnification (40×–60×) can reveal additional cell details, namely the presence of nucleoli that confirm the diagnosis of carcinoma in some cases.

Our results demonstrate that a neural network trained to detect prostate carcinoma in core biopsies uses similar morphological characteristics to pathologists at low and medium optical magnification. The neural network applies the same features (especially the presence of small glands (typical for patterns 3 and 4), single layered epithelium (typical in pattern 3), high cellular density (characteristic for most cases in pattern 4 and 5), and hyperchromatic nuclei with halo (often seen in pattern 4 and 5)) to label areas suspicious of being cancerous. Consistent with the available resolution, the features of nuclear detail (e.g., presence of nucleoli, chromatin structure) were not used, as they are not clearly visible in the scans.

Compared to an experienced pathologist, our neural network was less effective when considering the tissue context, such as gland density and shape. This may be attributable to the tiling scale used for training and analysis and should be further explored. Typical failures are represented by labeling of small vessels with prominent endothelium as suspect due to similarity to small tubules of pattern 3 or 4 carcinoma, by poor interpretation of tangential sections in non-cancerous epithelium, resulting in false recognition of high cellular density, typical for patterns 4 and 5 carcinoma, or in labeling neural tissue and some stromal cells because of larger hyperchromatic nuclei with halo, confused with atypical cells found in pattern 4 and 5 cancers. Not surprisingly in the above sense, admixed colon mucosa was labeled as cancer (Supplementary Figure S2.3). On the other hand, the most frequent mimickers of Gleason pattern 3 cancer in core biopsies (i.e., atrophy, partial atrophy and post-atrophic hyperplasia) were recognized as suspect with only low probability, similar to a pathologist.

It is important to note that our consecutive datasets did not contain rare carcinoma types, e.g. foamy cells or pseudohyperplastic tumors, and inflammatory lesions were under-represented. However, the main task of this study was not to develop a screening tool, but to identify a method for identifying, visualizing, and analyzing the specific features that are important for the decision making of neural networks—to pierce its black box. It was an important finding that the analysis of tissue patterns selected by occlusion maps according to their contribution to the classification requires not only truly positive, but also negative tissues containing morphological mimickers. This should be considered when planning similar studies. The occlusion-based explanations pointing to critical structures represent a useful tool for fine-tuning and optimization of neural networks in histopathology, and potentially for identification of previously unrecognized morphological features related to histopathological diagnosis, prognosis, and prediction [23, 24]. Finally, unravelling the large quantity of features within the network and exposing the key elements will help to promote trust in these and similar AI-based methods in pathology, enhancing the opportunities for incorporation into clinical use.

## Acknowledgements

Computational resources were supplied by the project “e-Infrastruktura CZ” (e-INFRA CZ LM2018140) supported by the Ministry of Education, Youth and Sports of the Czech Republic. The authors wish to thank to Philip Coates for revision and editing of the manuscript.

## Ethics Approval

The project was approved by the Ethical Committee of Masaryk Memorial Cancer Institute, No. MOU 385 920.

## Footnotes

1 https://computationalpathologygroup.github.io/ASAP/

2 https://openslide.org/formats/mirax/

3 https://www.bbmri-eric.eu/services/access-policies

4 https://negotiator.bbmri-eric.eu/

