## Supplementary Information for "Shedding Light on the Black Box of a Neural Network Used to Detect Prostate Cancer in Whole Slide Images by Occlusion-Based Explainability"

### S1 Deep learning details

#### S1.1 Data Preprocessing

The train and test split was performed on a patient level, whilst train and validation split was performed on a slide level. We ensured that every split contains at least one positive slide, and one negative slide.

| | | Number of cases | |
|---|---|---|---|
| ISUP grade | Gleason | Training | Test |
| 1 | 3+3 | 45 | 5 |
| 2 | 3+4 | 33 | 1 |
| 3 | 4+3 | 49 | 1 |
| 4 | 4+4 | 11 | 1 |
| 5 | 4+5 | 14 | 2 |
| Total with cancer | | 122 | 10 |
| Negative | | 51 | 0 |

| | Number of slides | |
|---|---|---|
| | Training | Test |
| WSI with cancer | 322 | 37 |
| WSI without cancer | 539 | 50 |

**Table S1.1:** Distribution of cases in the training and test sets according to grade.

Due to their large size (105185×221772 px), the slides were cut into smaller patches of size 512×512 px. To conserve memory and speed up the training process these patches were not stored on the disk. Instead, they were sampled directly from the slides on-the-fly during both the training and testing using the *openslide-python* package.

**Slide Preparation** First, we restricted the sampling process only to tissue area. To do this, every slide was downscaled 16×, converted to HSV and Otsu's thresholding was applied on the saturation channel [1–3]. A morphological closing followed by morphological opening was applied on the thresholded saturation channel with a disk of size 10. The result is a binary background mask with only the tissue area being highlighted.

To assign a label to each patch we created another binary mask using the annotations provided by pathologists. These annotations were created in the ASAP [4] annotation tool and were stored in an XML file. The XML file described polygons by listing the $x$ and $y$ coordinates for each vertex of an annotation polygon. We created the binary mask by drawing the polygons described in the accompanying XML file. The resulting masks are downscaled by a factor of 2.

Next, we applied a sliding window with a step size 256 px on the background mask. All patches whose tissue coverage did not exceed a threshold of 75% were ignored. To assign labels to the remaining patches, we used the annotation binary mask. A patch was assigned a positive label if its center area of size 256 px contained at least one pixel belonging to the cancer category [1, 5]. For each slide we then saved the extracted patches in a table describing only the $x$ and $y$ coordinates of the top-left pixel of the patch and the label. No table exceeded 100 KB in size.

**Input Preprocessing** The dataset is highly imbalanced. The training set contained over 8 million negative patches and over 1.5 million positive patches. To combat this imbalance, we applied image augmentation techniques to artificially enlarge the dataset. We used the following augmentations: random vertical and horizontal flips with 50% probability, random brightness perturbations in range $[-64, 64]$, random hue perturbations in range [−10,10], random saturation perturbations in range [−64,64] and random contrast perturbations in range [0.7,1.3] [1, 2, 5]. We used the *imgaug* package to augment the patches. Finally, all patches were scaled to $[-1, 1]$ range.

**Patch Sampling** During the training a three step sampling process is used to deal with the imbalanced dataset [5]. In the first step a label $L$ is selected uniformly at random. In the next step a single slide $S$ is picked uniformly at random from all the slides containing at least one patch with label $L$. In the last step a patch with label $L$ is selected uniformly at random from the slide $S$. This procedure also ensures that patches will be drawn from all slides equally regardless of how much tissue area or cancer area is present in the slide.

#### S1.2 Network Architecture

We used a VGG16 network pretrained on imagenet dataset for feature extraction. The model was trained using an RMSprop optimizer with the following parameters: momentum=0.9, epsilon=1.0, rho=0.9, initial learning rate of $5 \times 10^{-5}$ and batch size of 1. Learning rate was halved after every 5 consecutive epochs without improvement on a validation data. The training would stop if no improvement on a validation data is made for 10 consecutive epochs.

#### S1.3 Explainability methods

Our choice of explanatory methods is driven by two fundamental principles: simplicity and fidelity. We prefer methods that are both simple to implement and easily clarified to a human user. By fidelity, we mean an explanatory method should be as accurate as possible to the behaviour of the model.

There is some concern in the explainable artificial intelligence community that, based on human evaluation of explanations alone, researchers might tend to be biased towards creating systems which are persuasive, rather than accurate, in their explanations. Indeed, recent work has shown that some popular saliency visualization techniques are independent of both model parameters and the training data [6].

We assess three well-established explanatory techniques: gradient-based saliency mapping [7], LIME [8] and occlusion sensitivity [9]. Rather than trying to explain the behaviour of the model as a whole in the global sense, all three methods aim at providing explanations which are local to a particular input, to hopefully reinforce trust in the decision.

**Gradient-based Saliency** In gradient-based saliency mapping, a salience mask is produced by computing the gradient $E$ of the class score $S_c$ of the class of interest $c$ with respect to the input $x$, i.e. $E(x) = \frac{\partial S_c}{\partial x}$. We can then obtain a 2D salience mask by taking for each pixel the largest absolute gradient value across color channels. Intuitively, the resulting mask signifies those pixels of the input $x$ in which a tiny value change results in the largest change in the prediction $S_c(x)$. The method passes the sanity checks presented in [6]. Gradient-based saliency additionally has the advantage of being very fast to compute, as it requires only a single weight backpropagation pass through the network.

One major disadvantage of gradient-based saliency is its inability to show the kind (positive or negative) of contribution of a pixel to the final network prediction (see Figure S1.1).

**LIME** LIME is a conceptual framework to generate explanations of model behavior by training a surrogate linear model which assigns weights to interpretable units of input information. The training

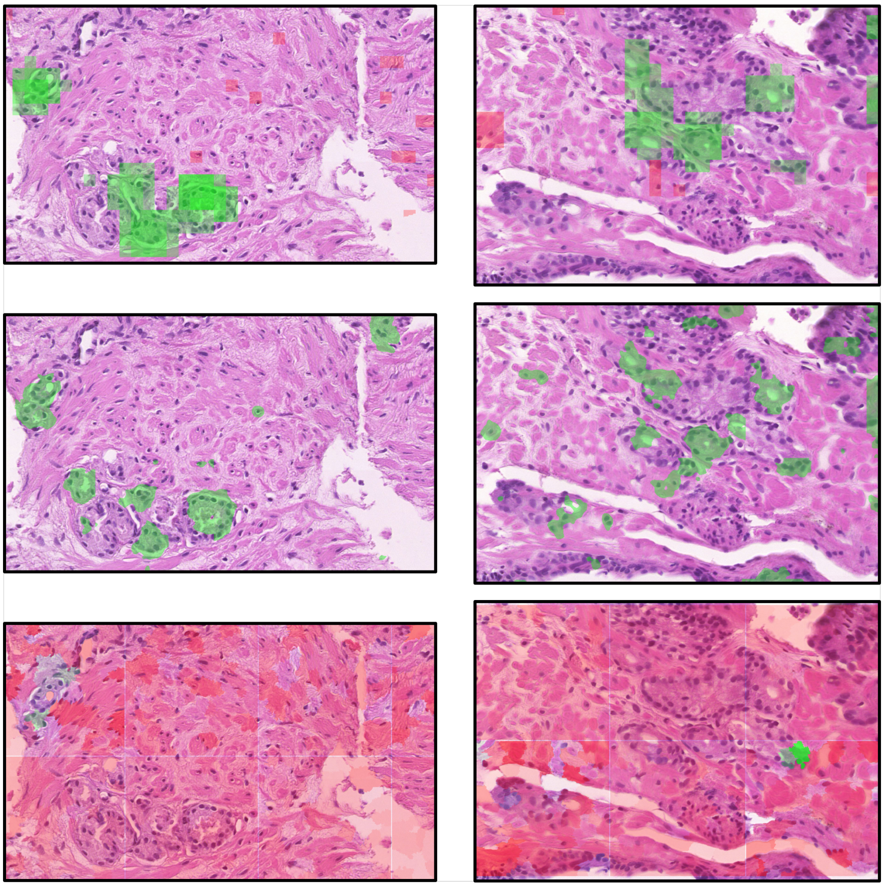

**Figure S1.1: Copmarison of tested methods.** Section **a)** occlusion analysis (top) and gradient-based saliency (bottom) – note that regions highlighted in a saliency map may in fact be contributing negatively to a cancer prediction, as seen in the occlusion map; Section **b)** occlusion analysis (top), gradient-based saliency (middle), and LIME (bottom) – note that LIME positively highlights only a minor portion of the tissue, while the occlusion and saliency maps correspond well in highlighting significant areas.

is done by randomly removing the chosen units of input information from an input, and treating the main model as an oracle, which produces ground-truth labels.

When applied to image data, the interpretable units of input are particular portions (superpixels) of the image. In this context, LIME first applies a segmentation algorithm to the input image to obtain the superpixels, which are then repeatedly randomly covered and the perturbed input obtained in this way is passed in to the network. At the same time, a linear model with the same count of features as there are superpixels in the input segmentation is trained, such that each feature of the model corresponds to a particular superpixel. When a superpixel is covered in the input of the original network, its corresponding feature in the input of the linear model is set to 0, while all features corresponding to the unperturbed supexpixels are set to 1, and the network's prediction on the perturbed input image is used as the label for the linear model.

One problem with this approach is its dependence on a well-chosen segmentation algorithm and its parameters. As segmentation algorithm performance on a given input may vary even for small changes in its parameters, this approach is unstable. Furthermore, in our experiments, LIME wasn't able to capture most of the areas positively contributing to a cancer prediction in the whole slide image (see S1.1).

**Occlusion Sensitivity** An even simpler technique is occlusion sensitivity analysis, where a salience mask is created by systematically perturbing the input and observing the corresponding changes in the model's output. Specifically, we systematically cover (occlude) square patches of the input by setting their values to zero, and record the output change in a matrix, whose value positions correspond to the relative position of the covering patch in the input. This matrix can then be upsampled to obtain a mask covering the whole original input.

One major disadvantage of occlusion sensitivity analysis is the need to systematically create large quantities of perturbed inputs and record the change in network response for each one. Based on the occlusion parameters (the size of the occlusion window and stride of each step), the number of perturbed images needed to produce an occlusion heatmap for a single input will vary, with finer-grained maps necessitating much more operations than low resolution maps. In terms of asymptotic notation, the number of perturbed inputs necessary to produce an occlusion sensitivity map is in $O(w \times h)$, where $w$ and $h$ are the width and height of the original input image, respectively. Compared to gradient-based saliency, which needs a single backpropagation pass to generate its explanation map, occlusion sensitivity analysis is a significantly more time consuming process.

Our environment presents several idiosyncrasies which force us to further manipulate the obtained masks. Typically, the results of each method are normalized to a desired range to comprehensibly indicate those regions with the highest or lowest contribution to a particular decision. This is usually not a problem, as decisions are considered independently on one another.

Both our classification results and their explanations are presented as heatmaps covering the entirety of a WSI image, which, as described in S1.1, consists of thousands of individual inputs to the model. We cannot therefore treat the explanations for inputs obtained from a single whole slide image as being independent, but rather as part of one aggregate.

The challenge we face with occlusion sensitivity is appropriately expressing the same magnitude of increase/decrease in the model's response in different inputs in the context of the whole slide. Consider two inputs, $x$ and $y$, and their perturbations $x'$ and $y'$. Let us denote $\delta_x = S_c(x) - S_c(x')$ the difference between class scores for the original and perturbed input $x$, and analogously $\delta_y = S_c(y) - S_c(y')$. Now, suppose $\delta_x = \delta_y$, with $\delta_x, \delta_y > 0$, but $S_c(x) \gg S_c(y)$. Intuitively, in our binary classification scenario, even though the increase is the same for $x$ and $y$, the relative importance of the occluded regions in $x'$ and $y'$ should surely not be treated as equal in the global context. The method of visualizing occlusion maps described in the main paper adresses this problem adequately.

The modifications to the saliency-based maps are simpler. We are not concerned about the relative importance of the marked regions. Instead, as the heatmaps can be — somewhat imprecisely — interpreted as signifying those regions which "excite" the network the most, we are interested in their values as-is. However, since the distribution of the gradient values skews heavily towards 0, we employ a non-linear scaling function to spread the values out for easier visualization.

#### S2 Additional Explainability Examples

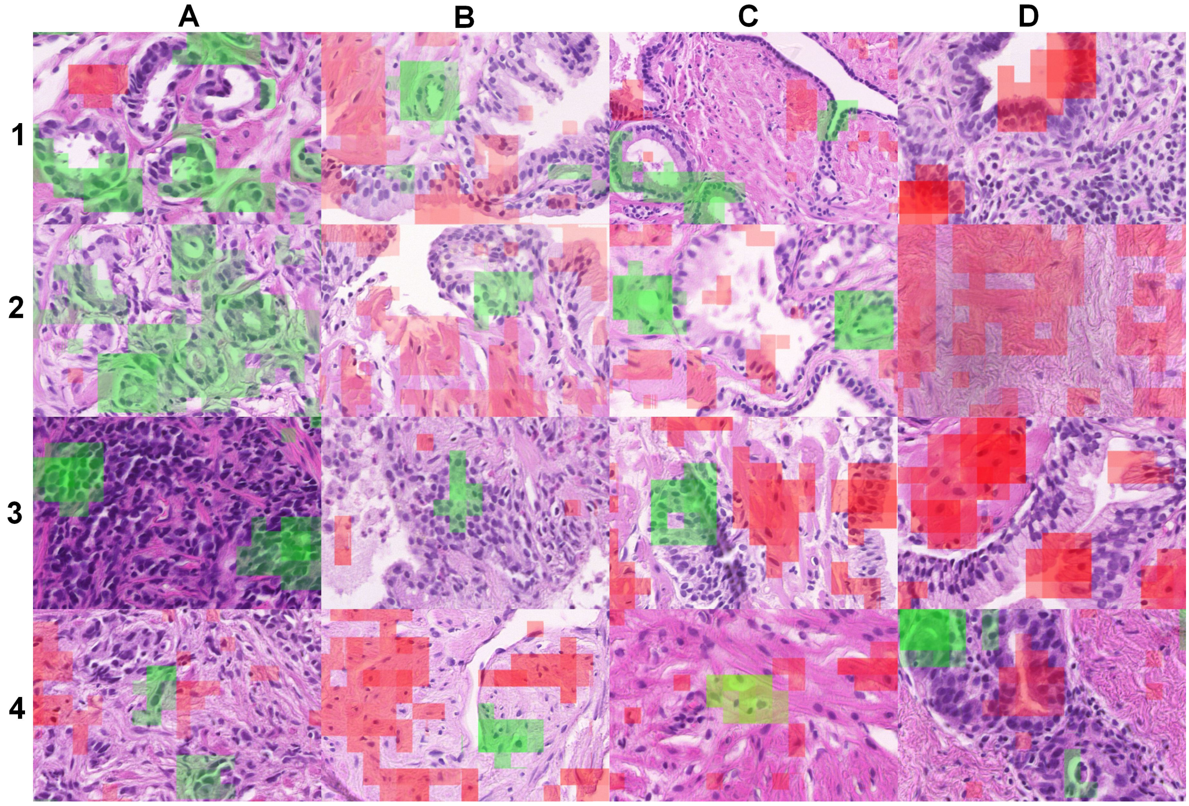

**Figure S2.2: Examples of typical explainability patterns.** Areas supporting the classification of a tile as cancer are highlighted in green, areas supporting the classification of a tile as benign in red. In column A are presented the typical true positive patterns in cancer samples: single layered epithelium (A1), small lumina (A2), high cellular density (A3) and hyperchromatic nuclei with halo (A4). Columns B and C show examples of their mimickers in non-cancerous samples. In column D are given the typical true negative patterns: Two layered epithelium (D1), low cellular density (D2) and highly polarised epithelium (D3) in noncancerous samples. In D4, a false negative pattern is shown: two layered epithelium in a cancerous sample.

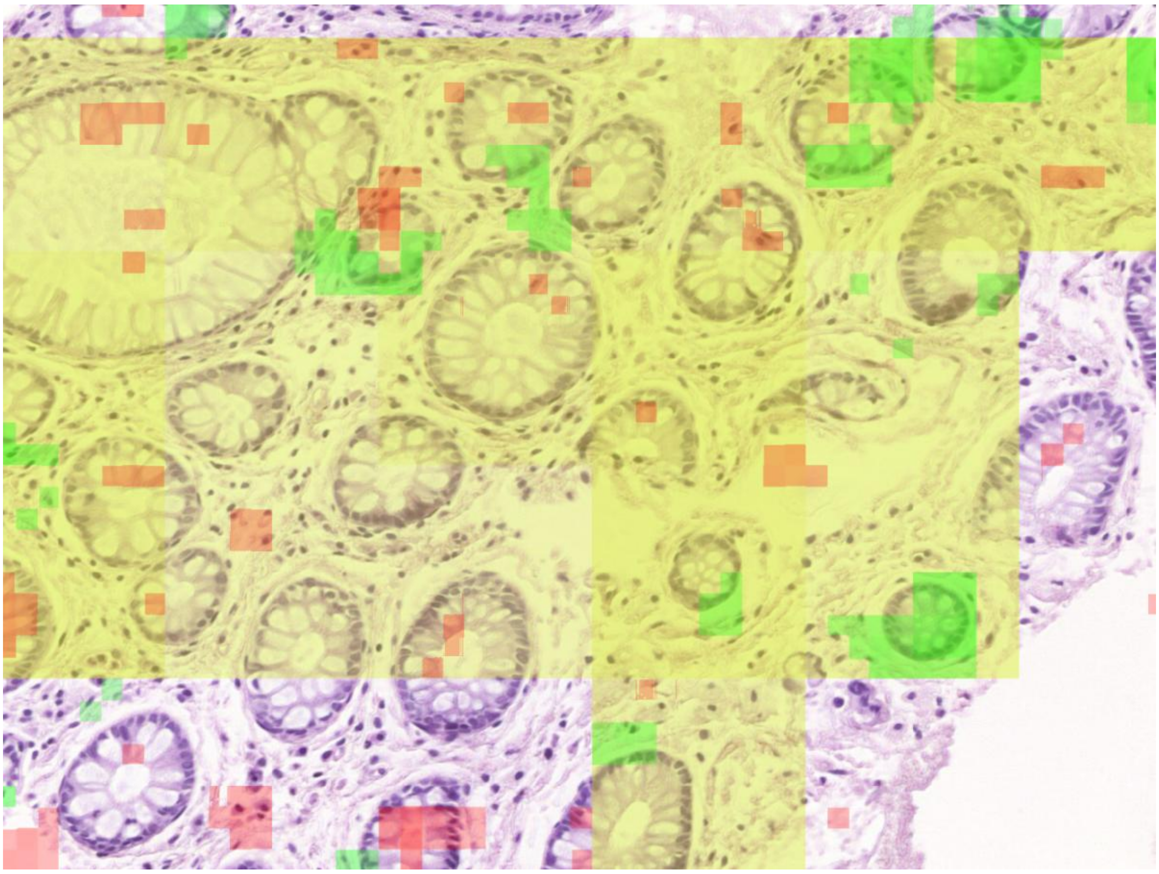

**Figure S2.3: Colon mucosa admixed to prostate sample evaluated by network.** The greater part of sample is evaluated as suspect of carcinoma (yellow overlay) due to the presence of single layer epithelium (green explainability overlay). The intervening stroma and highly polarised cells are evaluated as non-suspect (red explainability overlay).

#### S3 Overview of test WSI sections

An overview of the data used for testing follows. Note that some examples may exhibit a visual artifact in the form of a green or yellow line at the bottom, these are only an artifact of the overview generation, and are not present in the WSI overlay.

##### S3.1 Positive WSI

The following cases come from WSI which contain carcinoma. From left to right, the images presented contain: a portion of the original WSI; the same region overlaid by probability of cancer masks (yellow) and manual ground-truth annotations (blue); the same region overlaid by probability of cancer masks (yellow) and occlusion-based explanation masks (green and red, signifying positive and negative contribution towards cancer diagnosis, respectively).

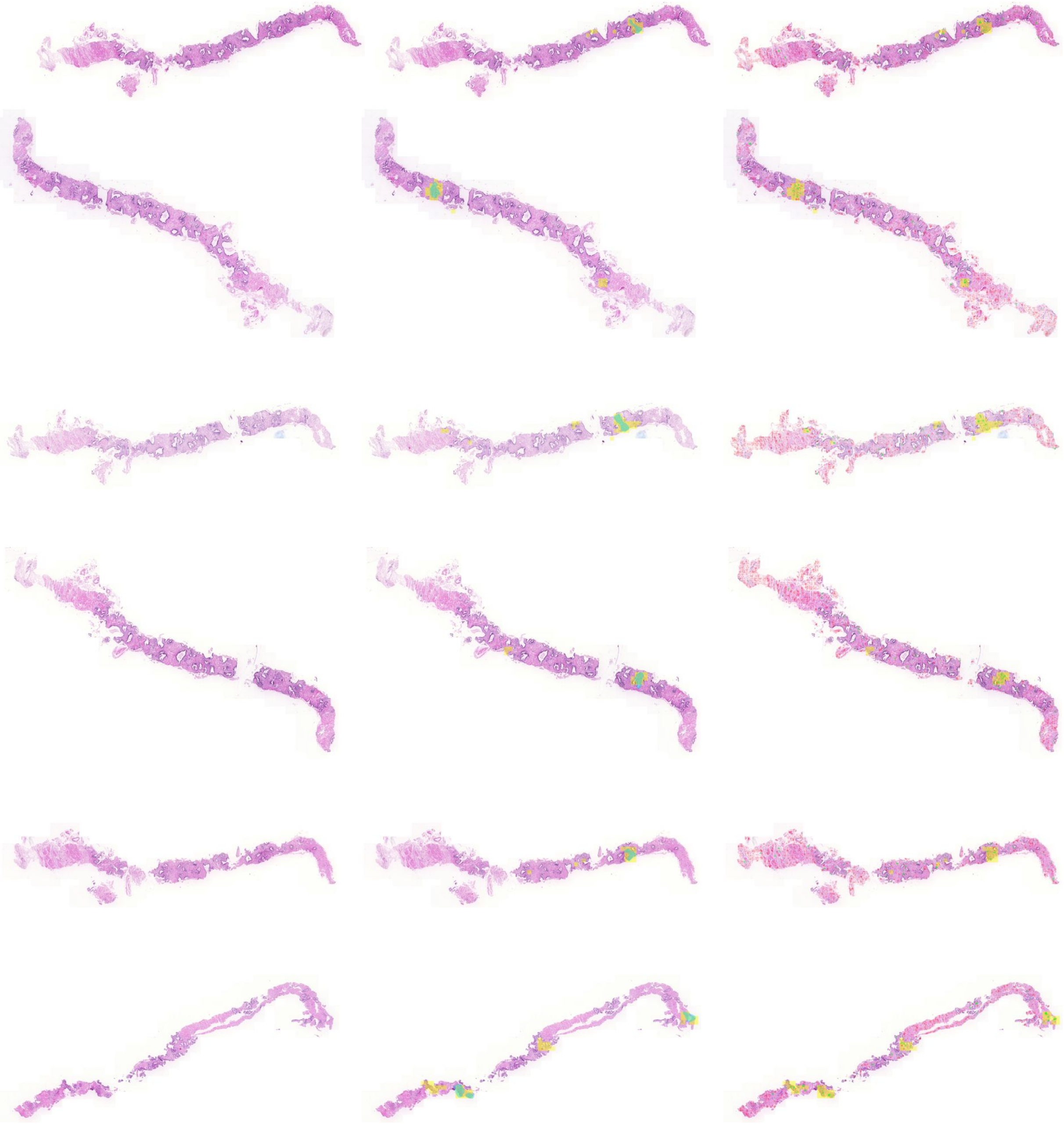

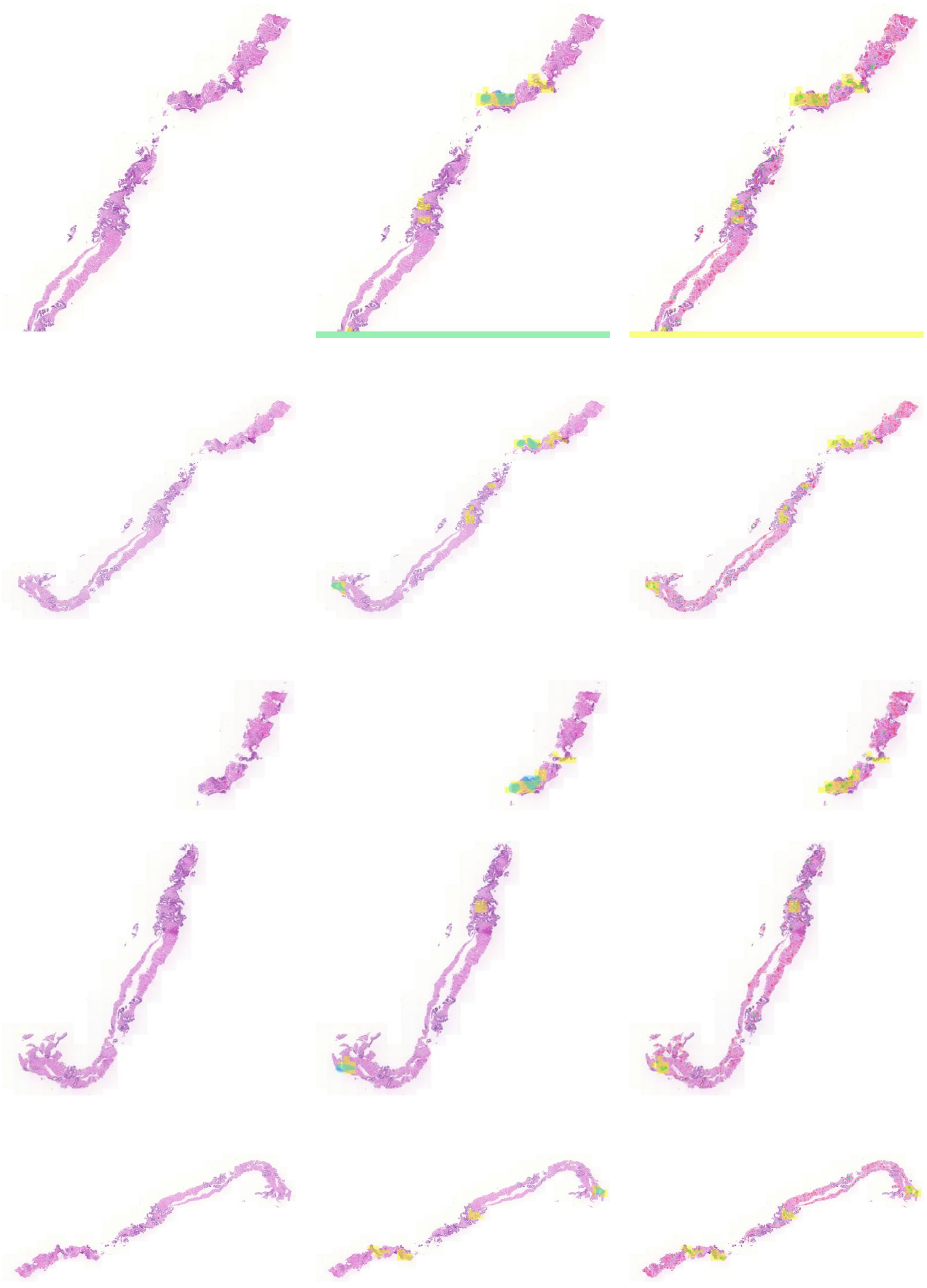

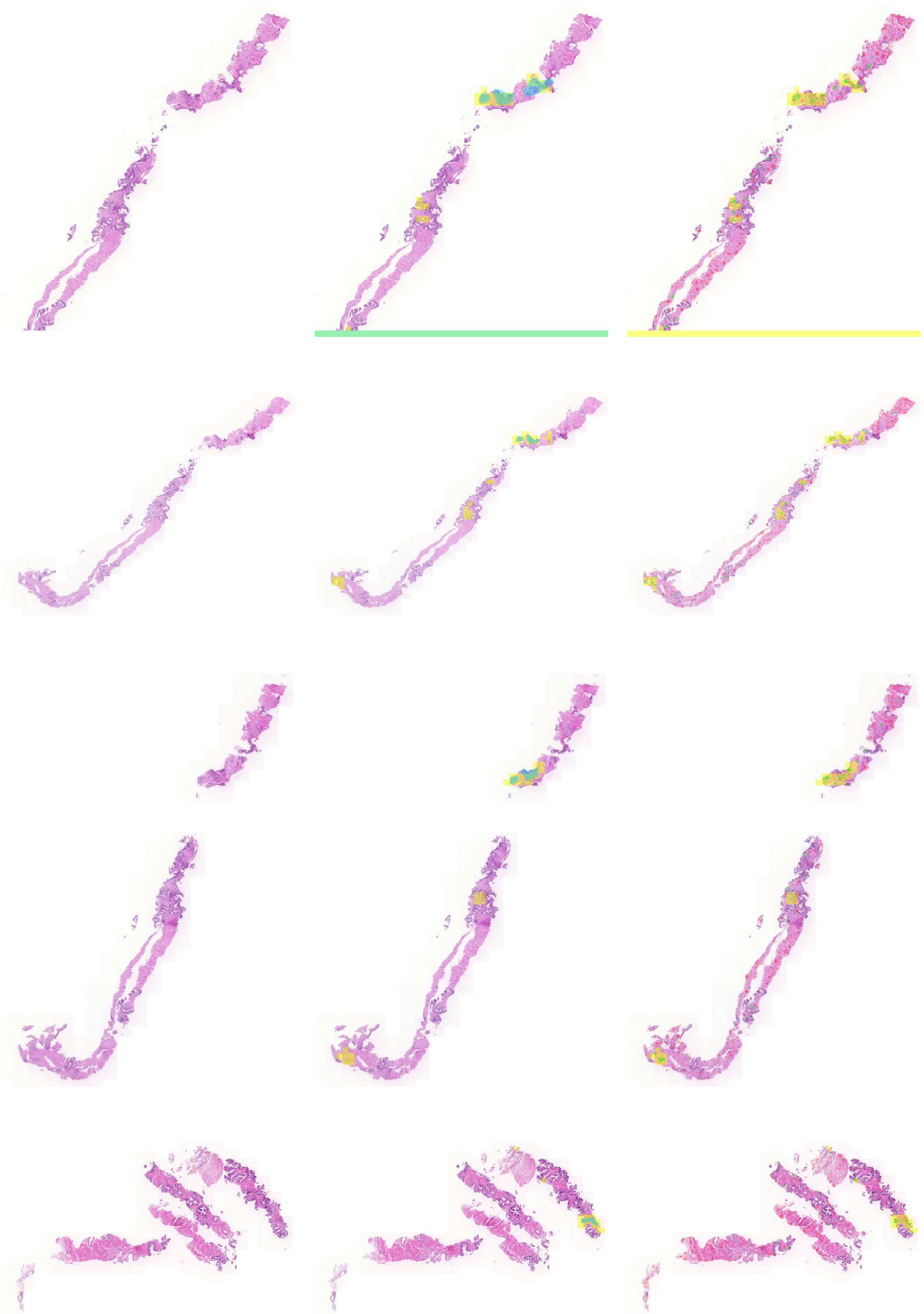

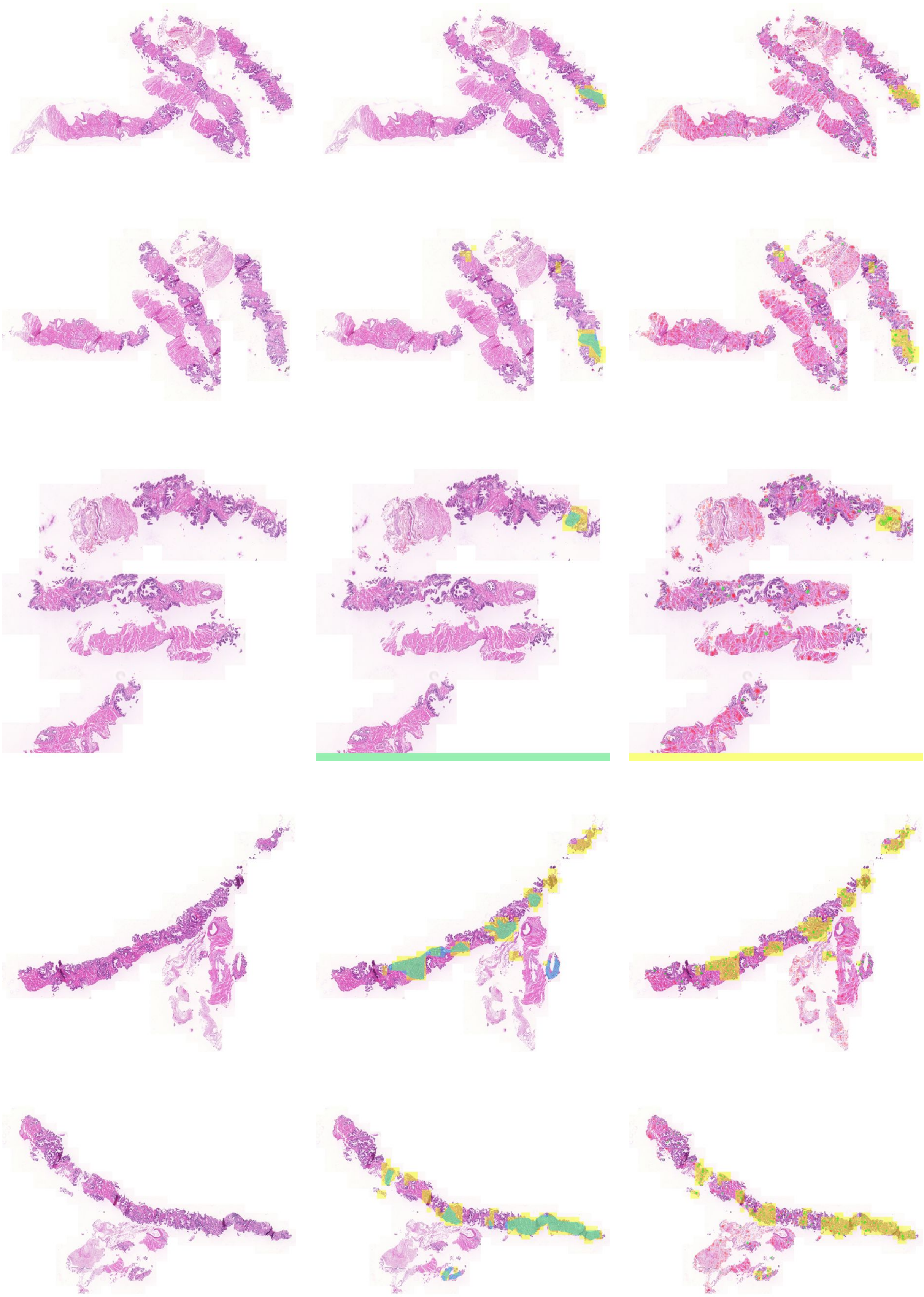

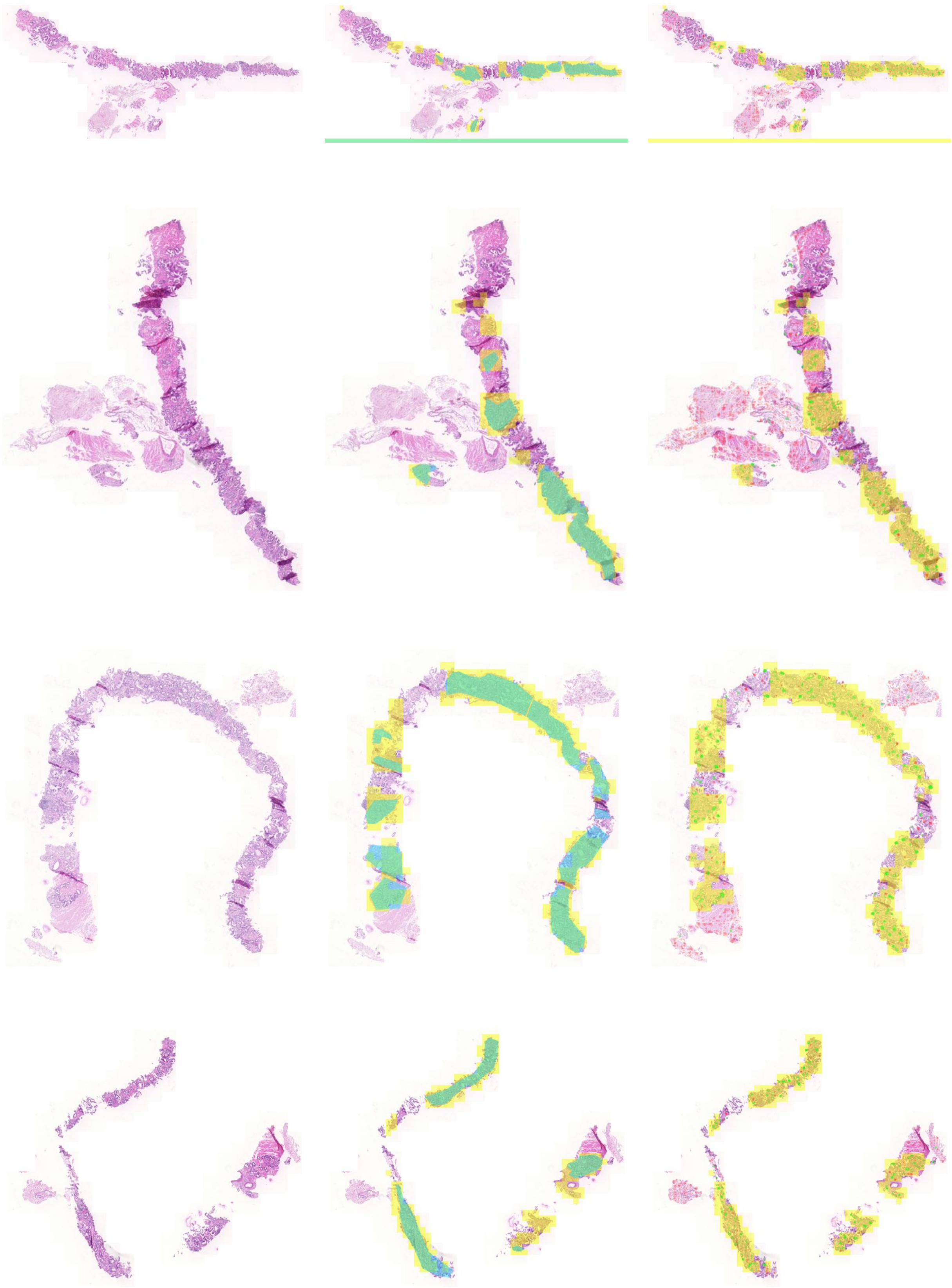

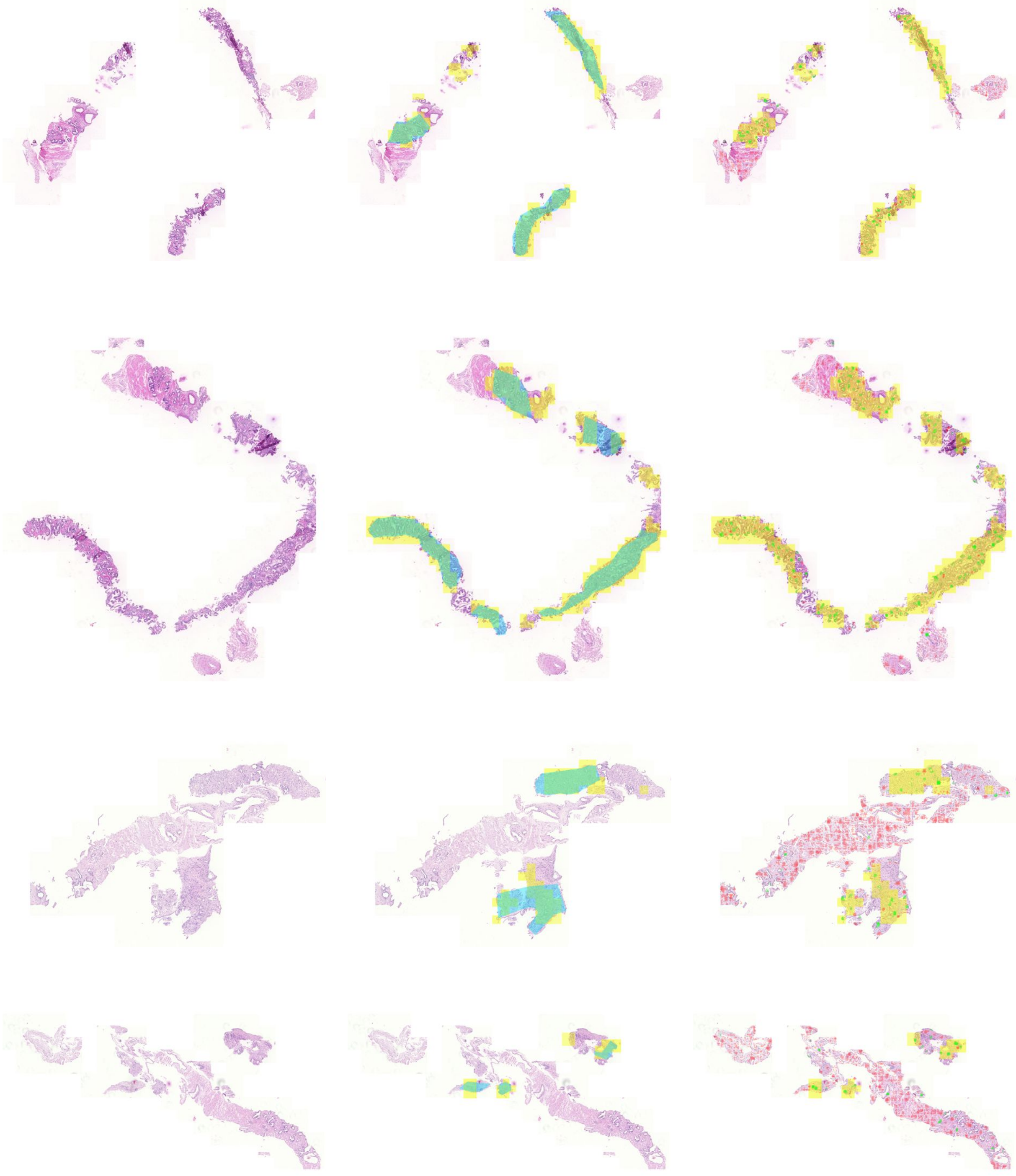

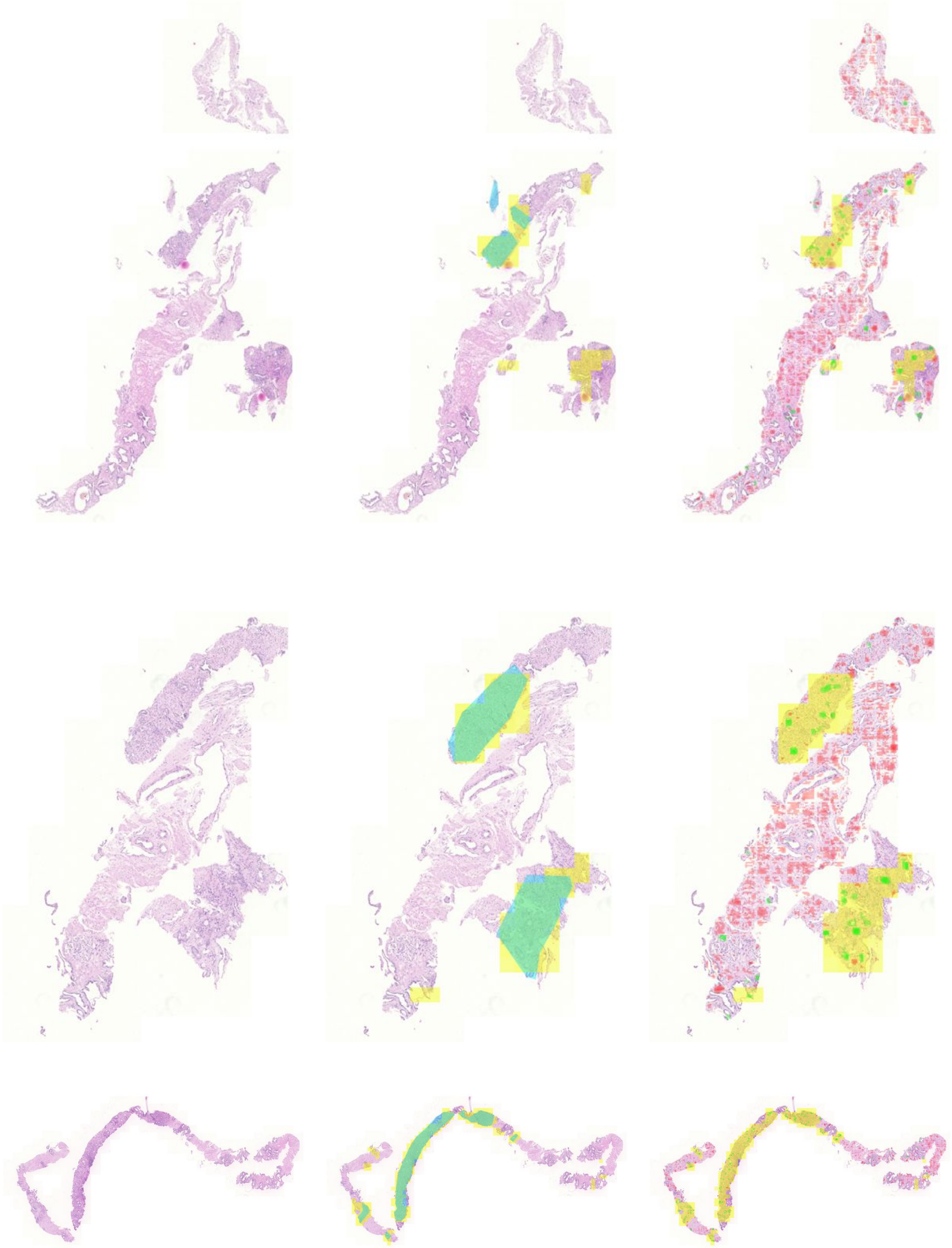

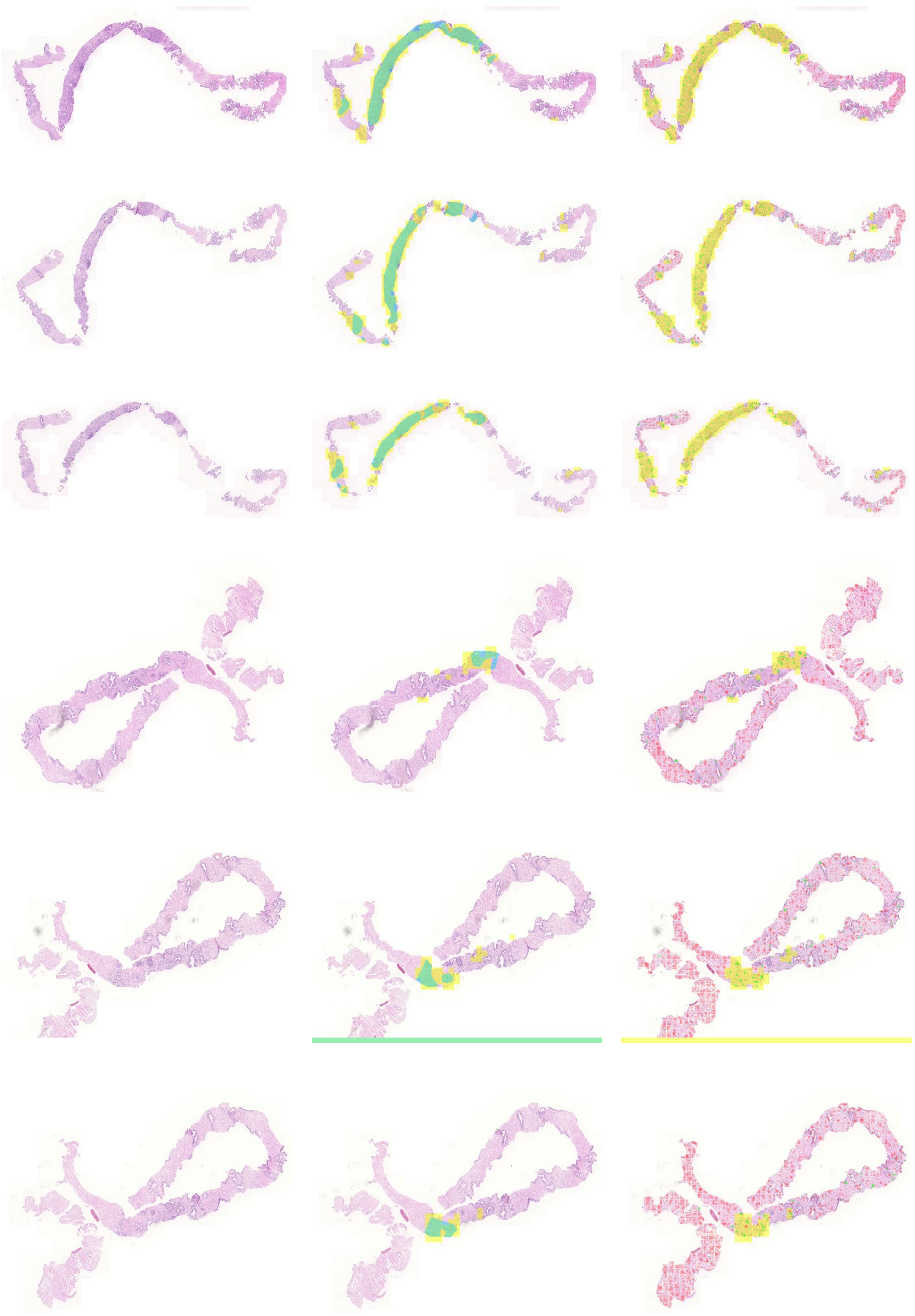

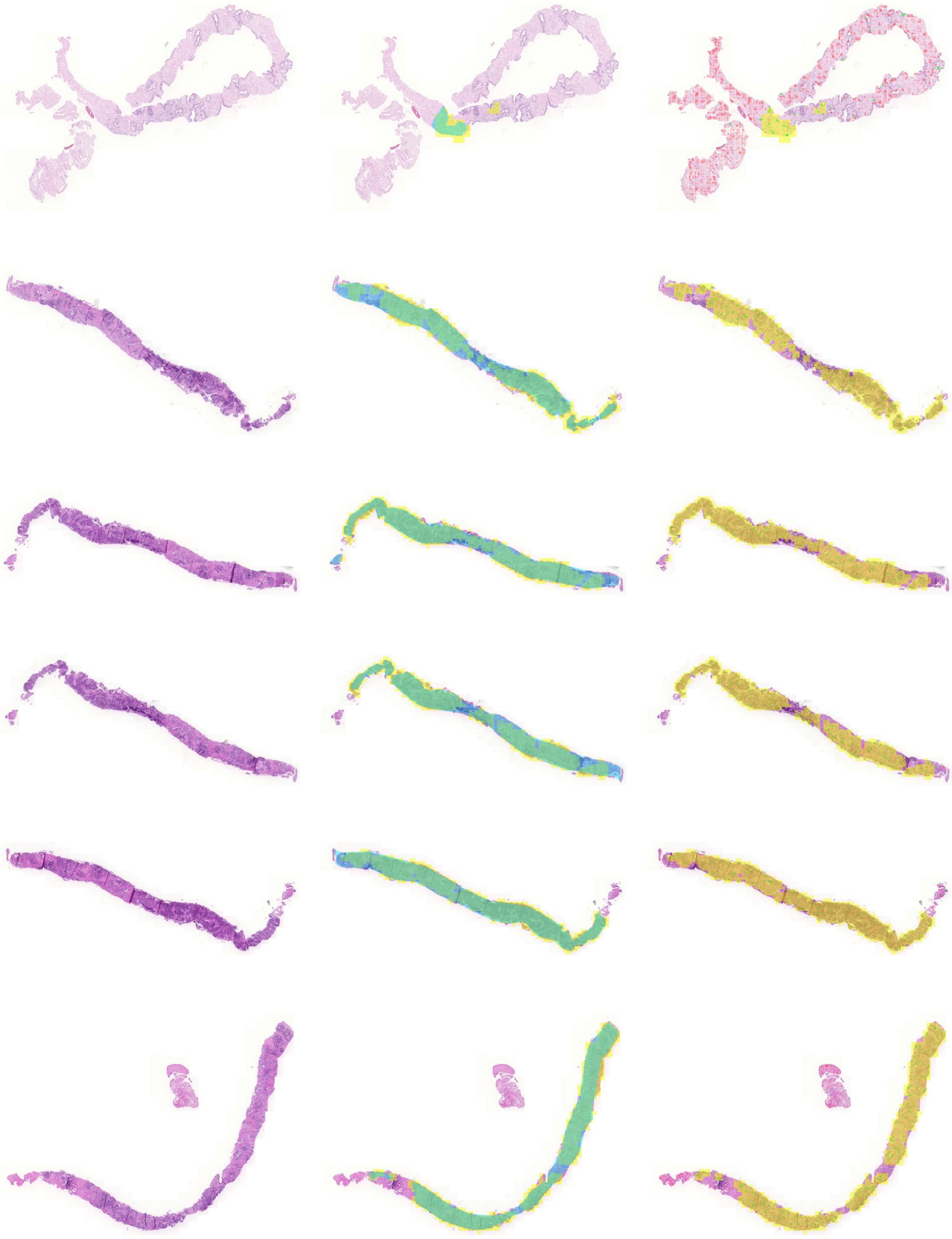

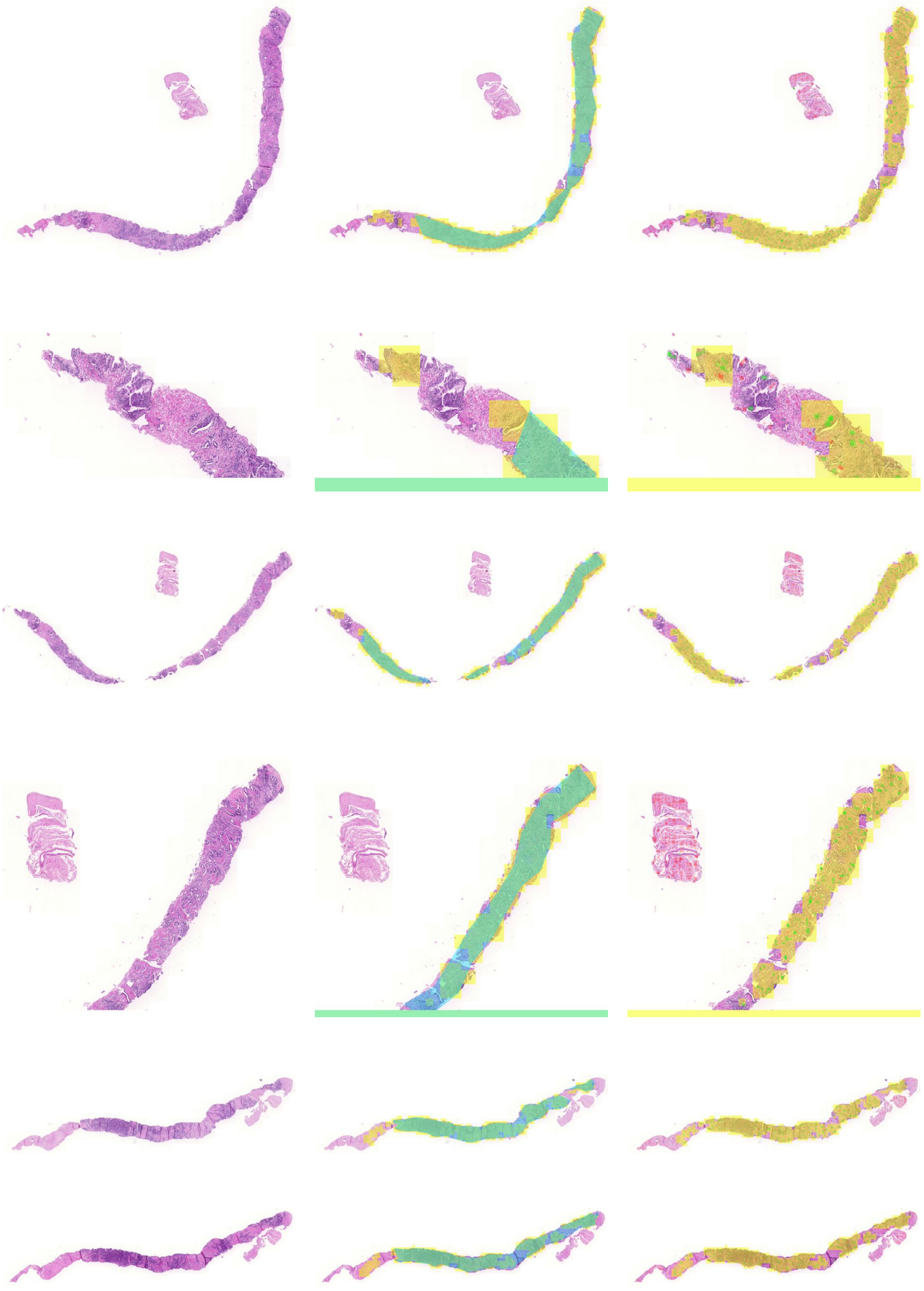

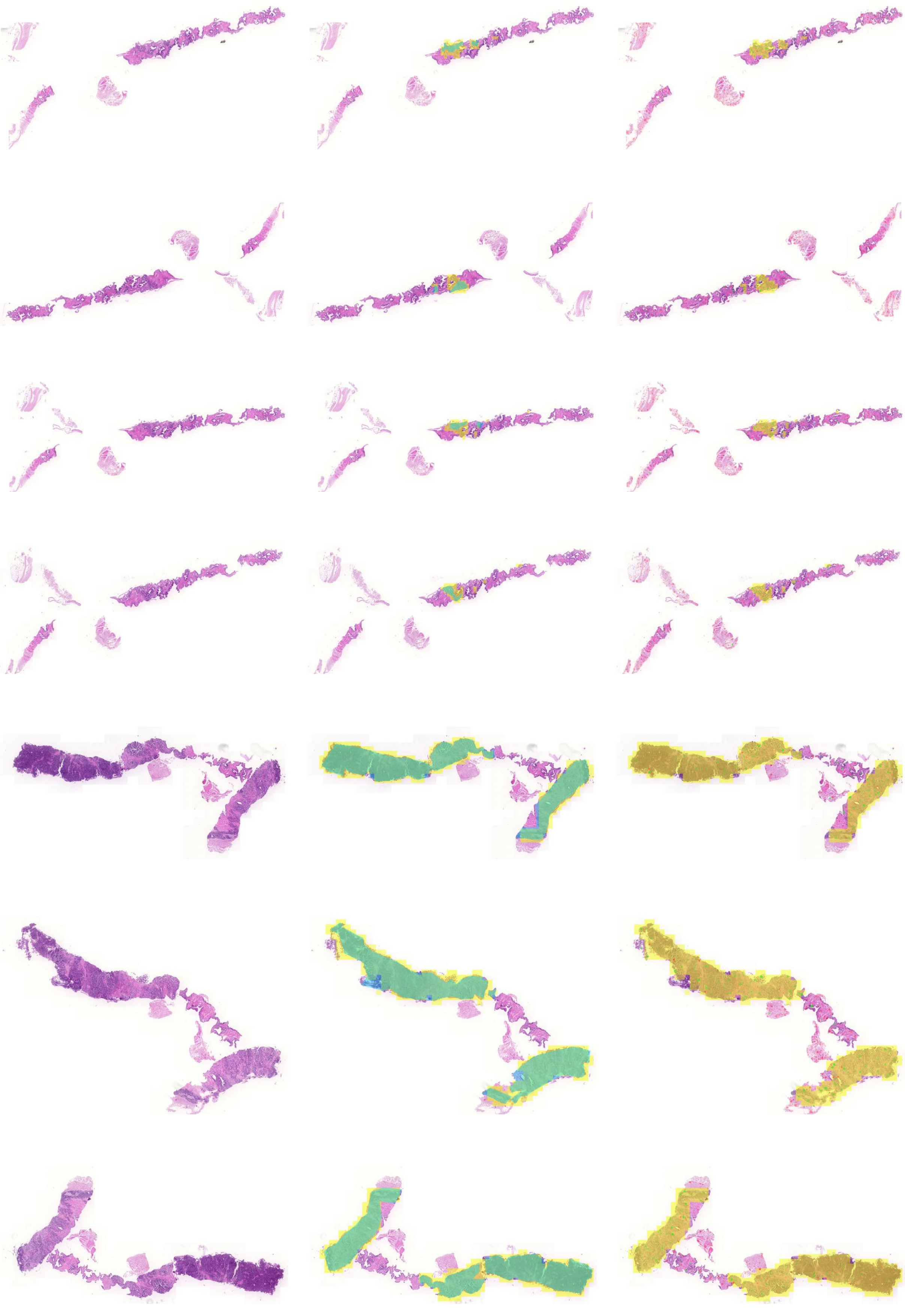

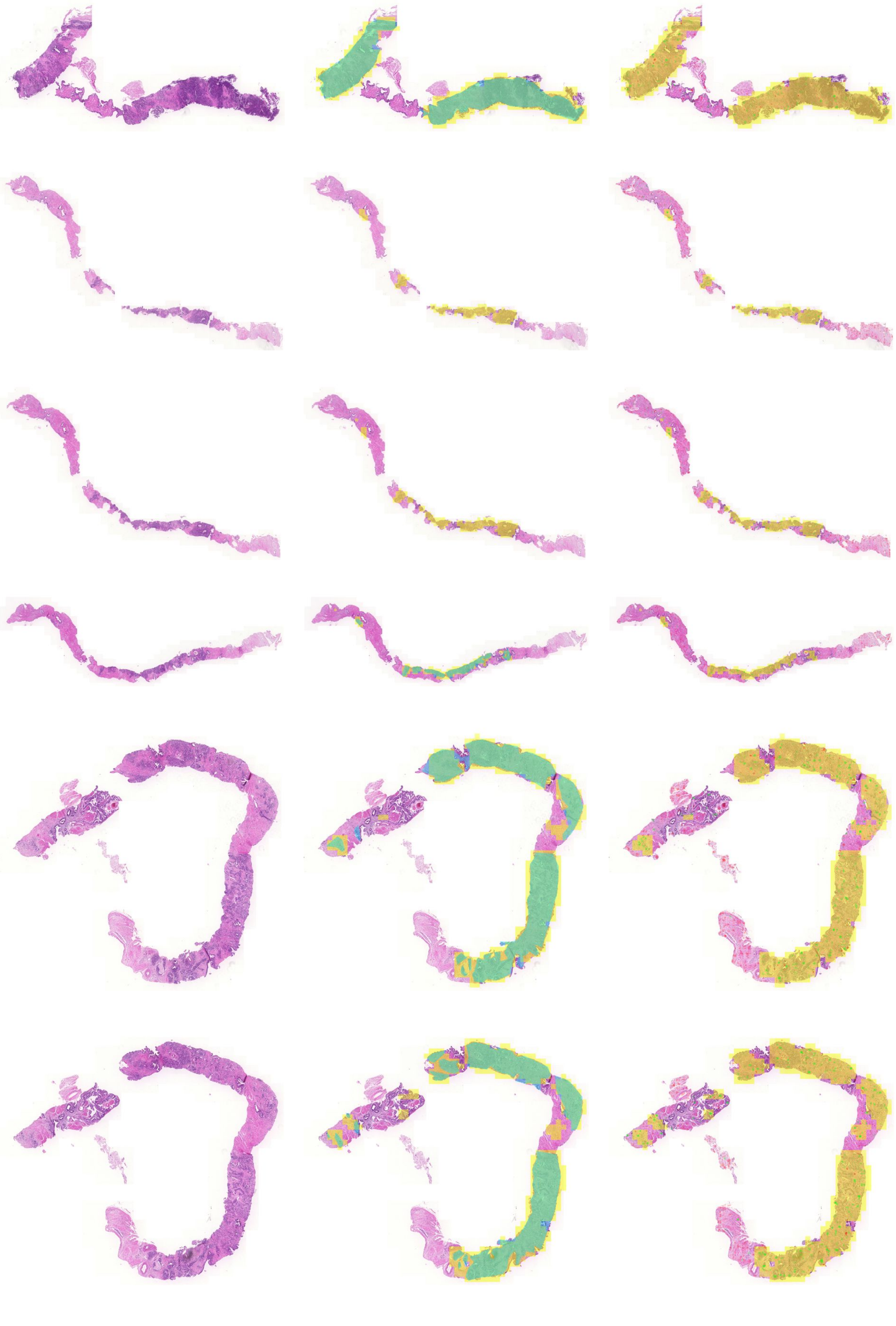

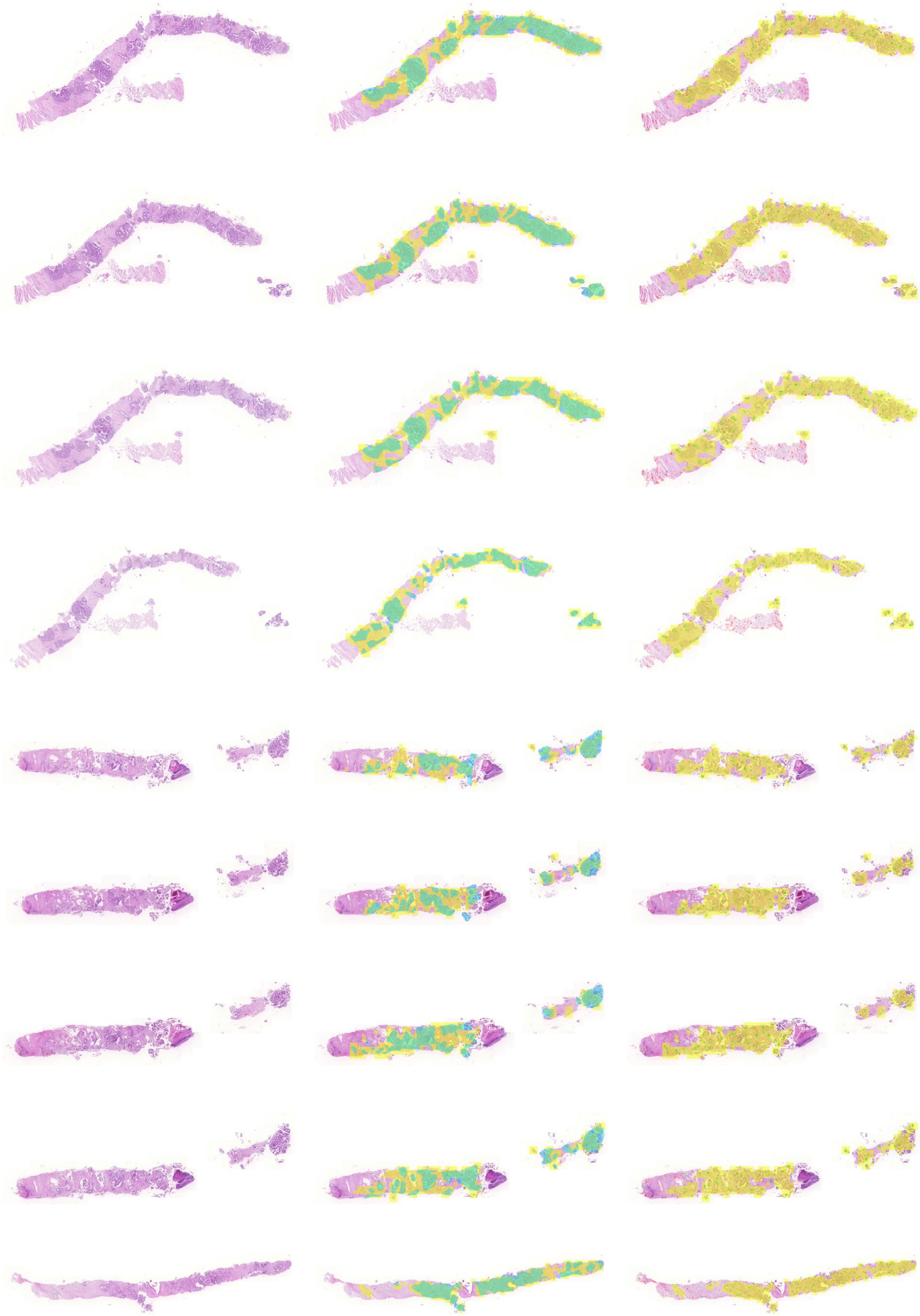

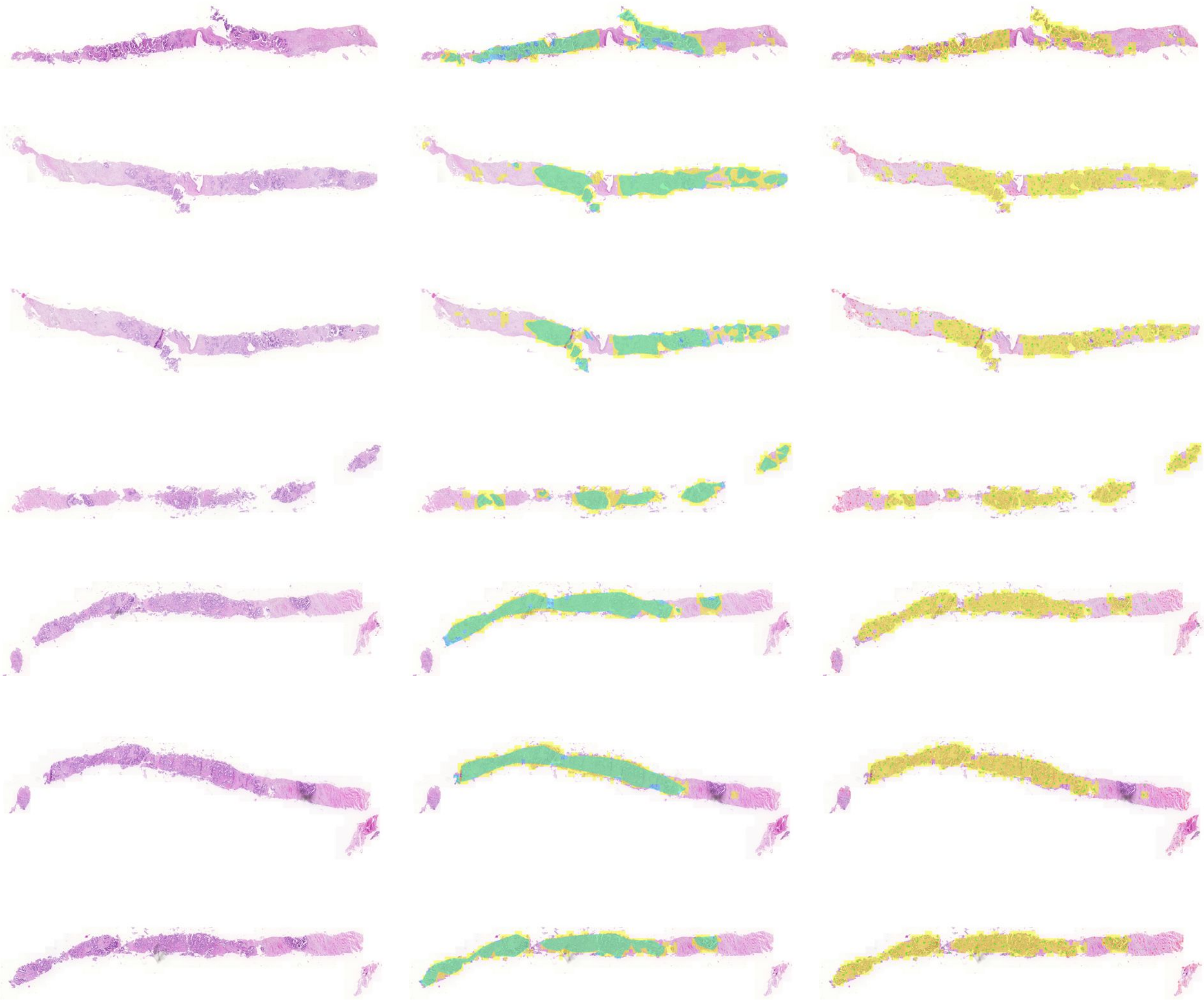

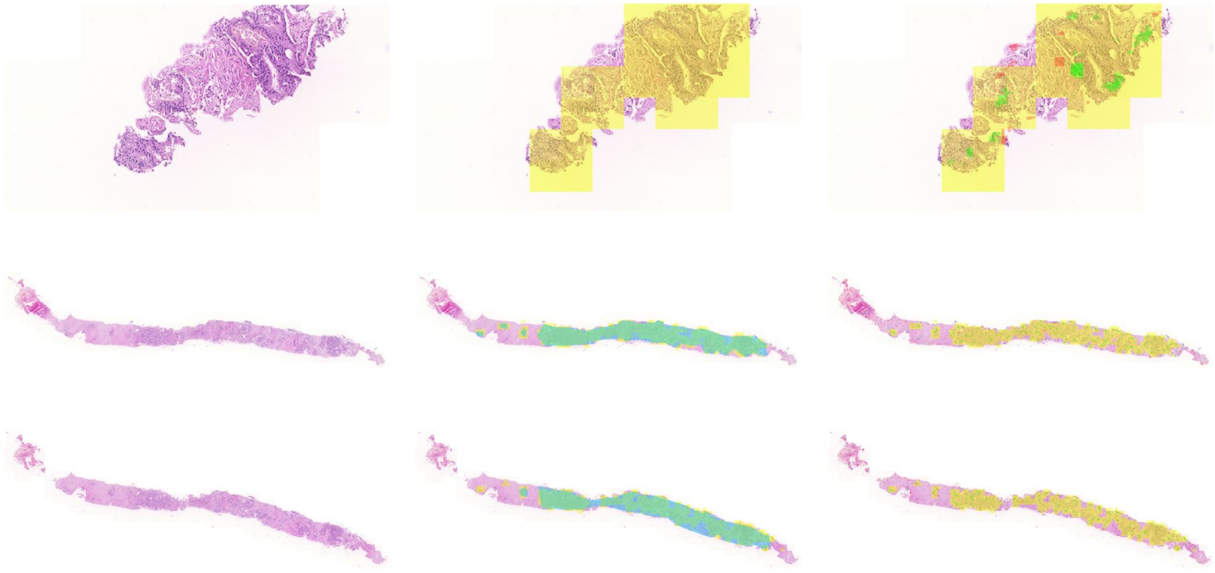

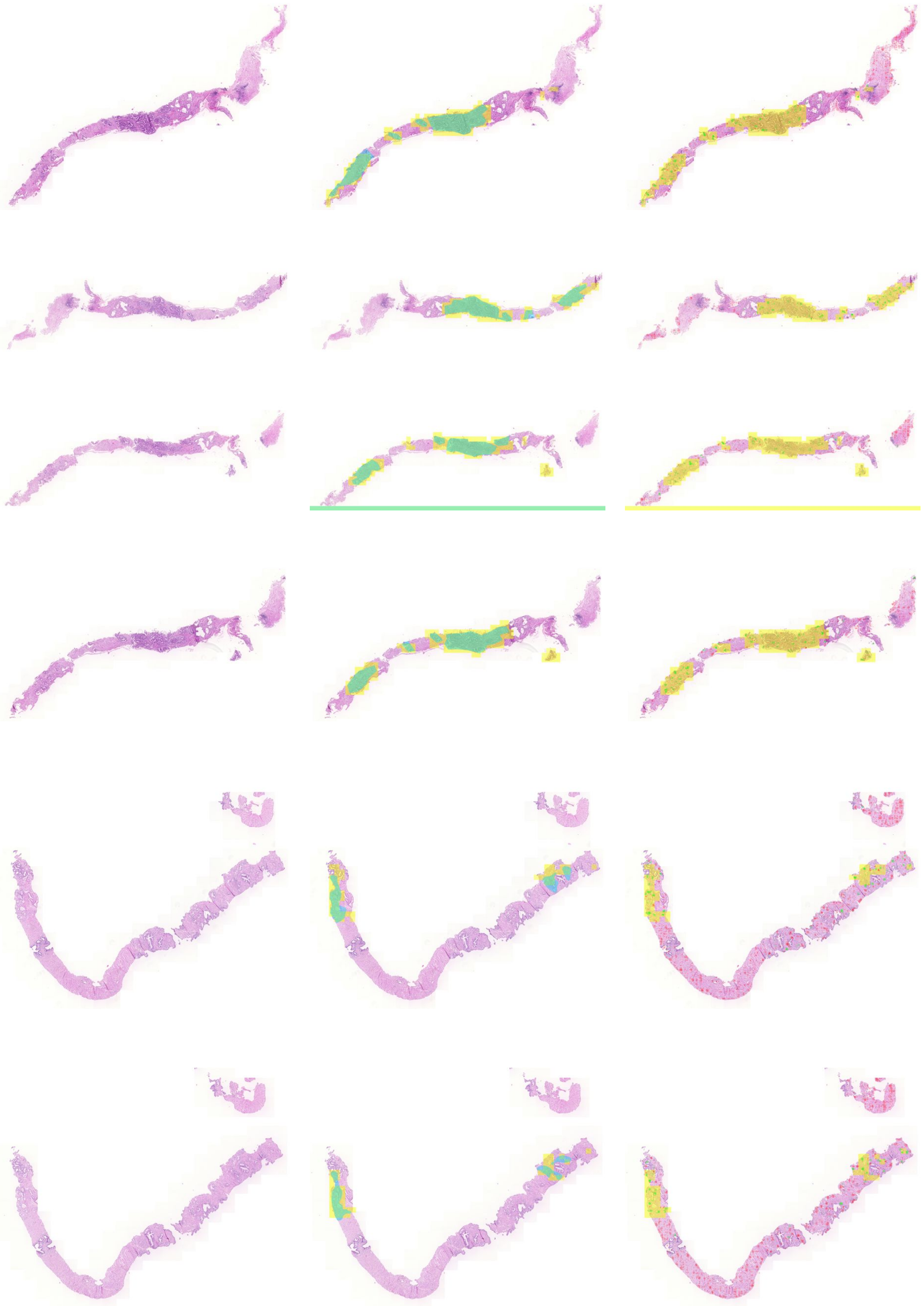

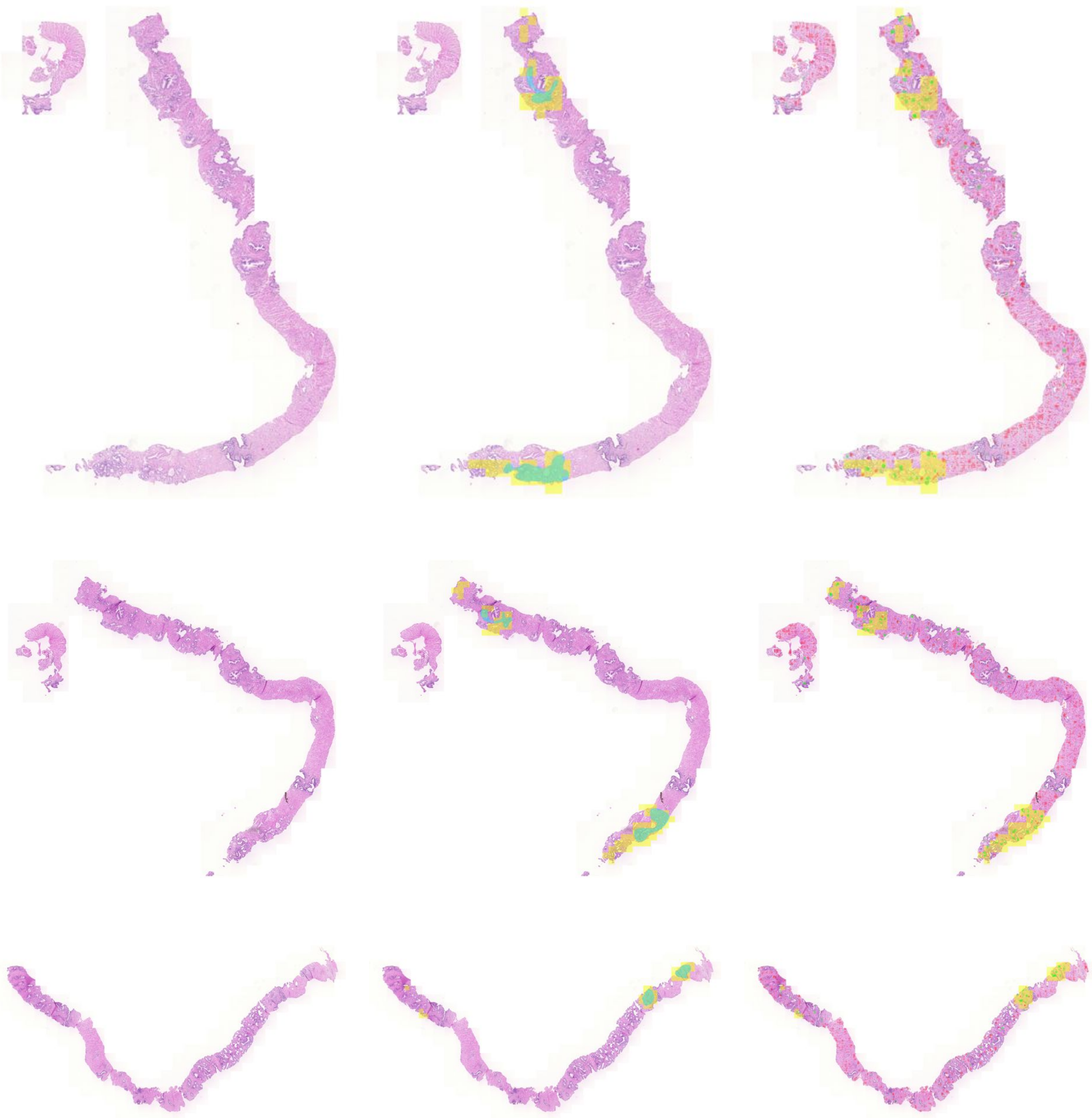

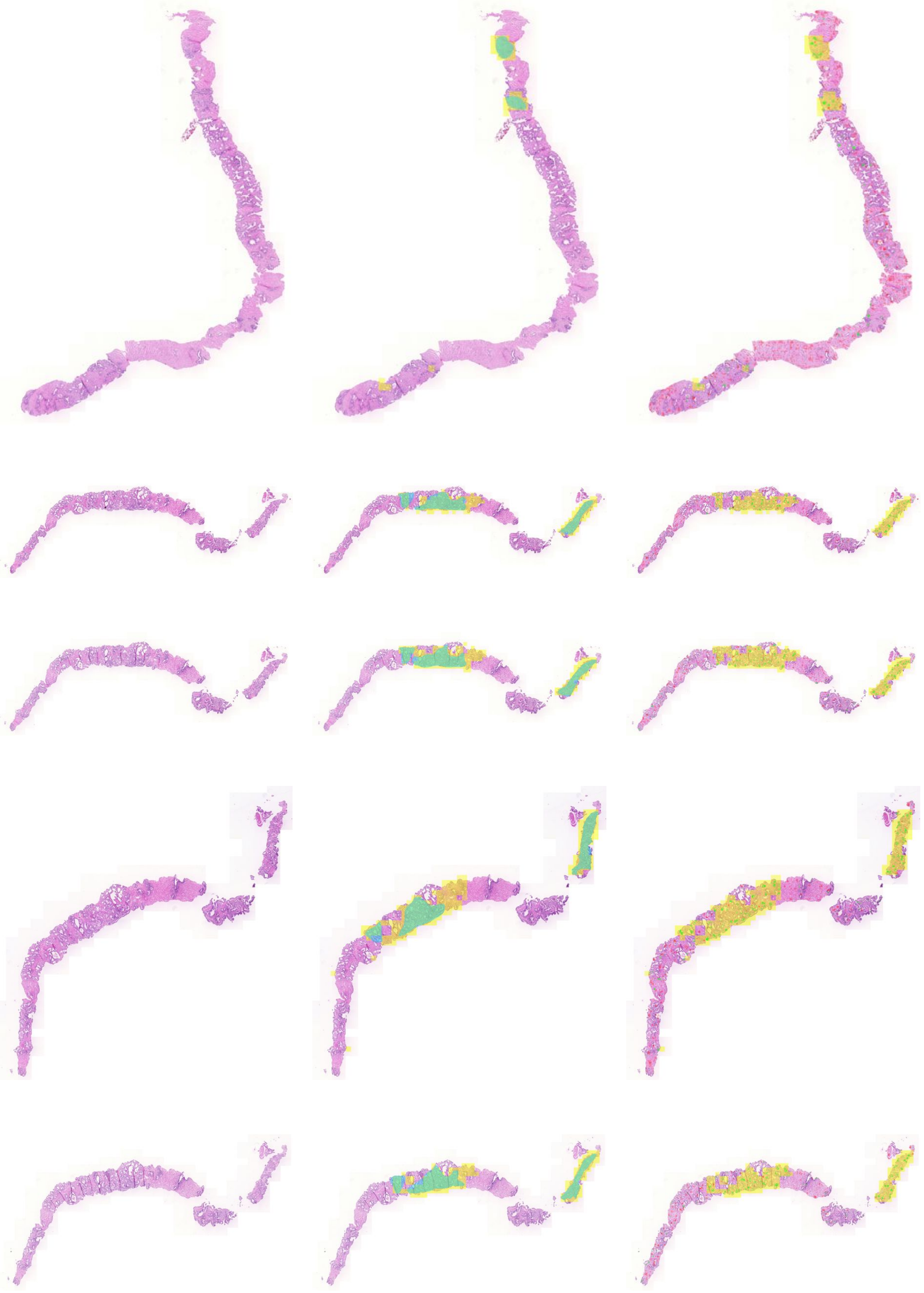

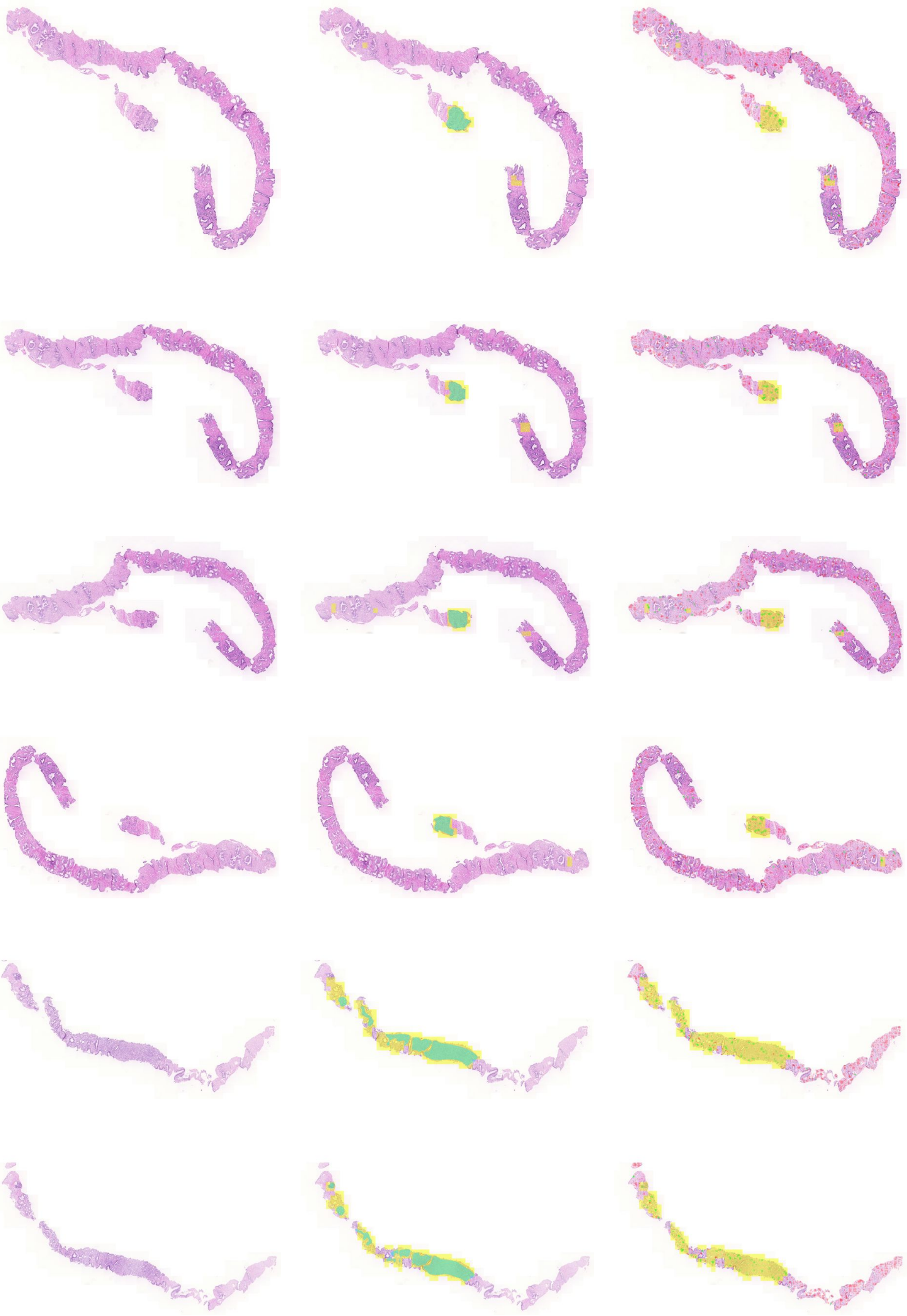

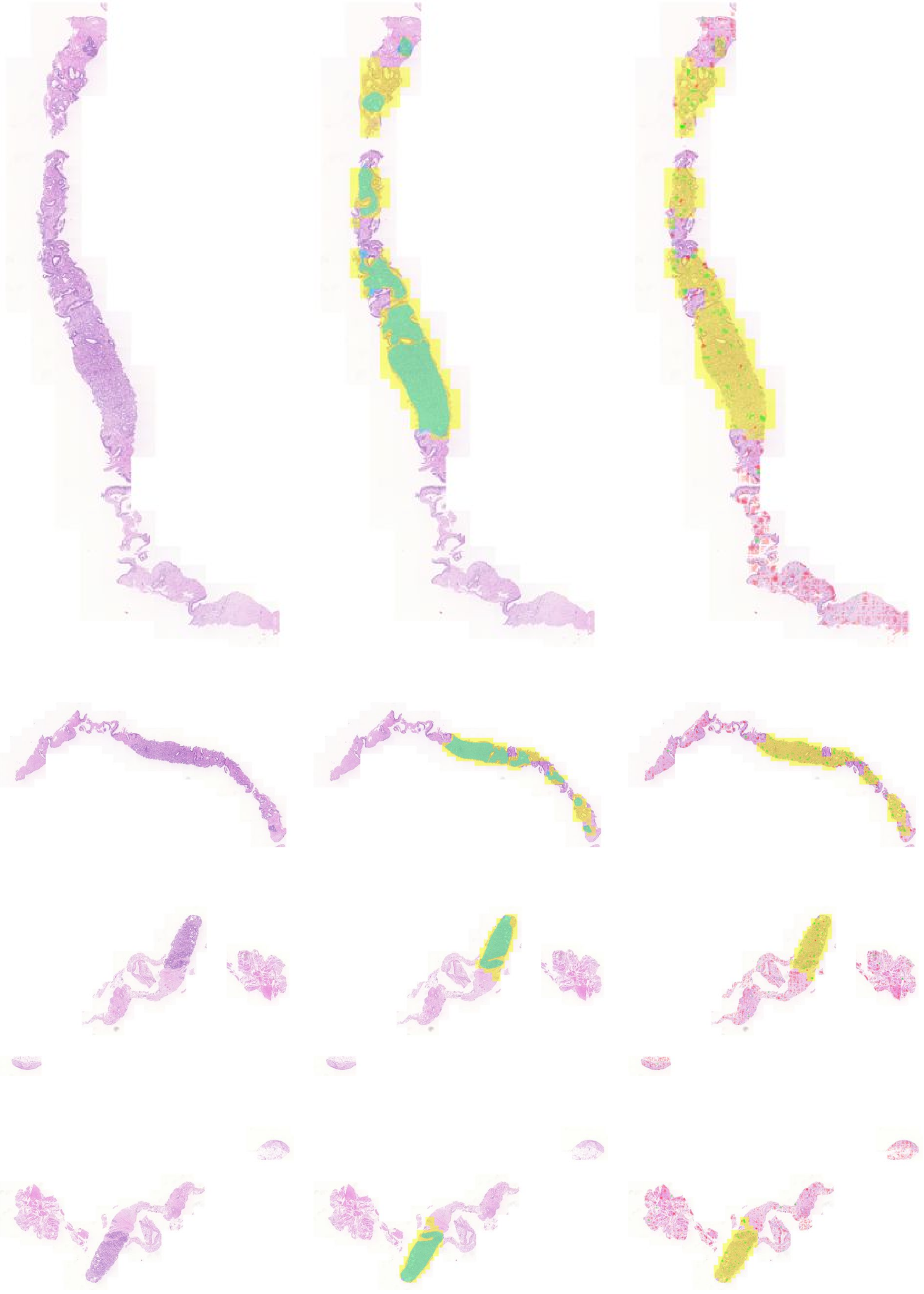

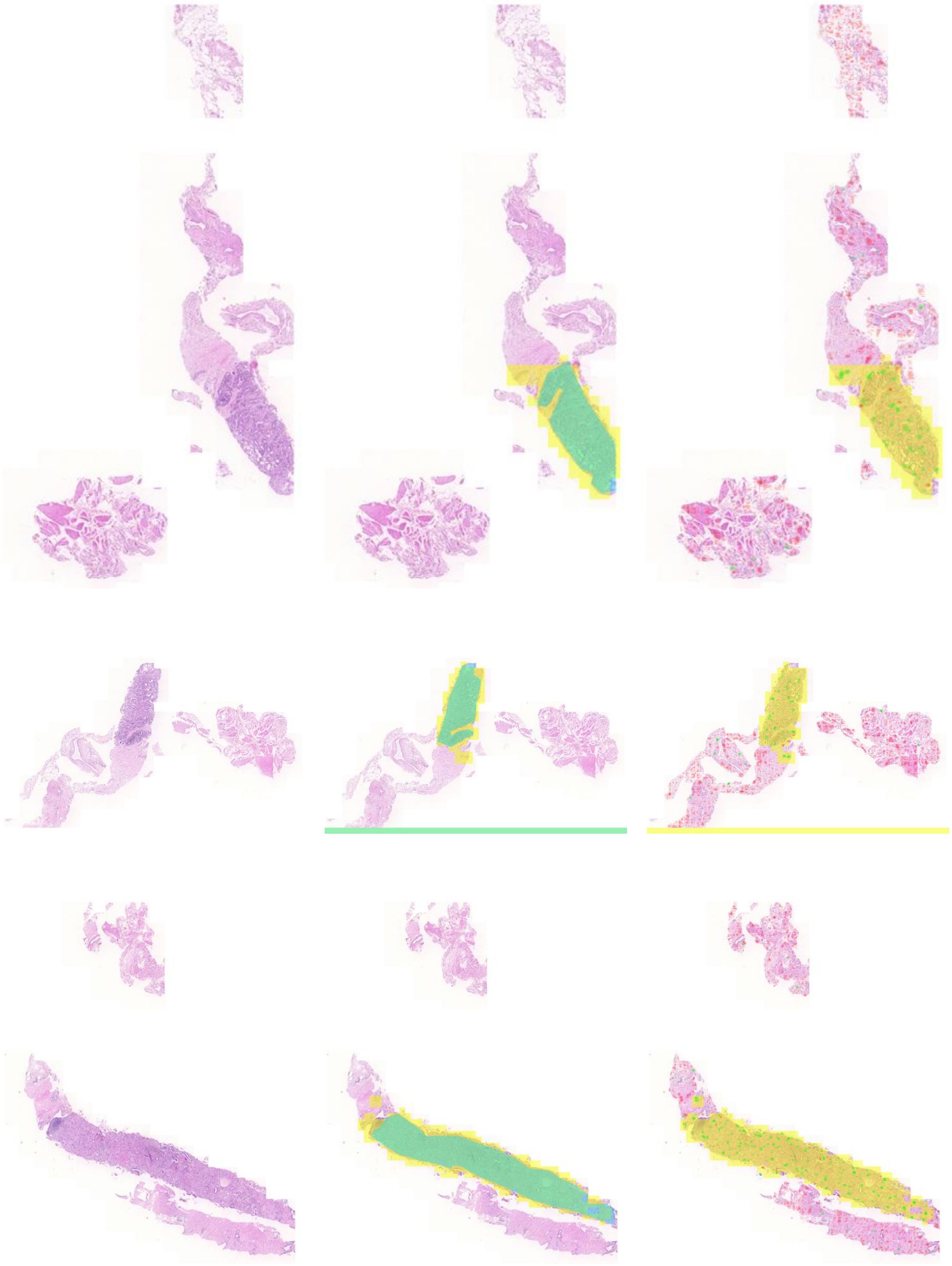

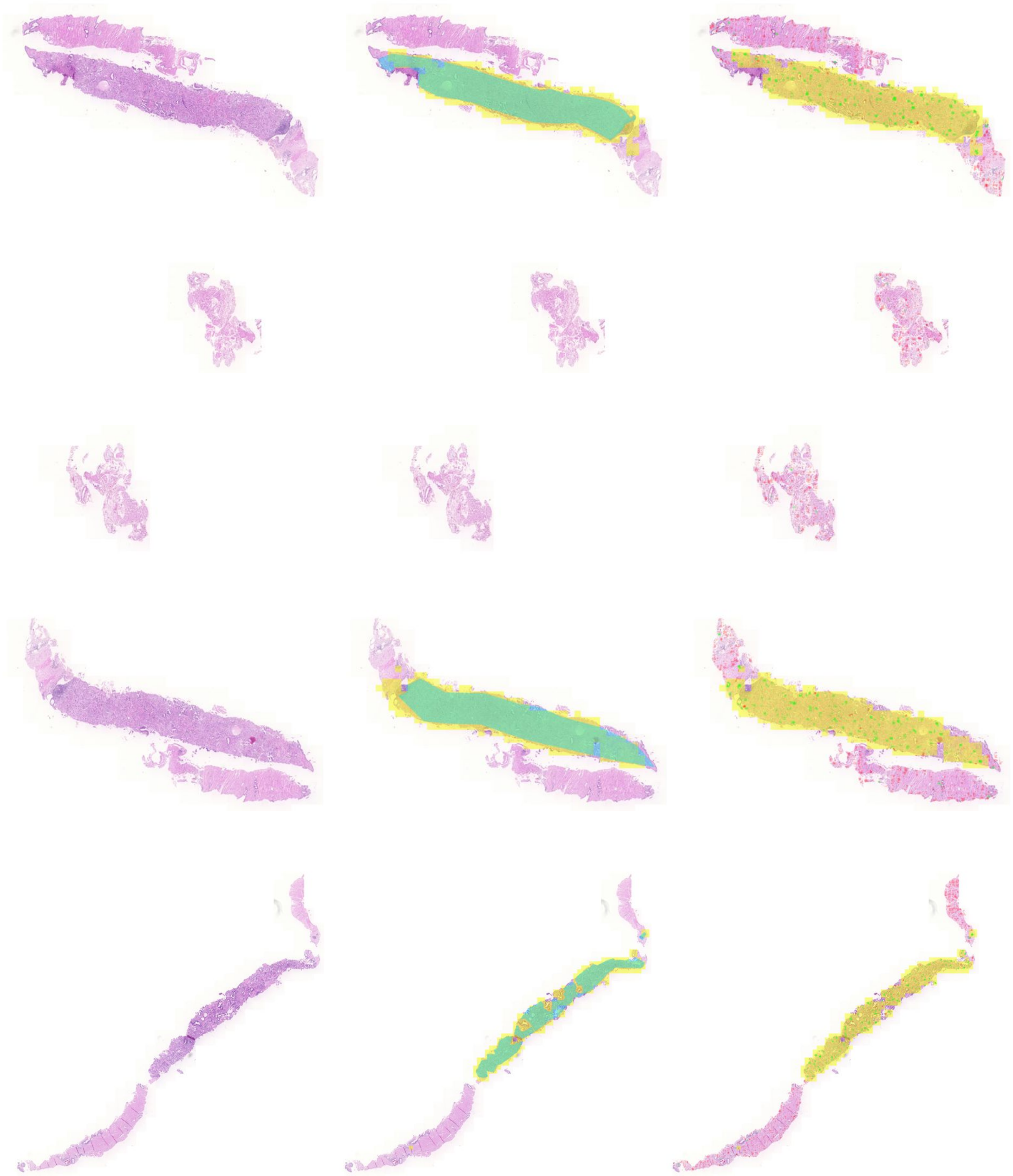

#### S3.2 Negative WSI

The following cases come from carcinoma-free WSI. From left to right, the images presented contain: left images show a portion of the original WSI; right images show the same region overlaid by probability of cancer masks (yellow) and occlusion-based explanation masks (green and red, signifying positive and negative contribution towards cancer diagnosis, respectively).

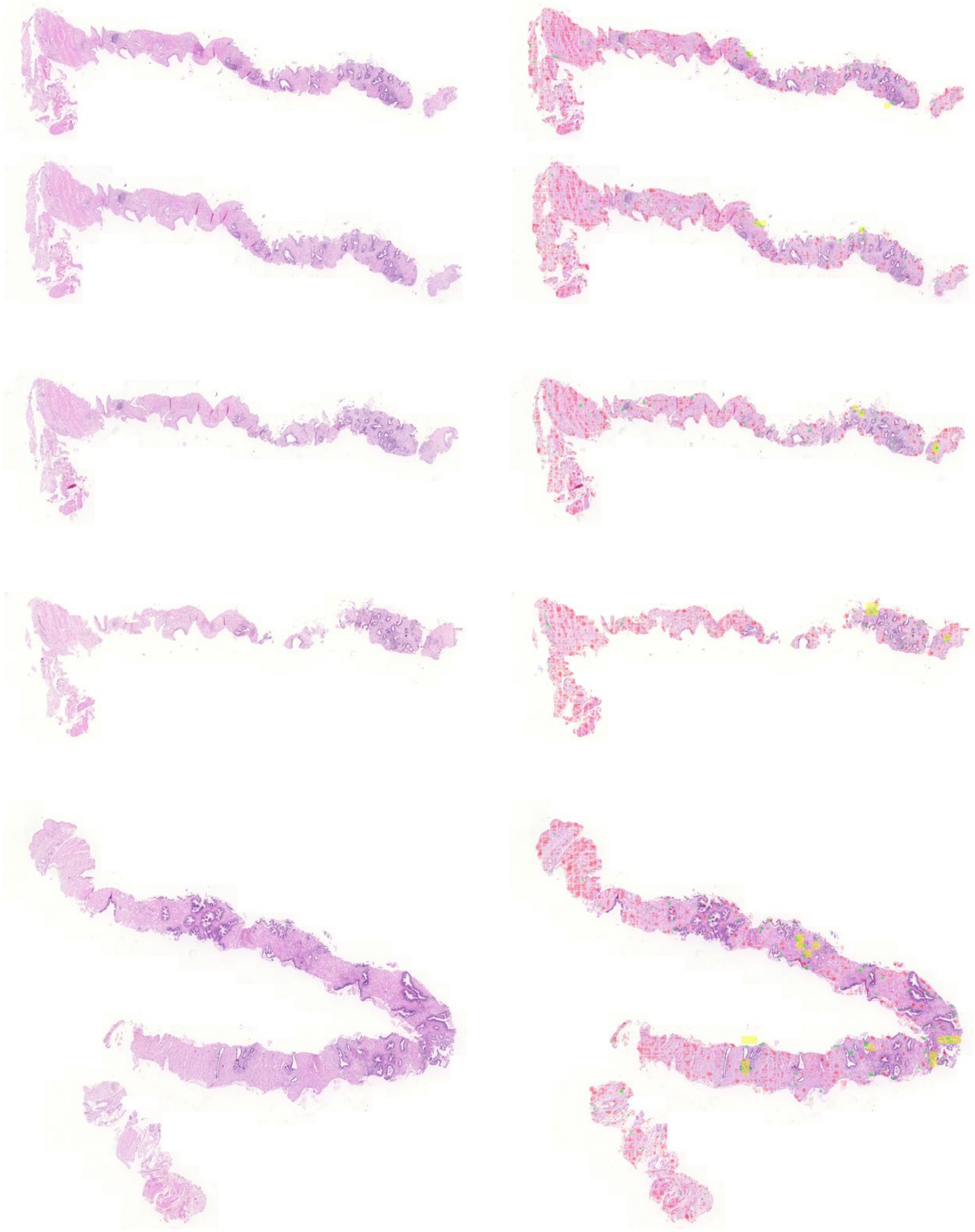

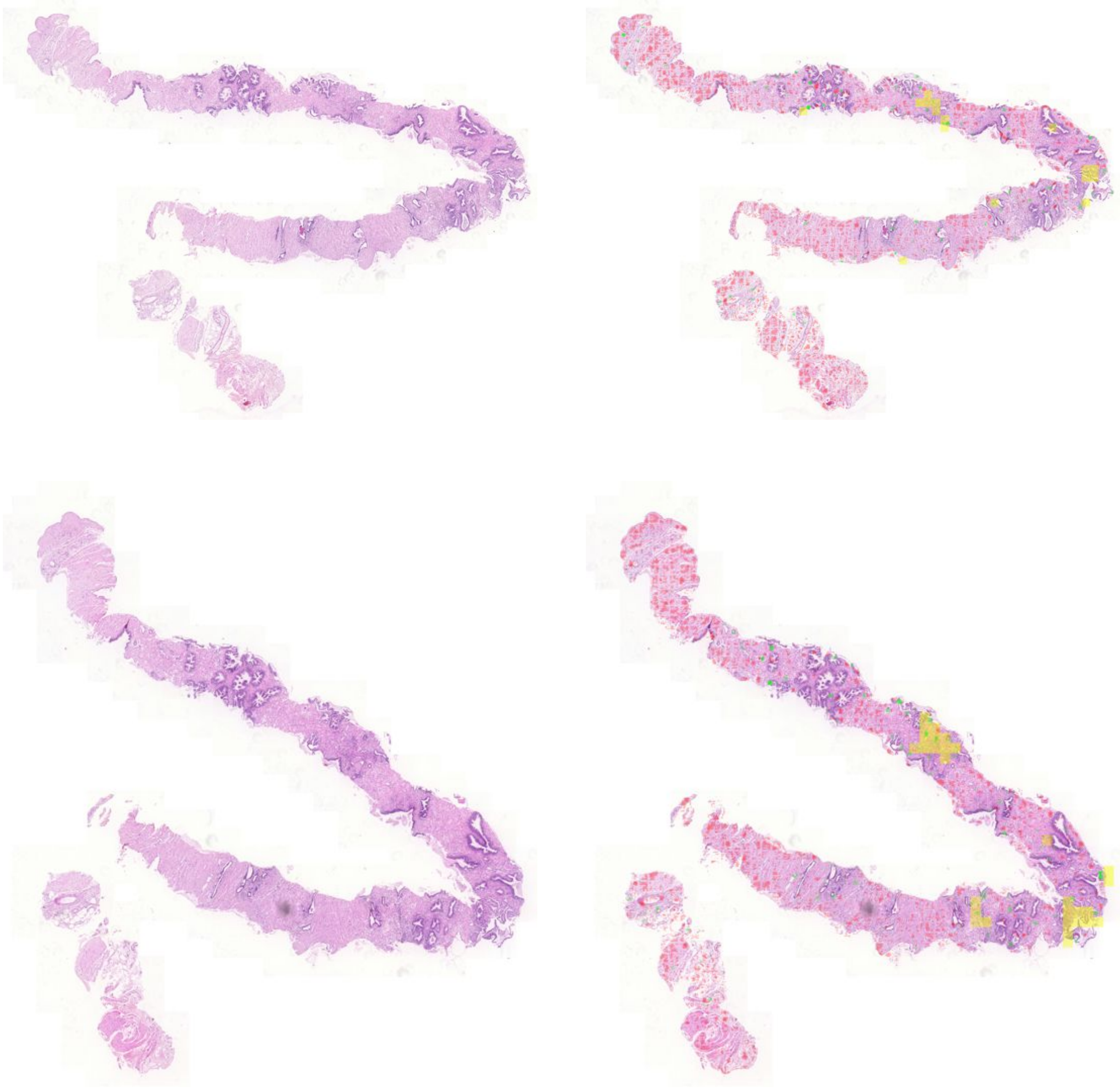

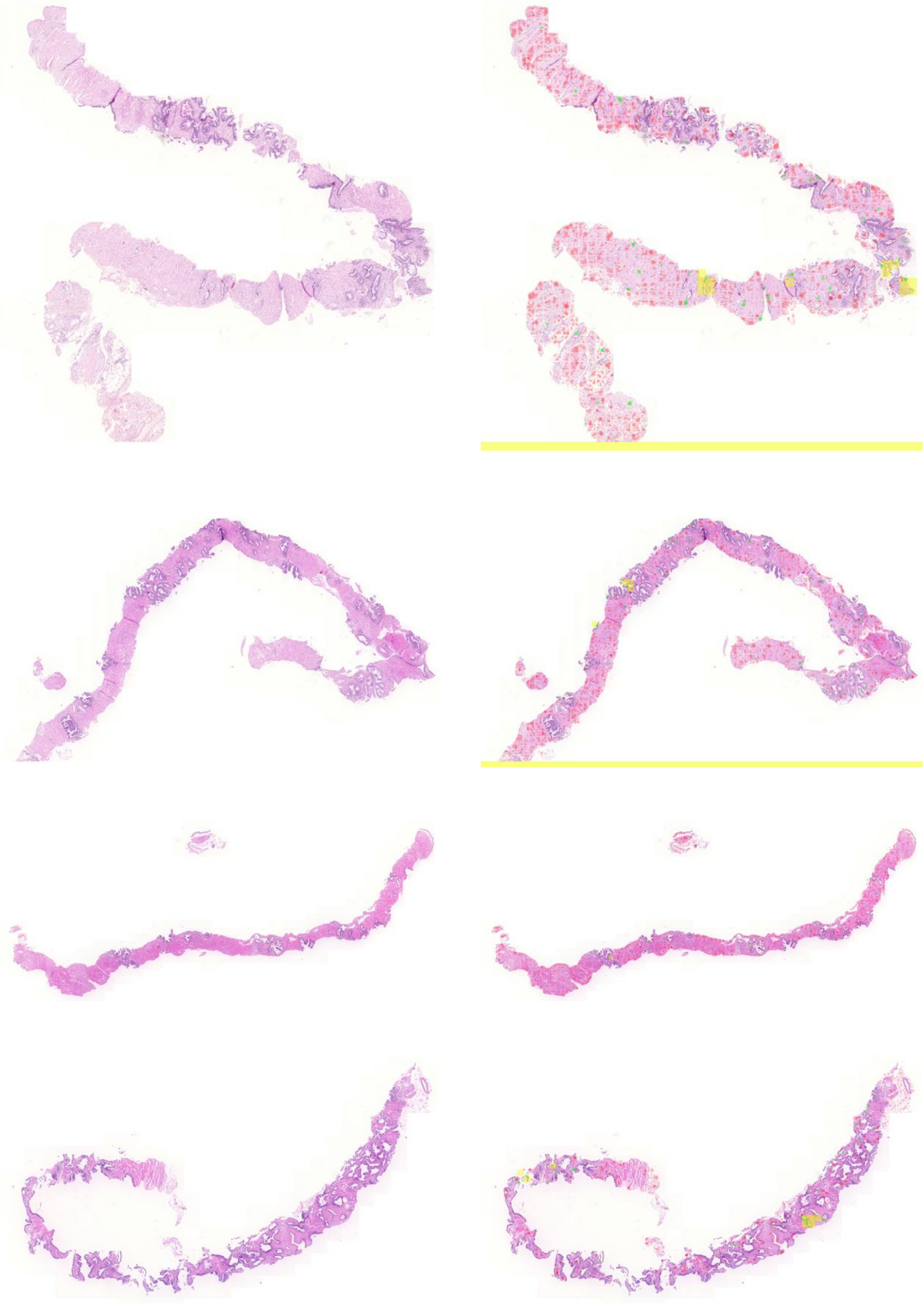

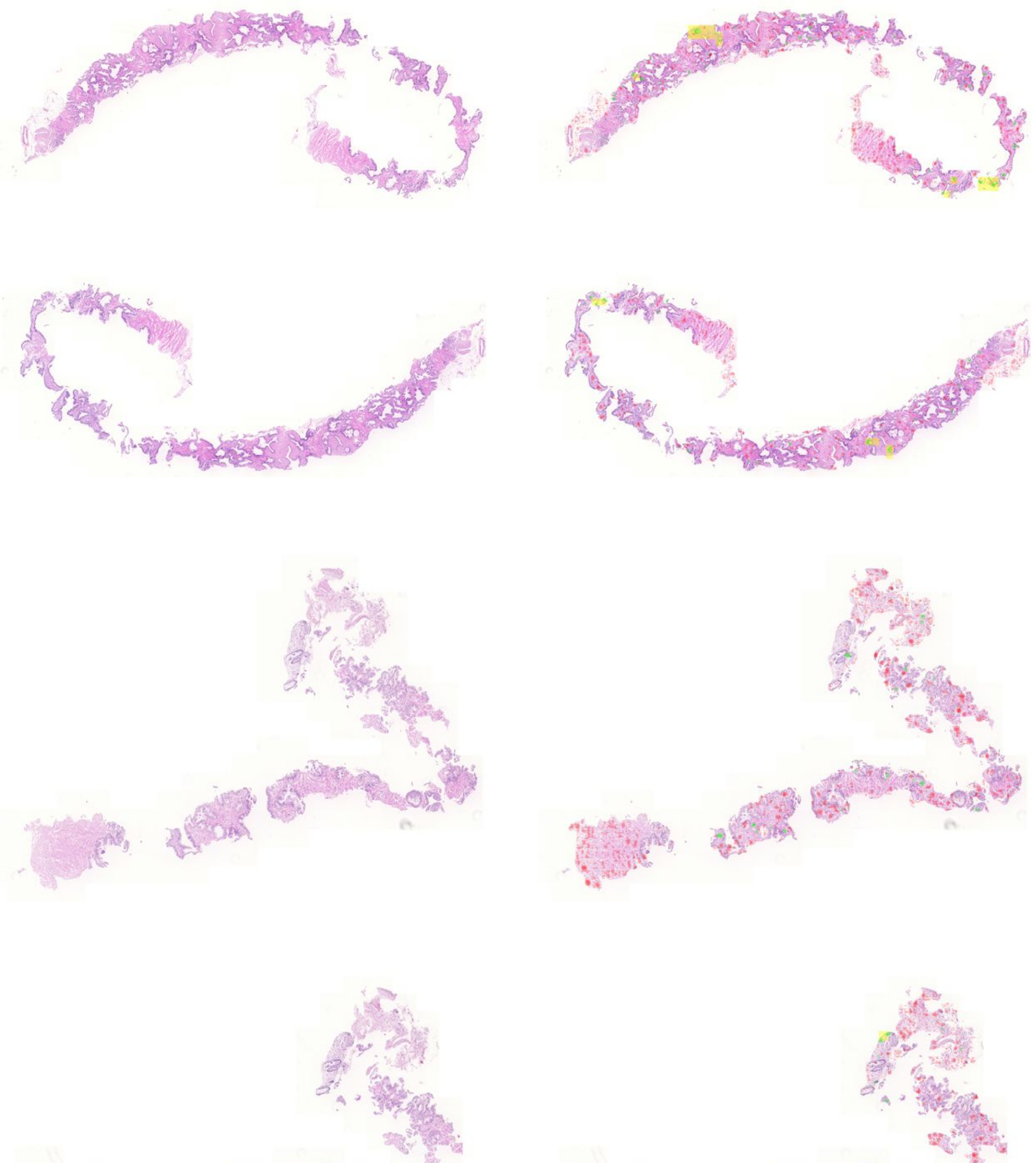

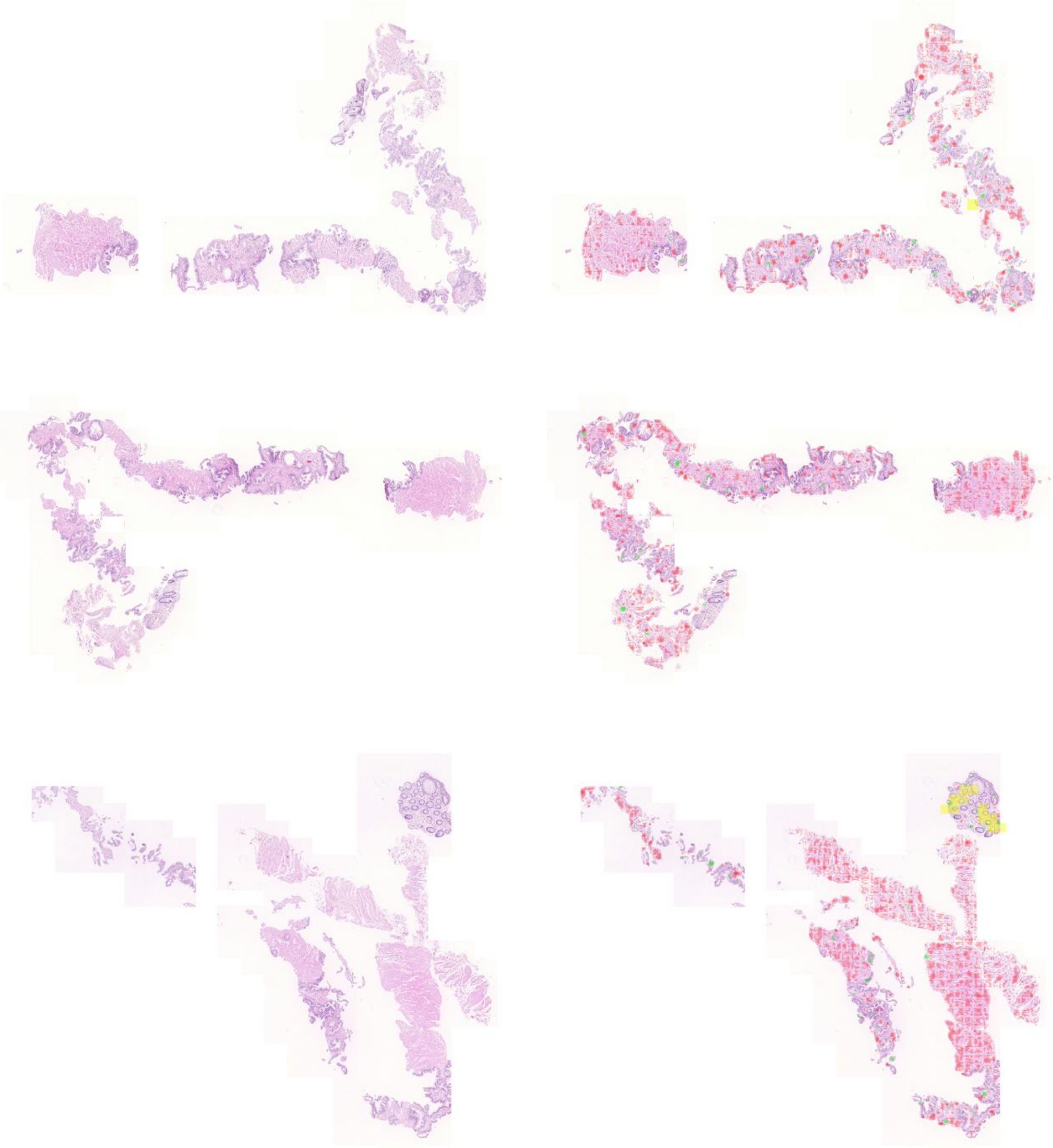

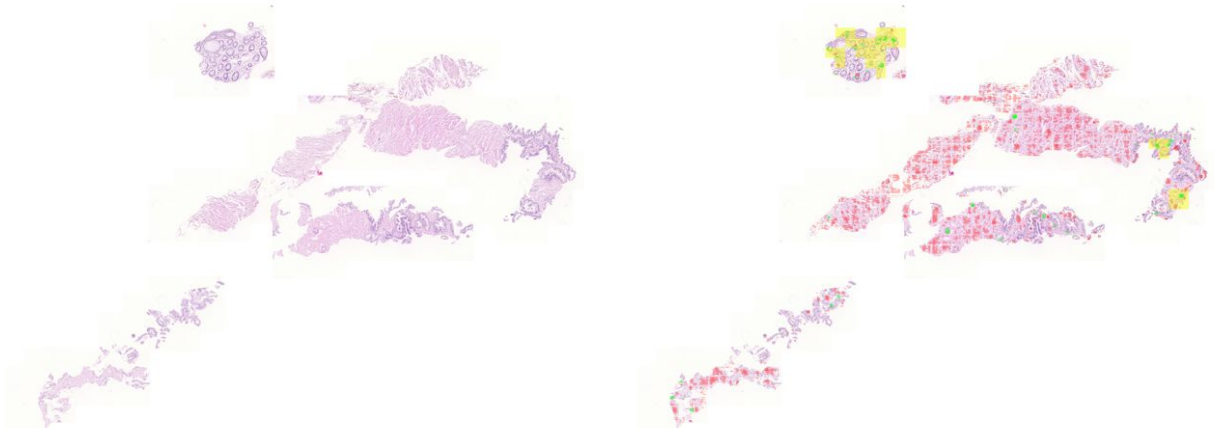

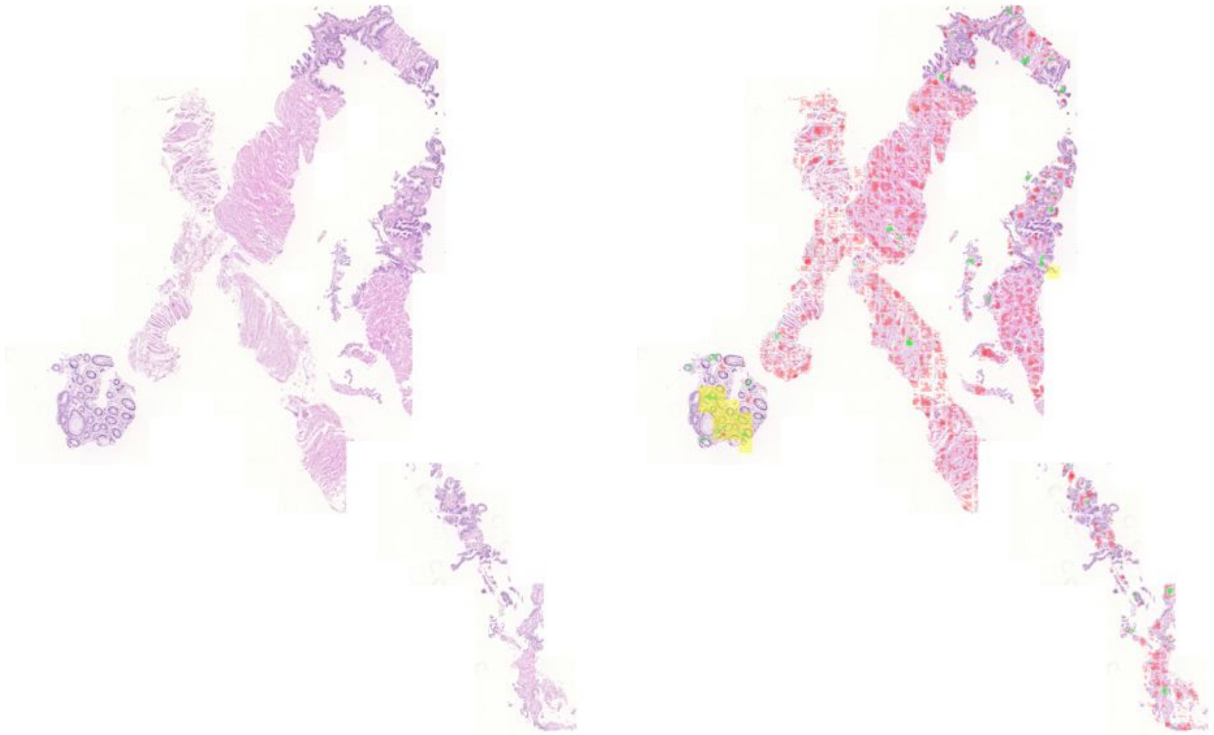

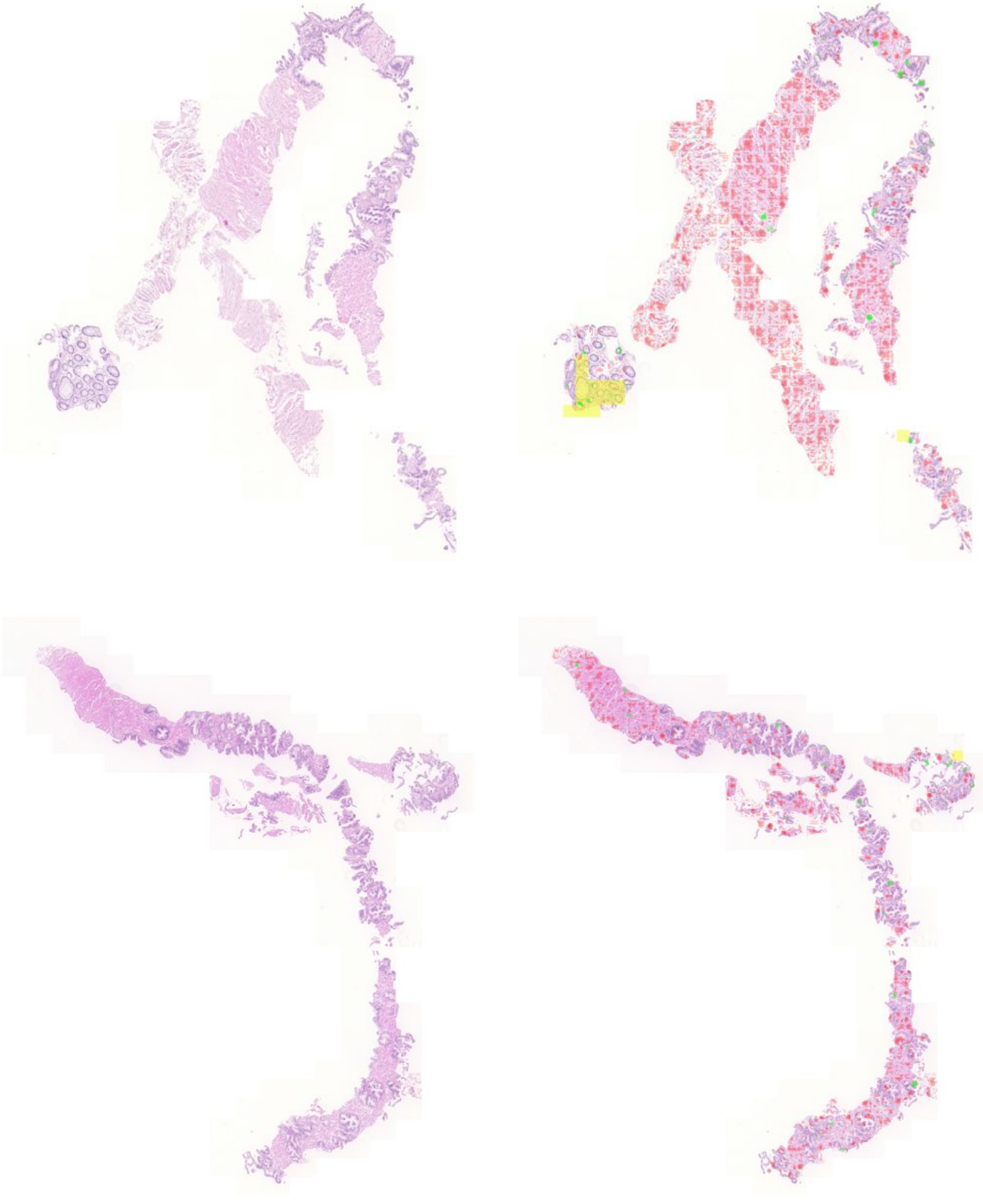

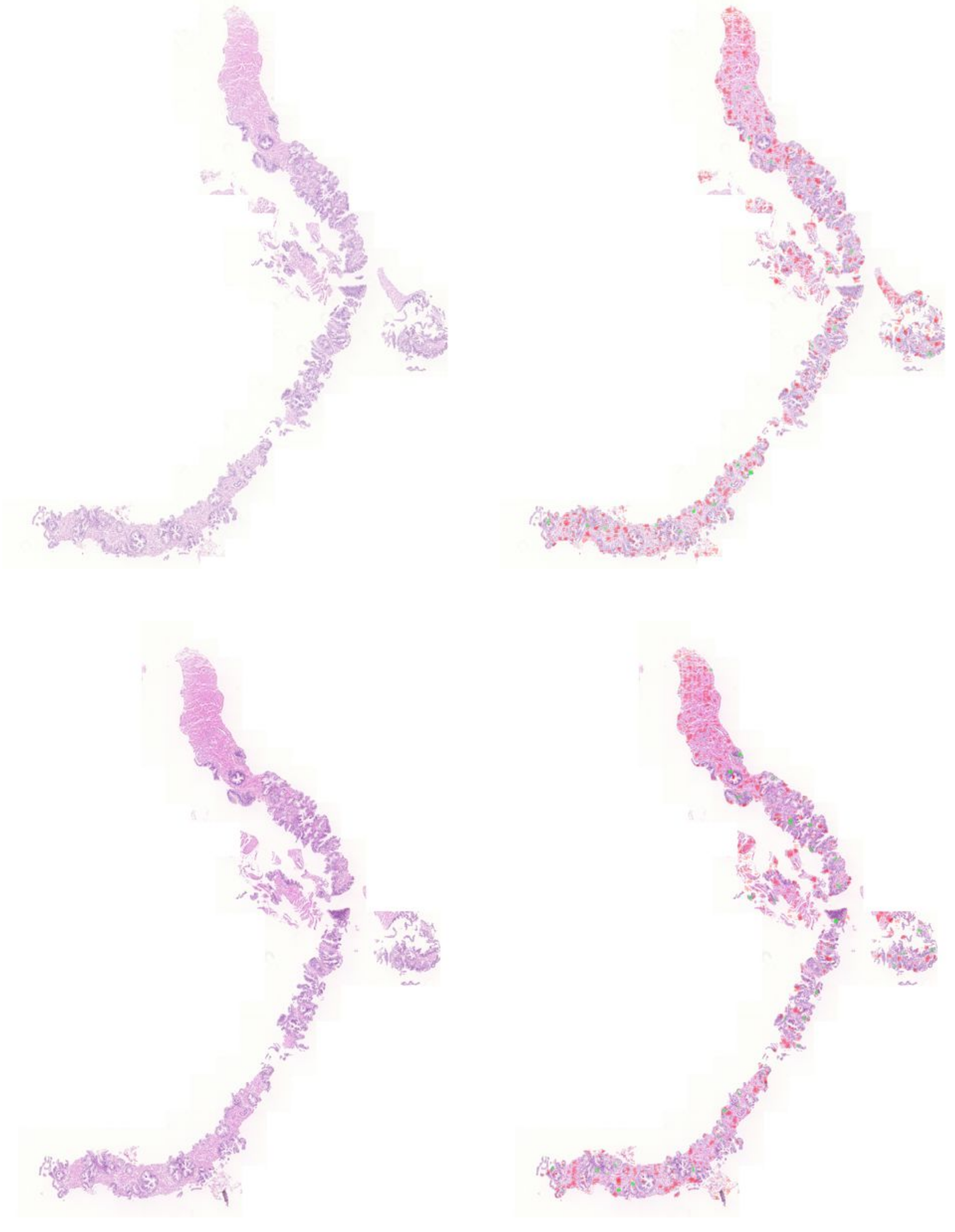

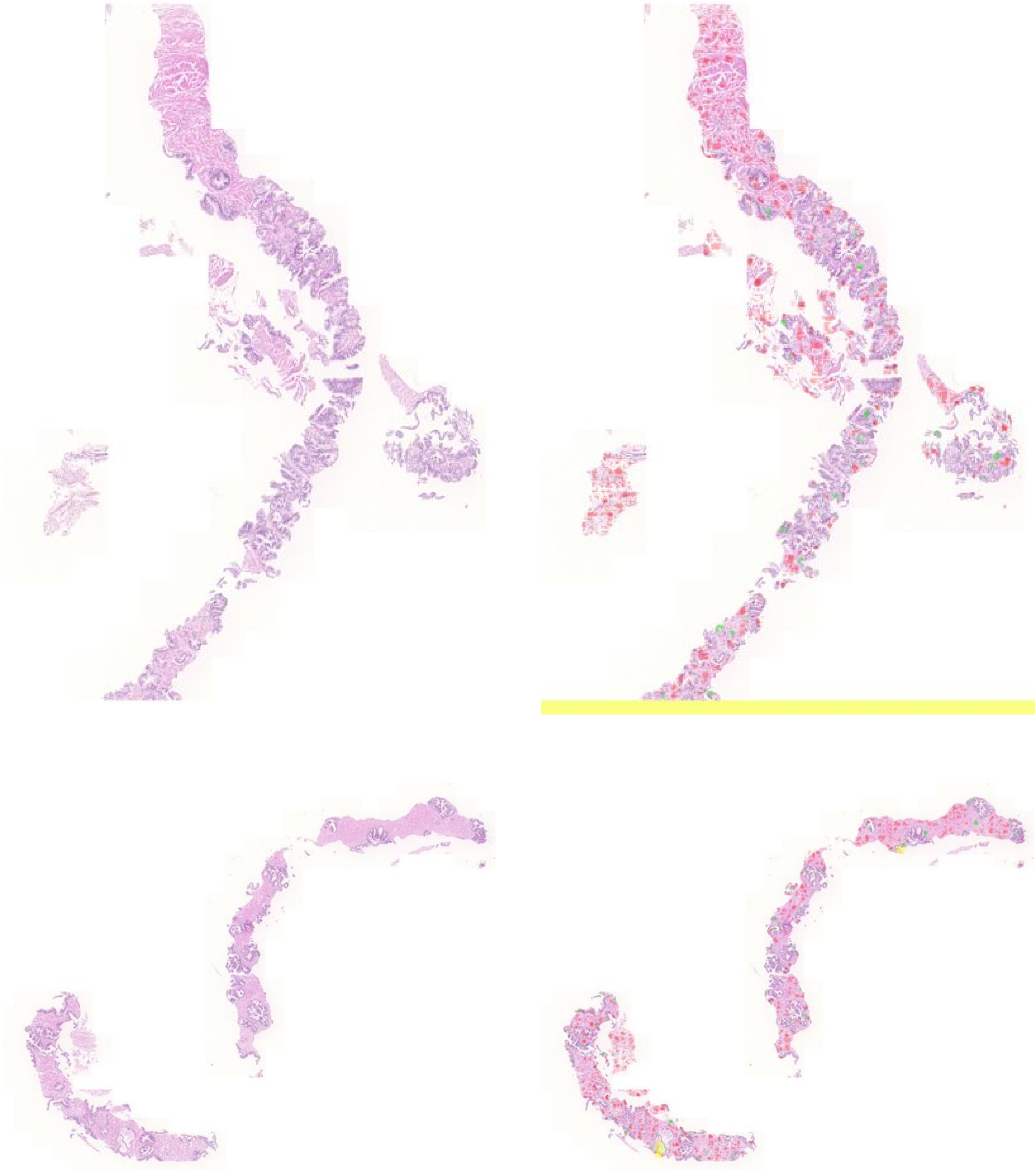

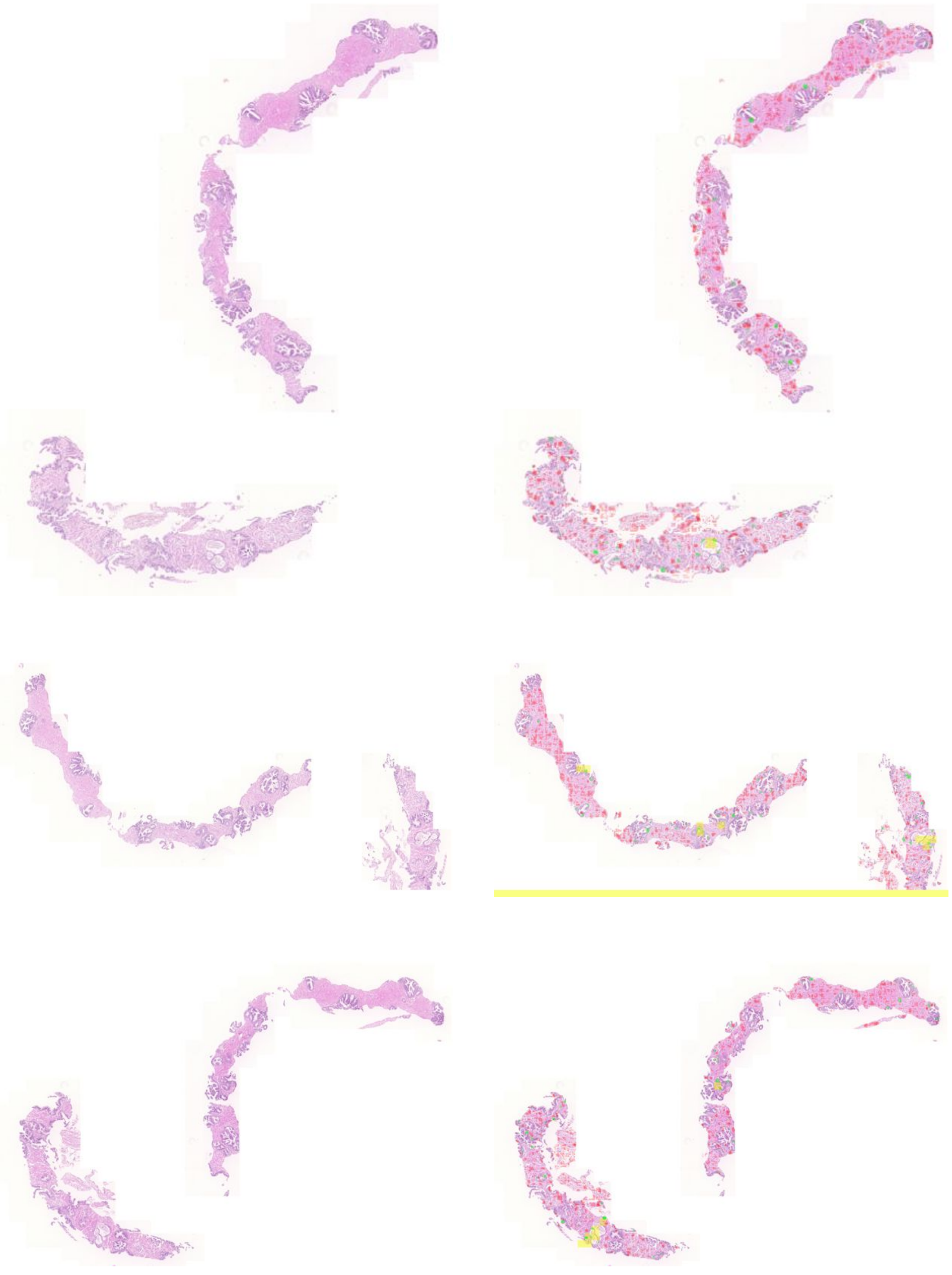

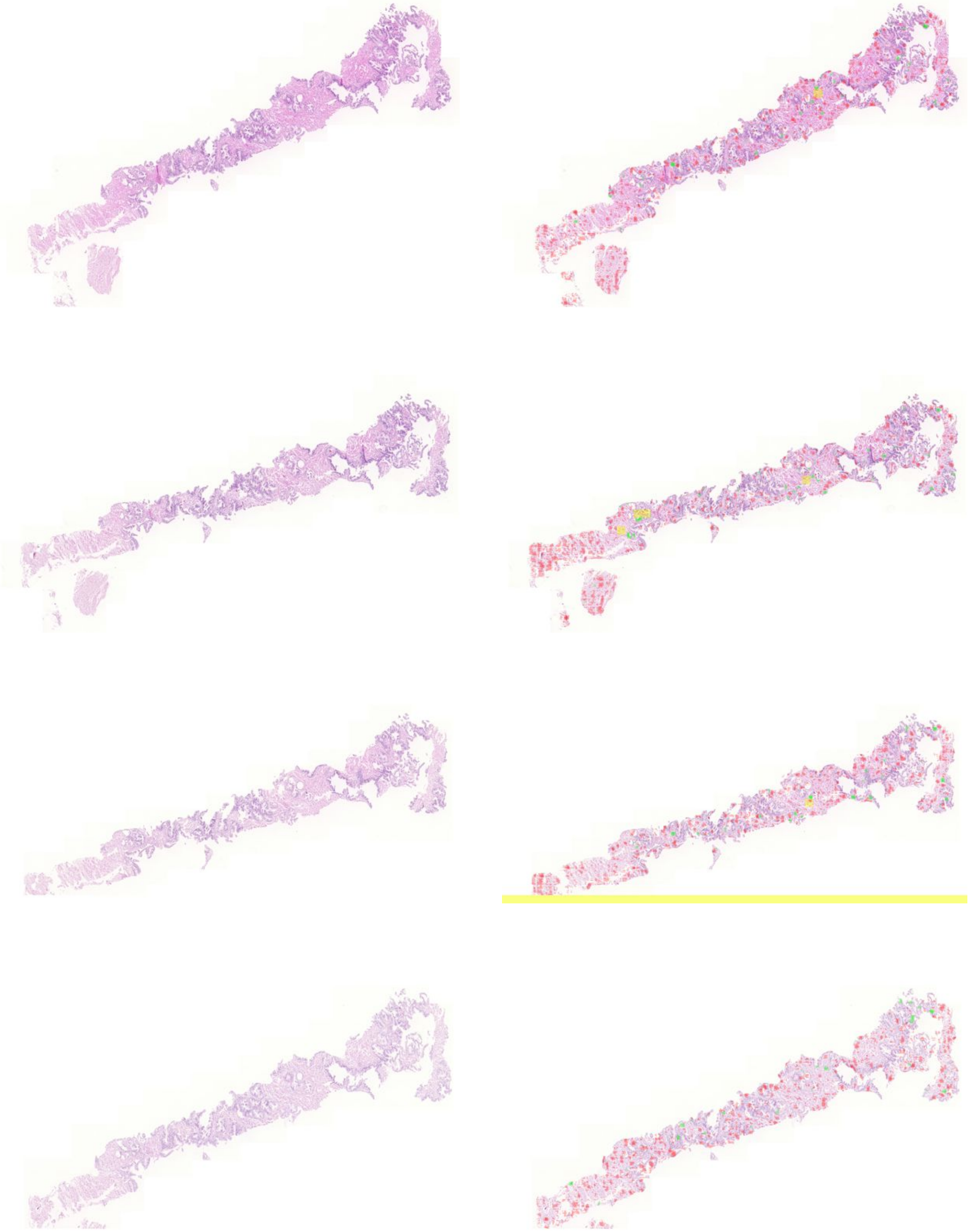

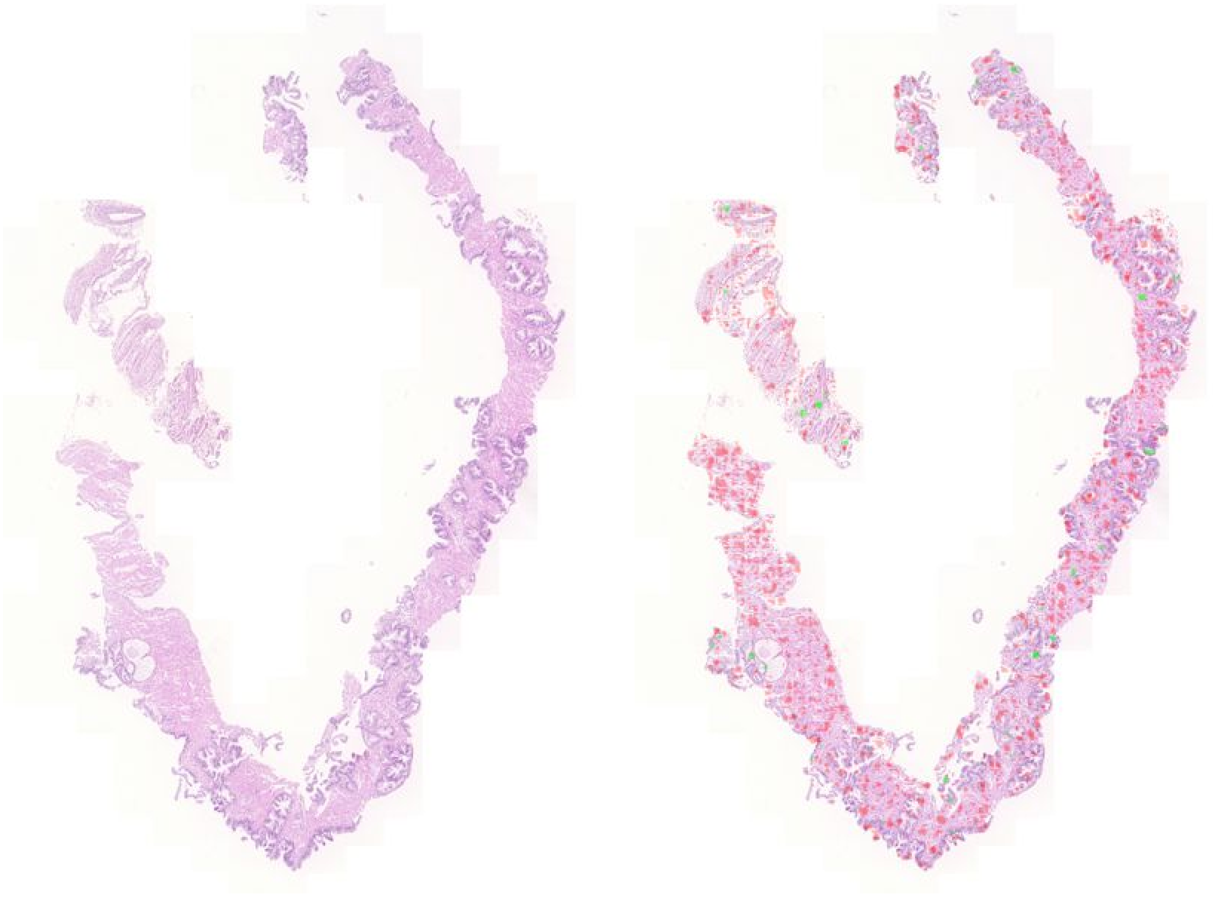

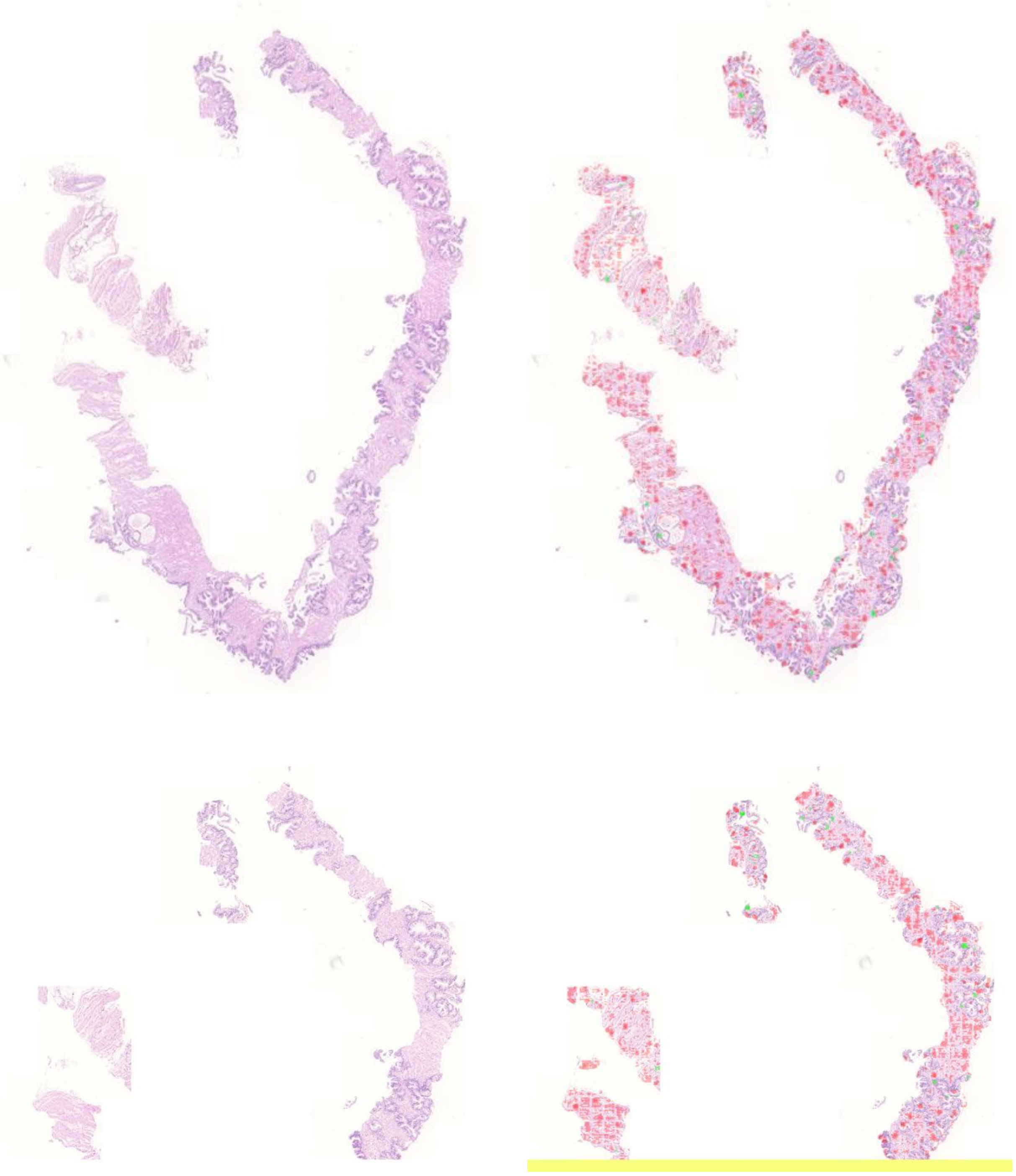

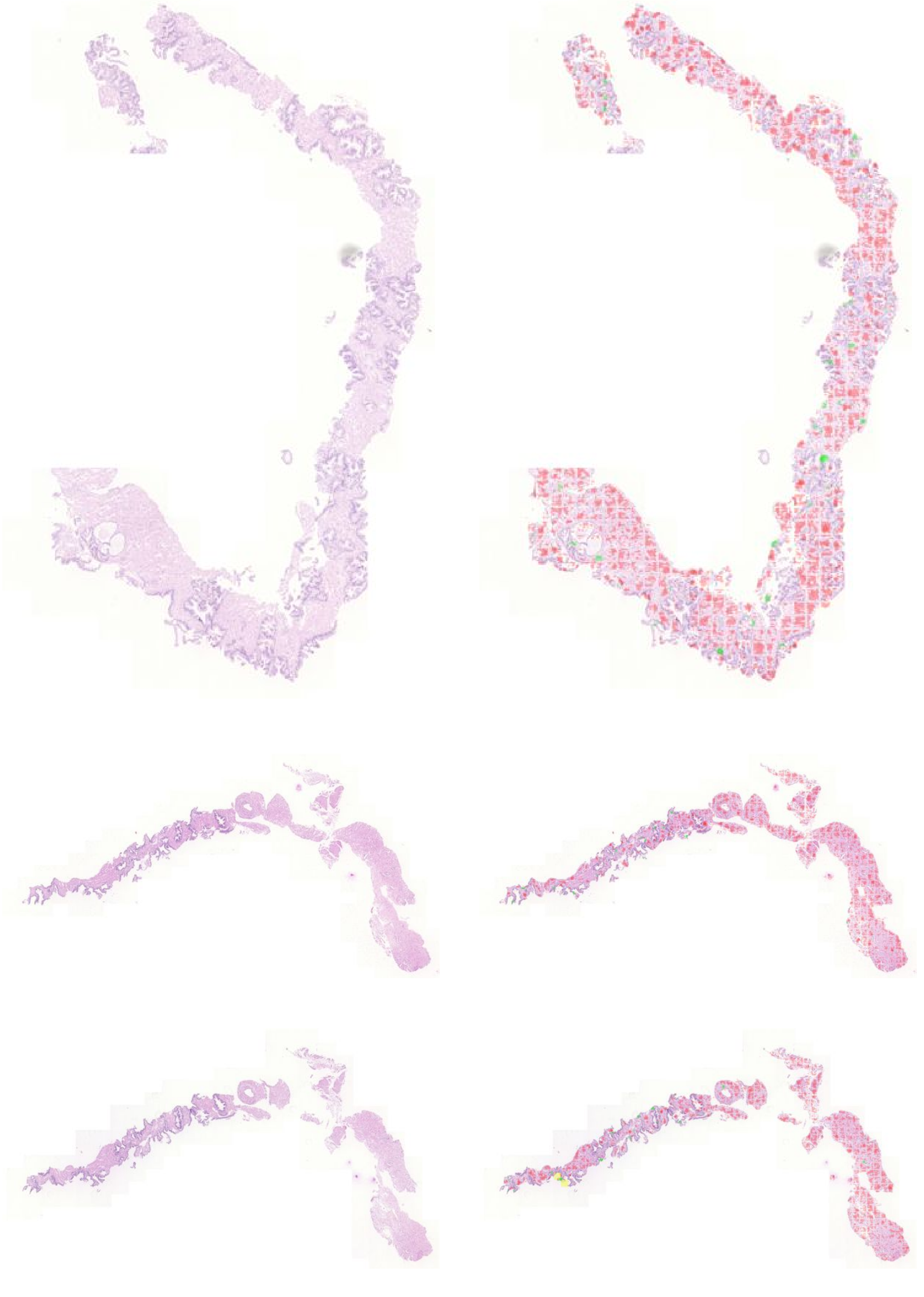

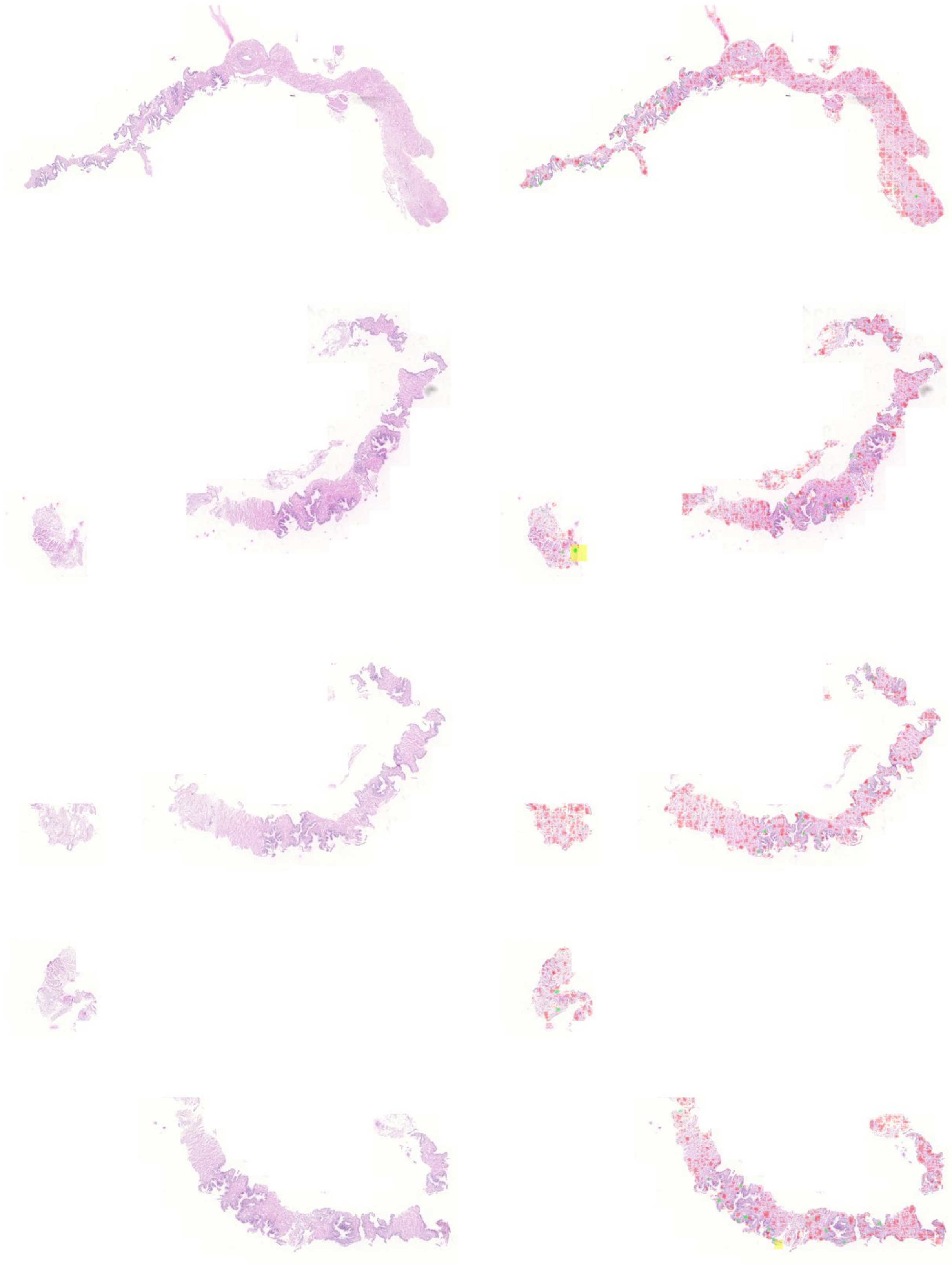

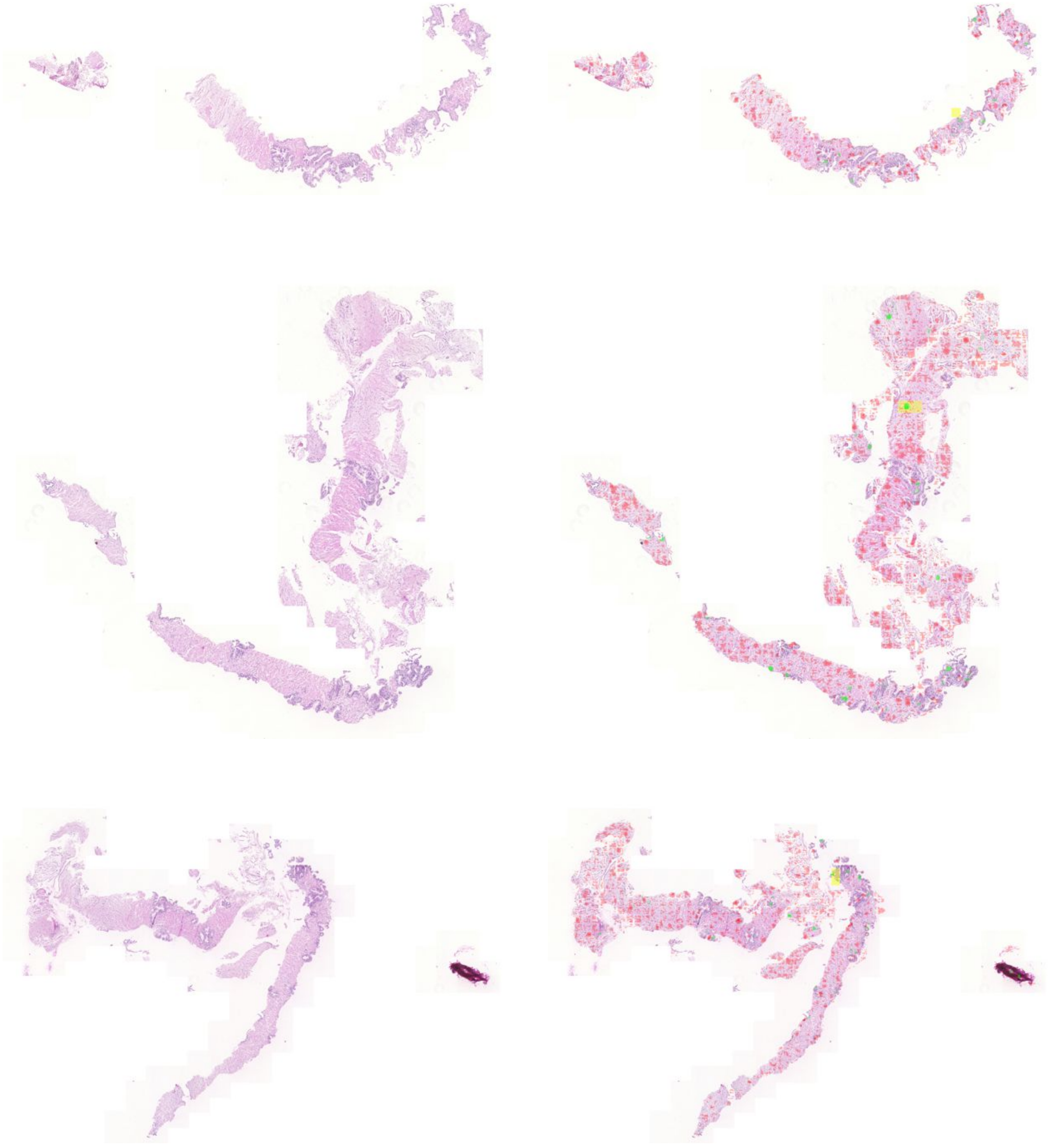

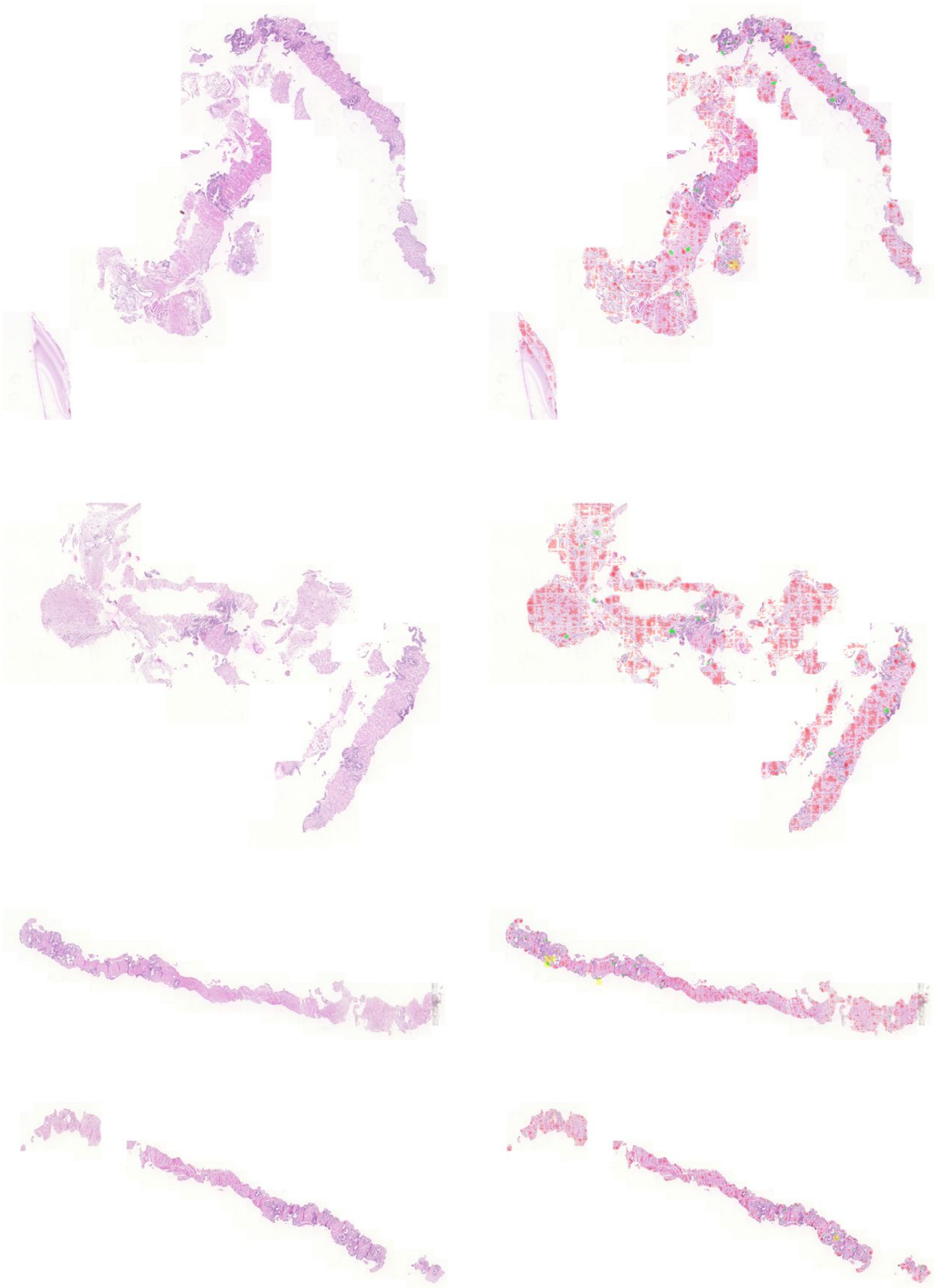

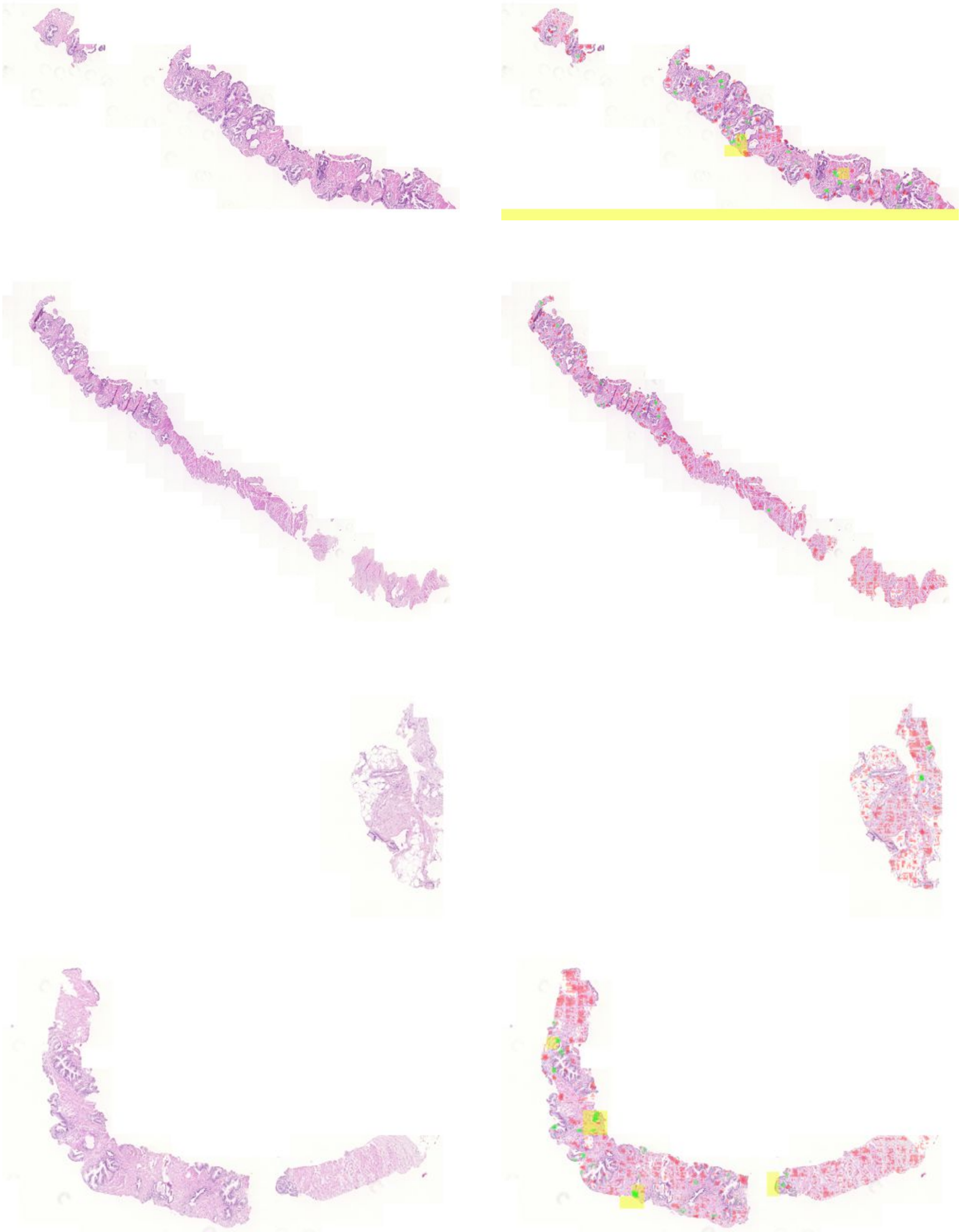

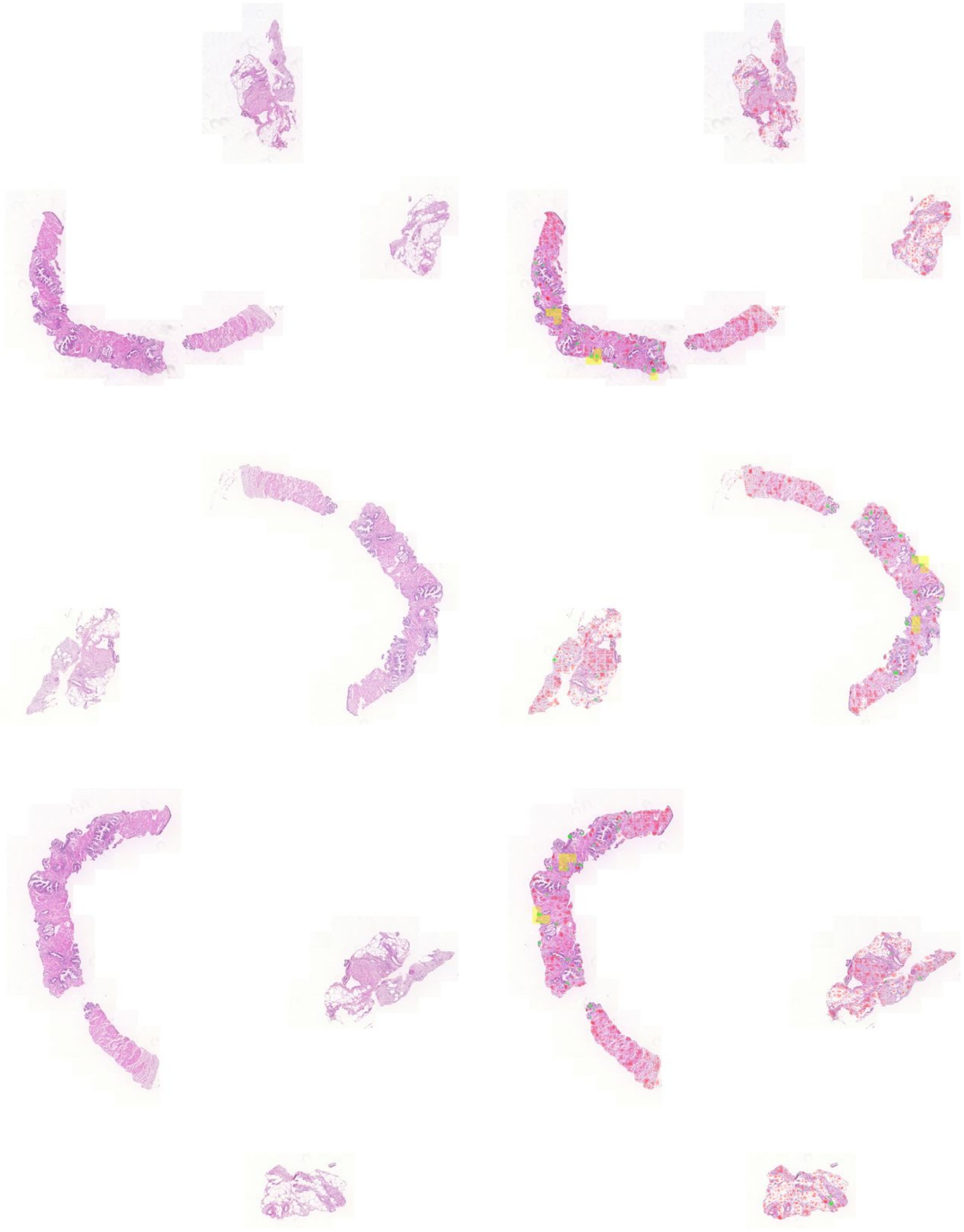

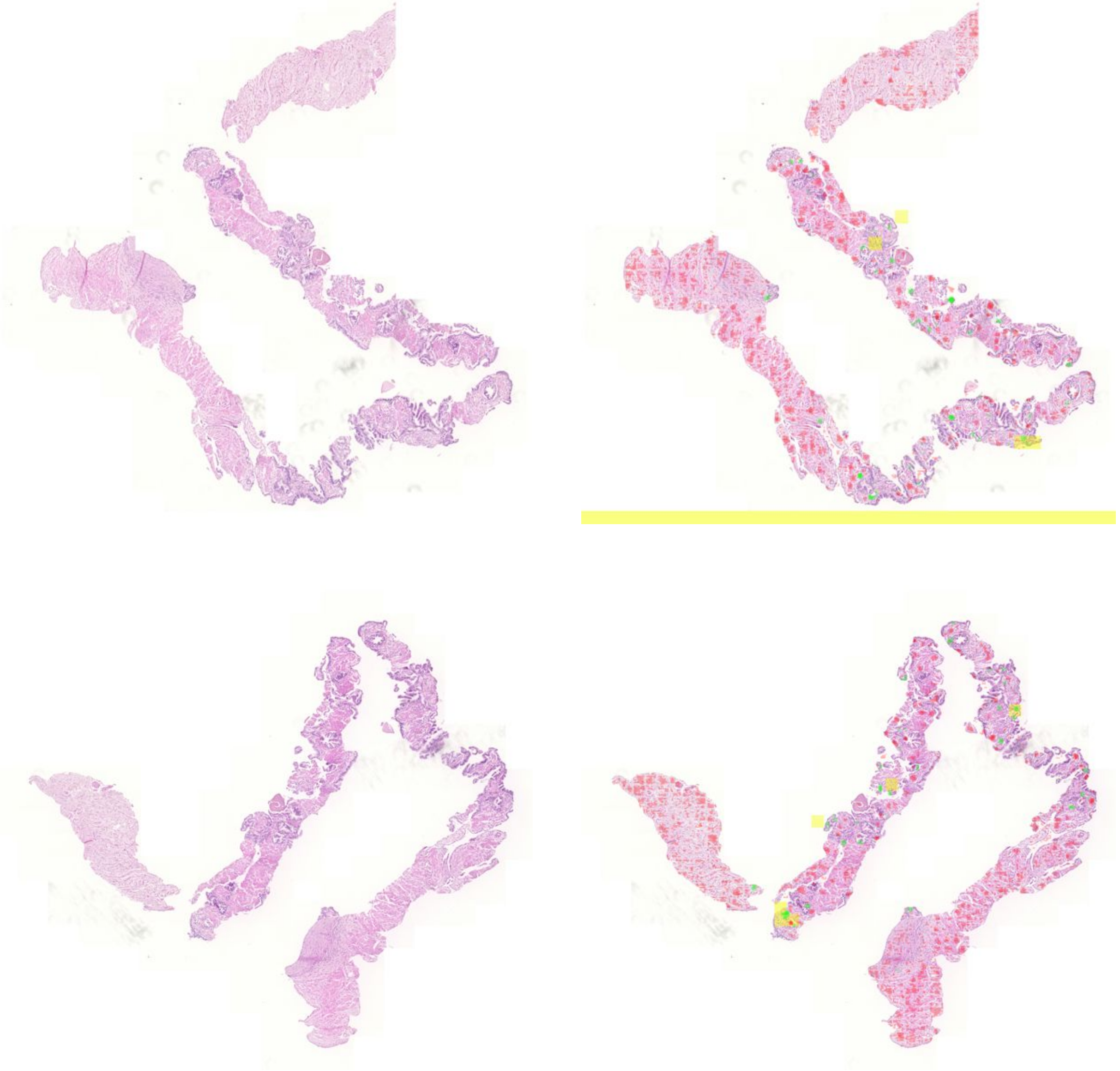

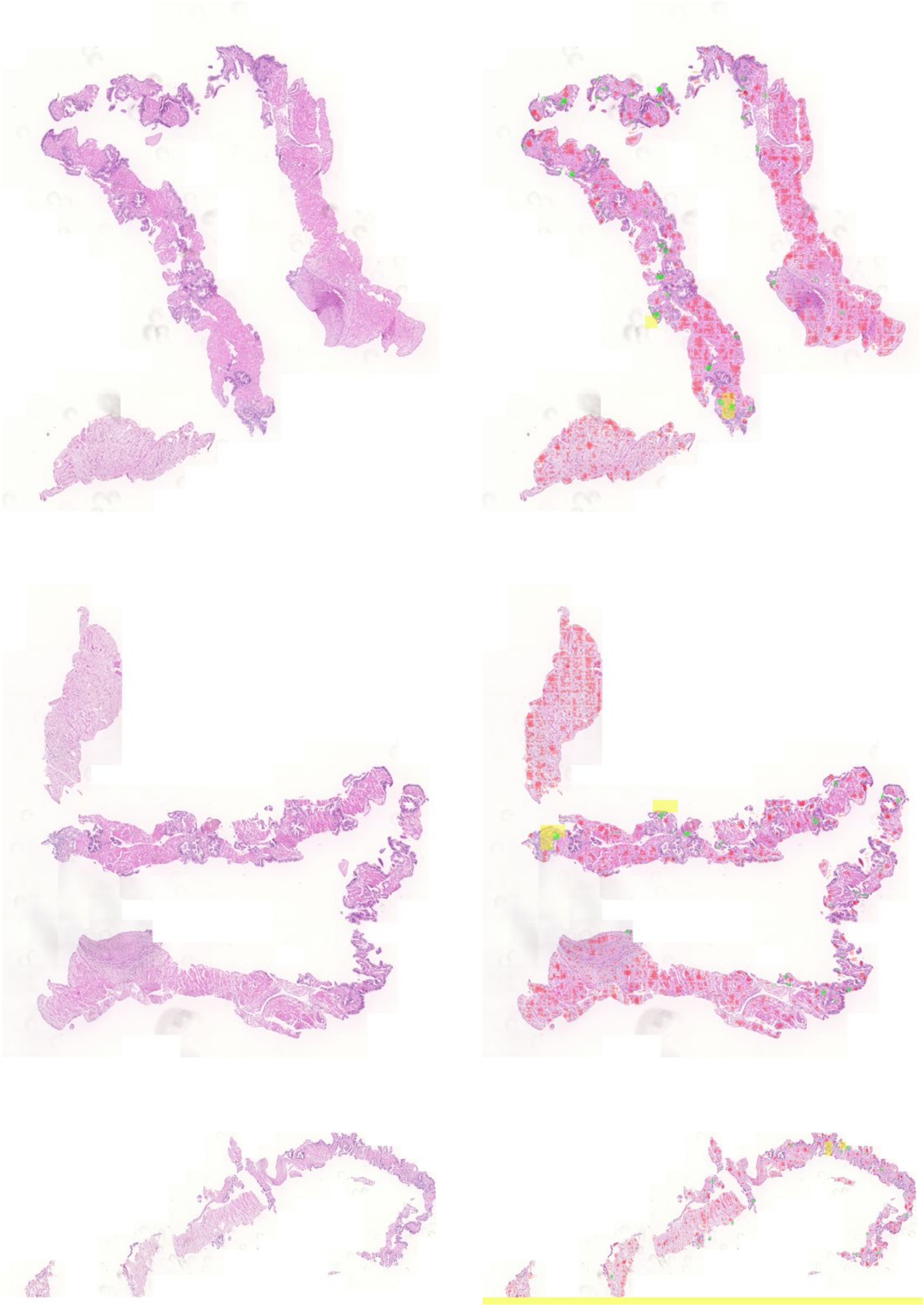

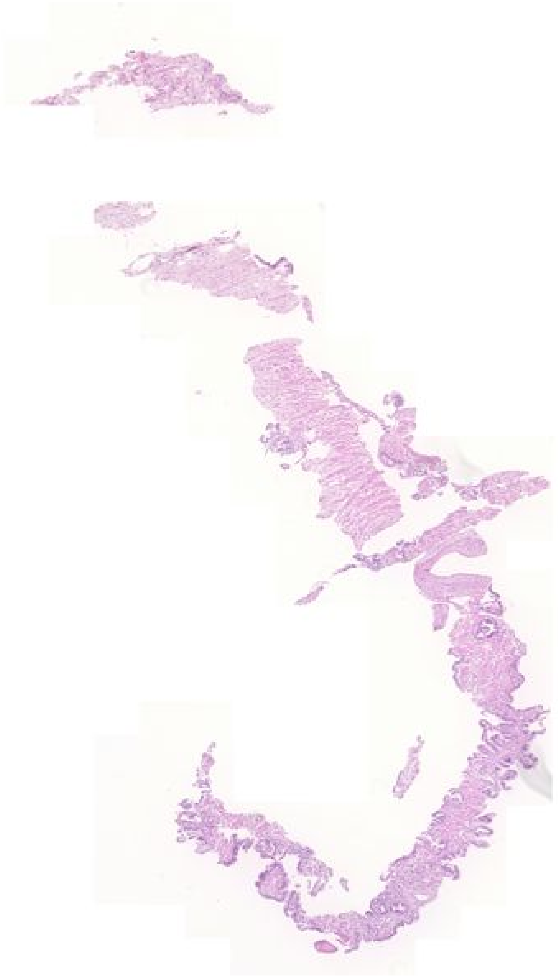

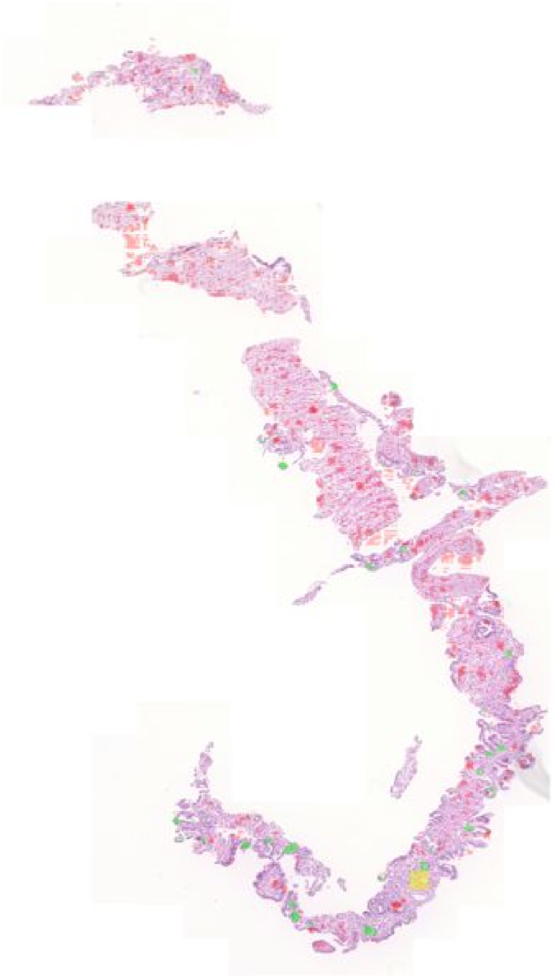

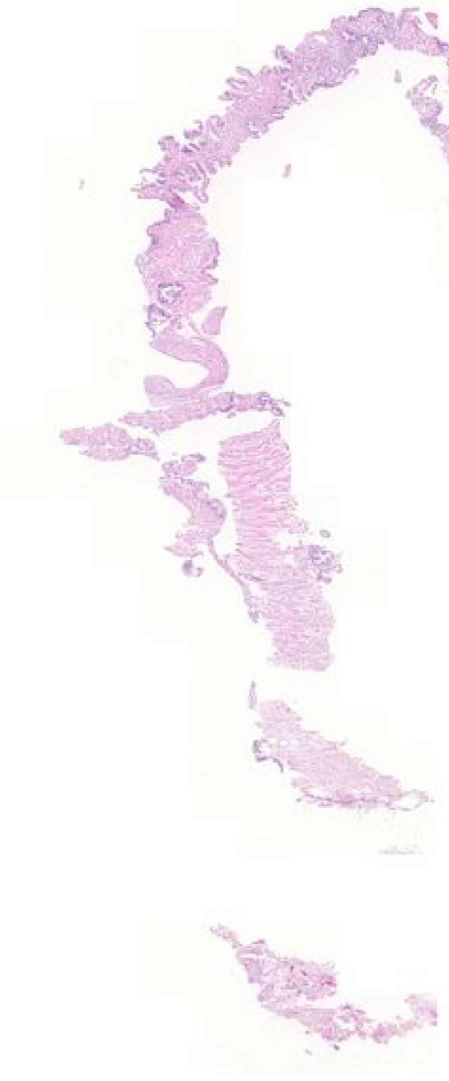

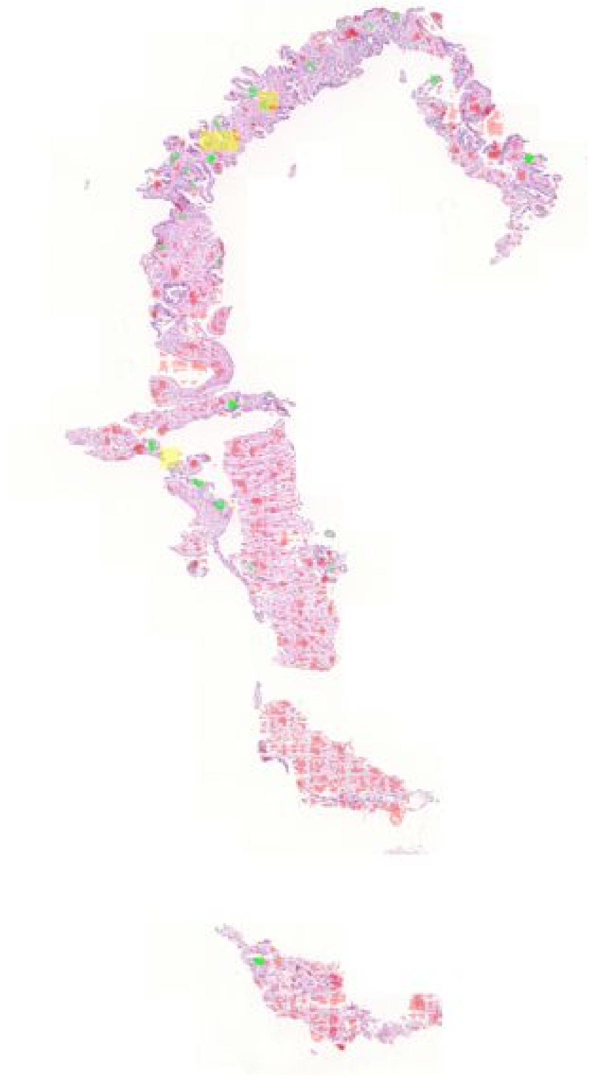

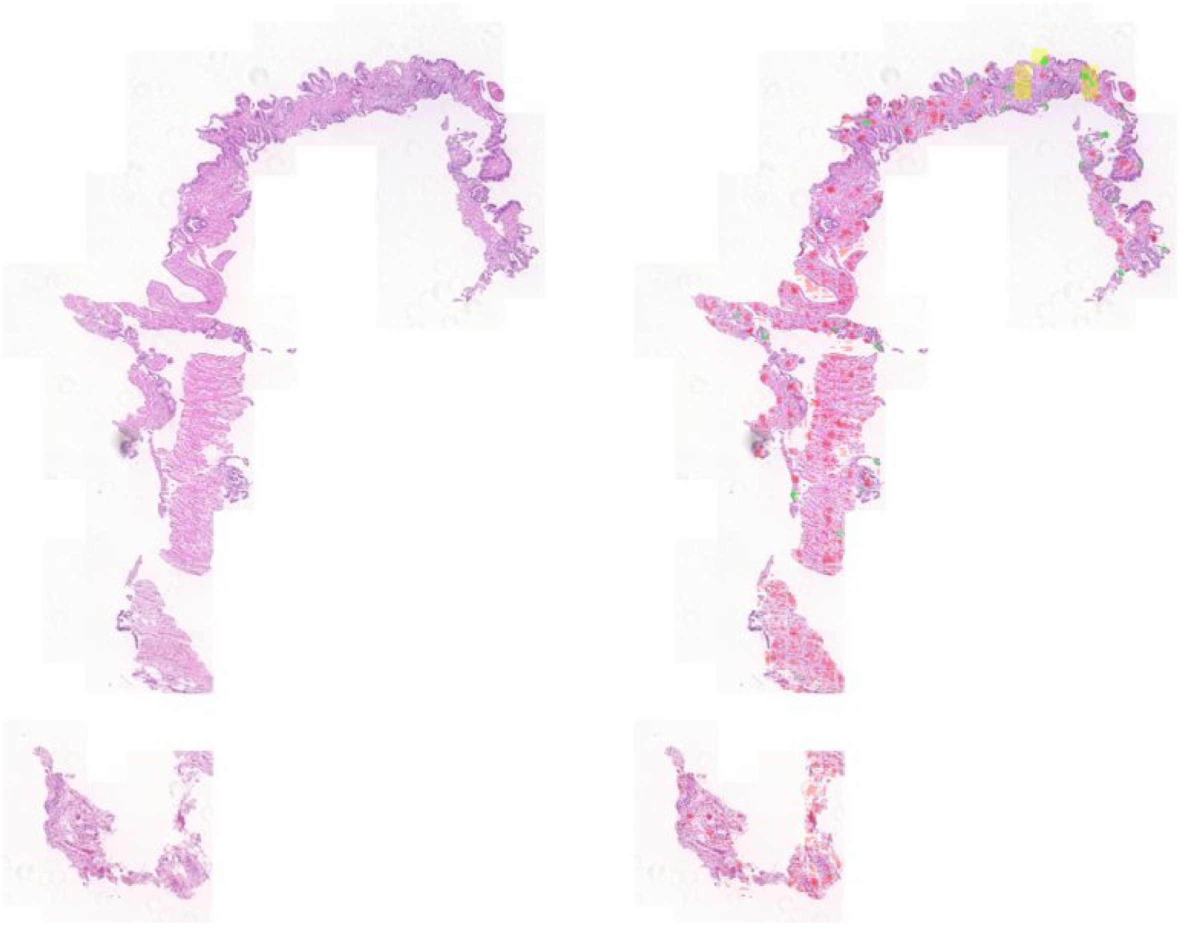

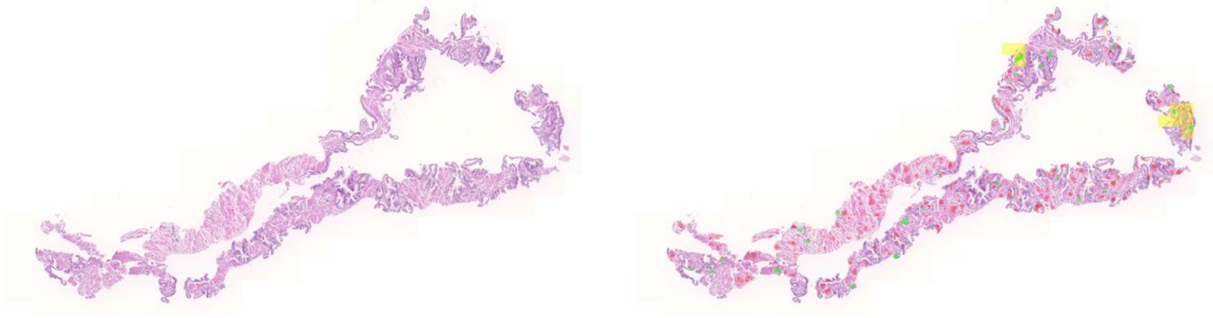

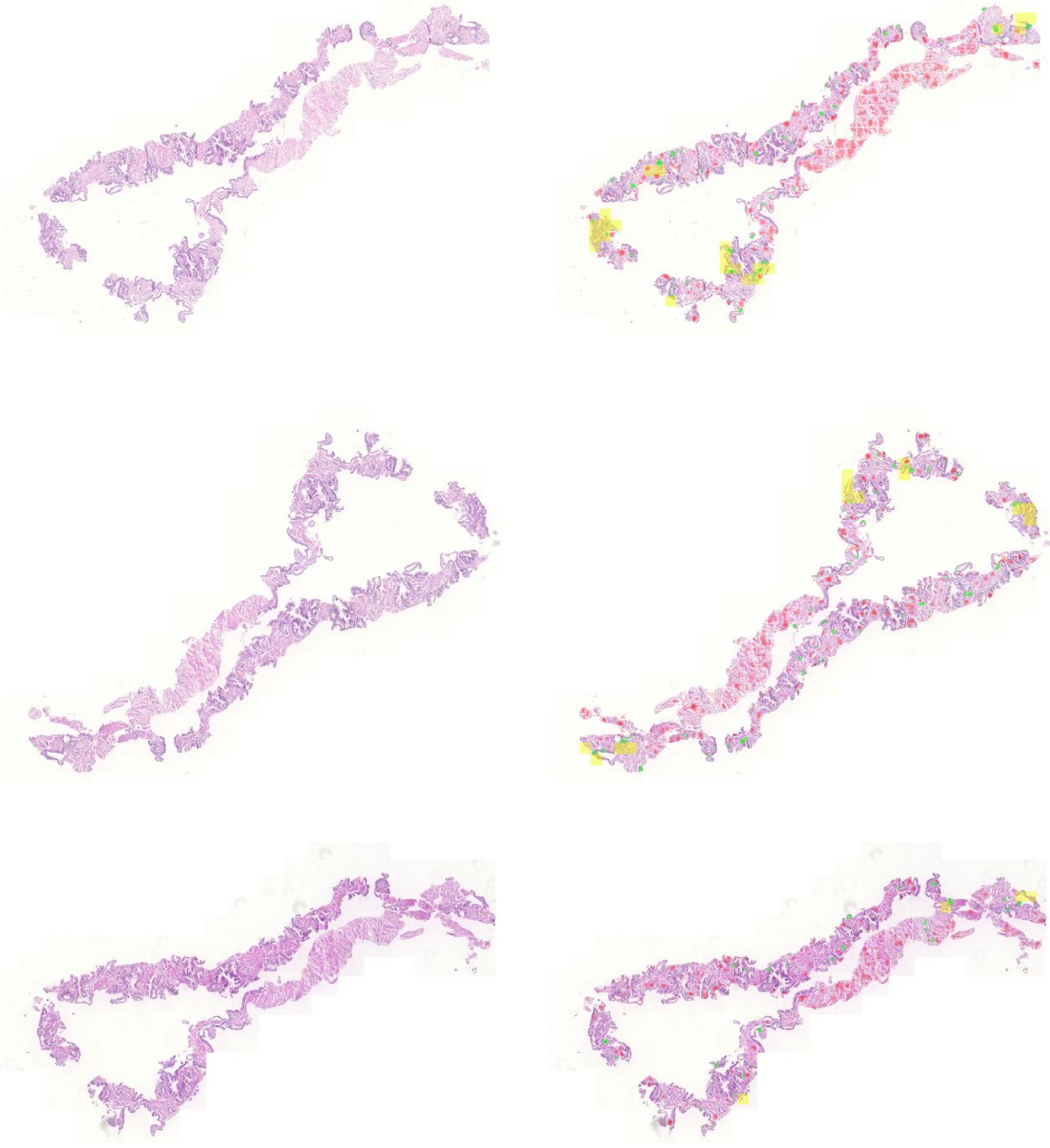

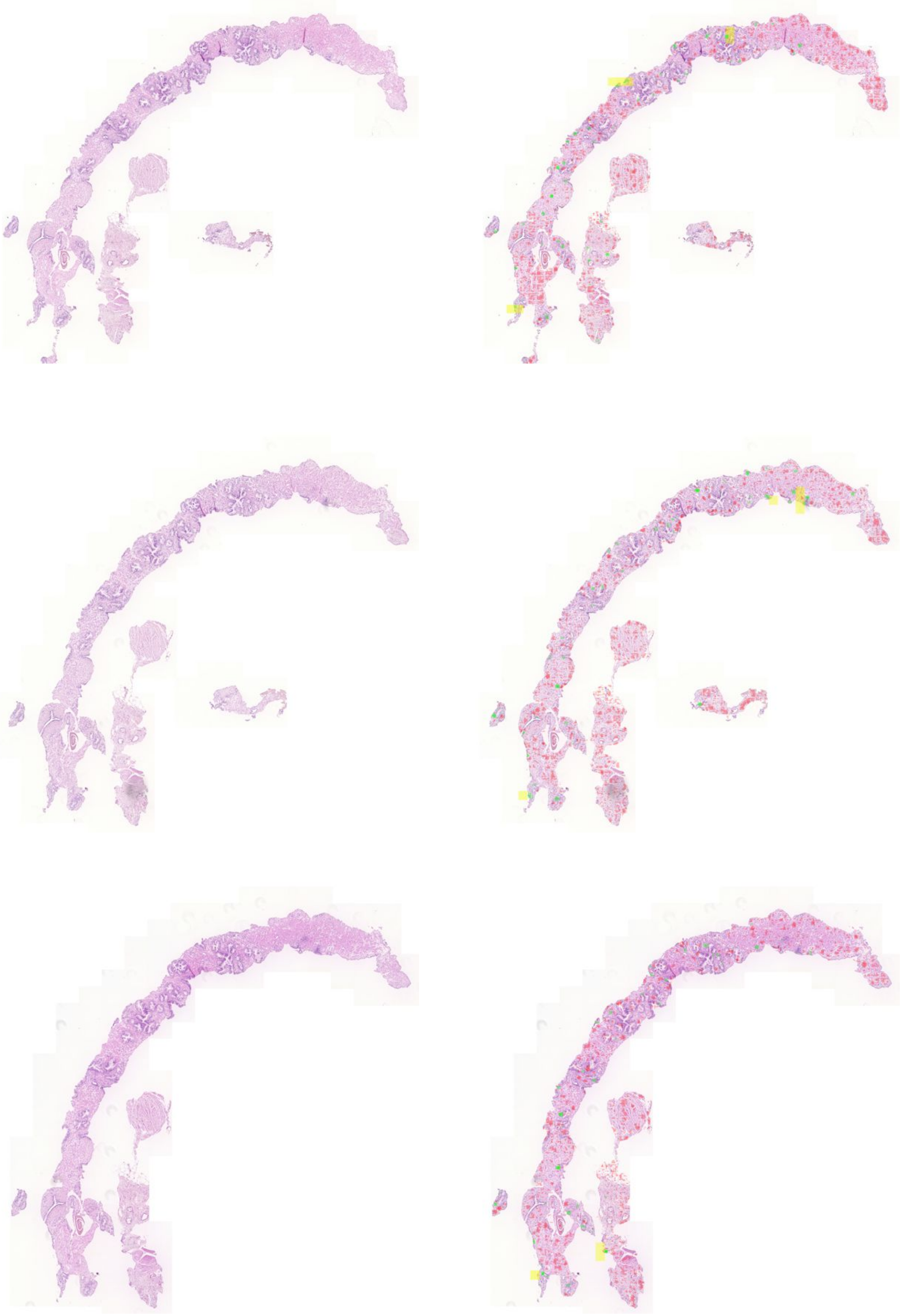

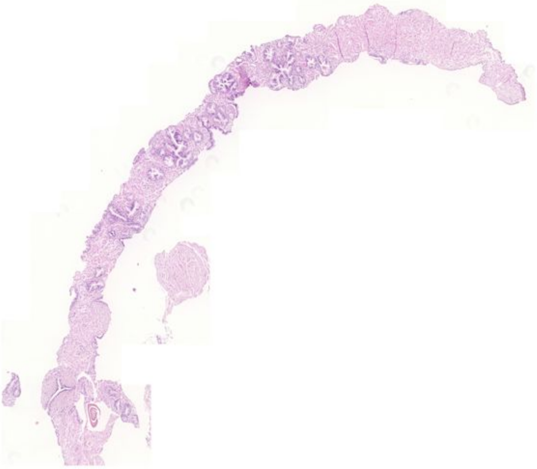
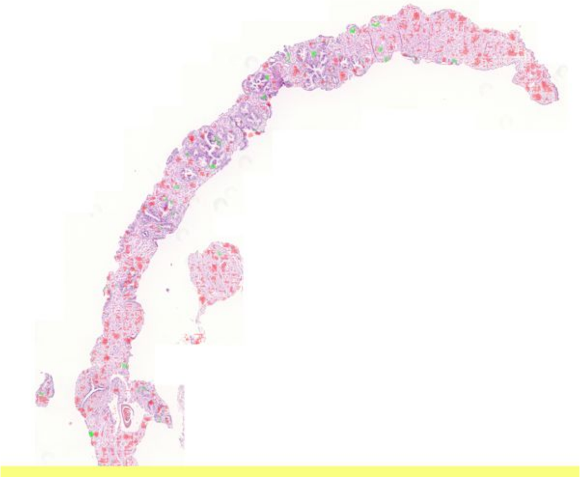
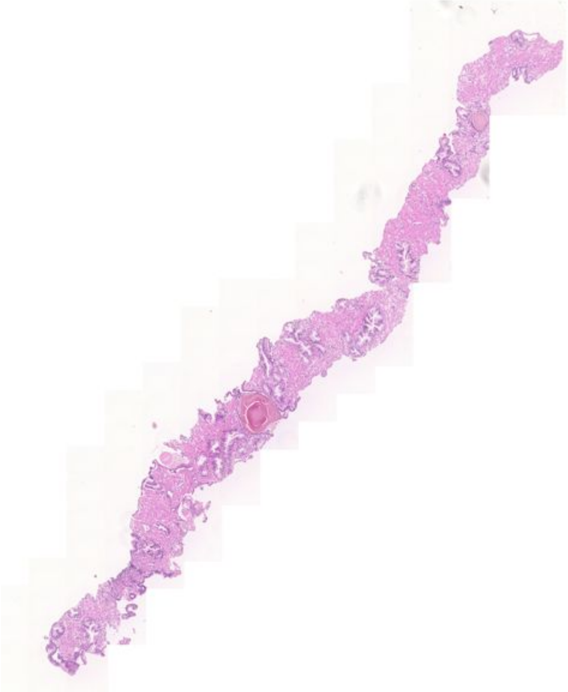
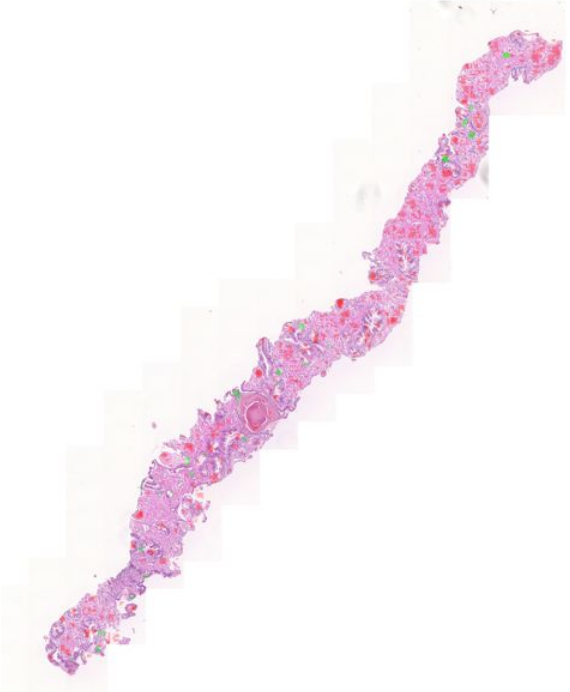
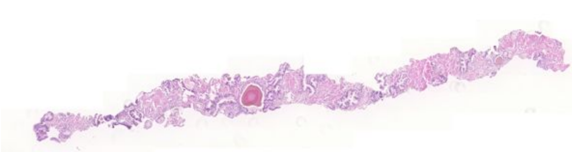
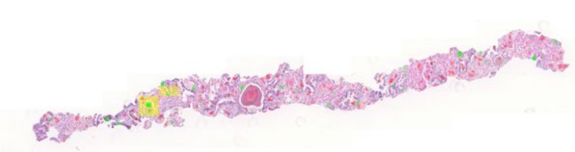
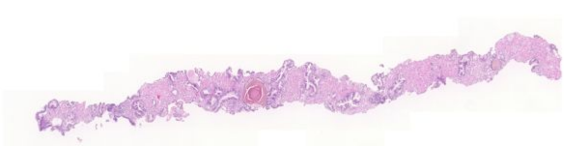
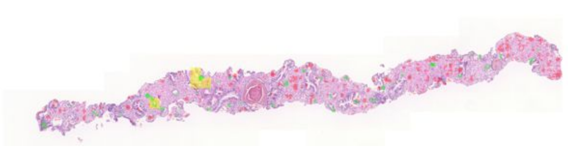

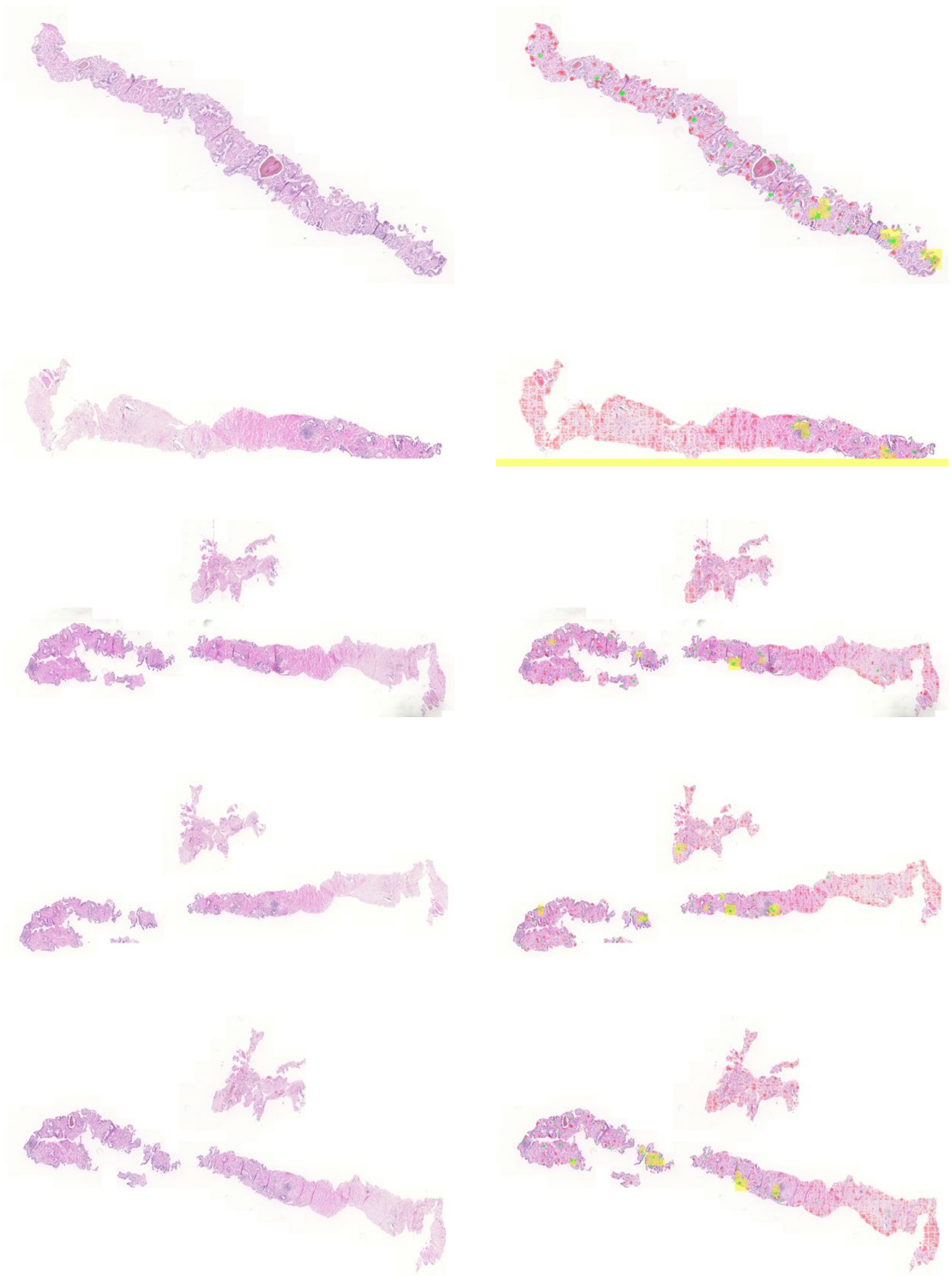

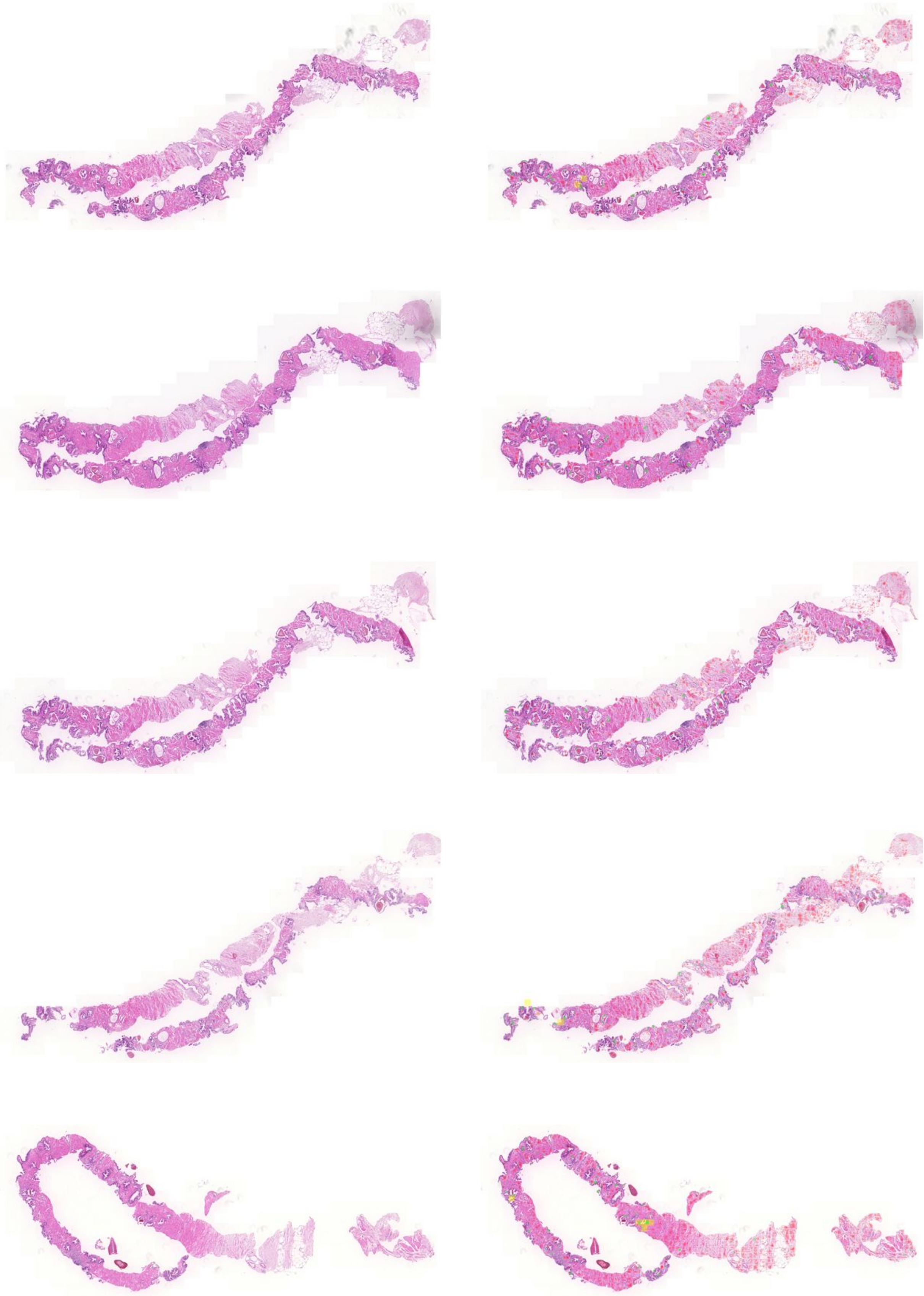

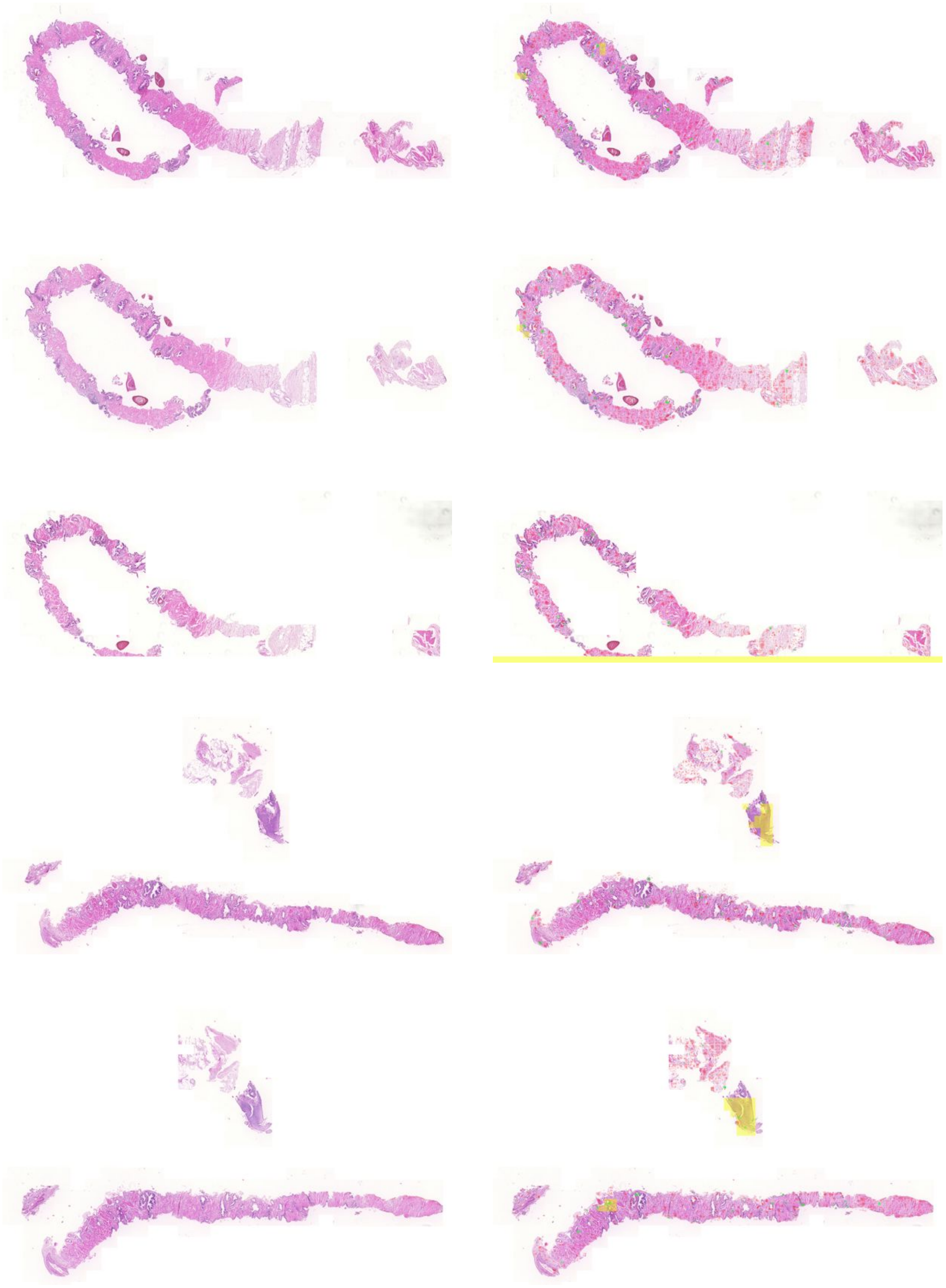

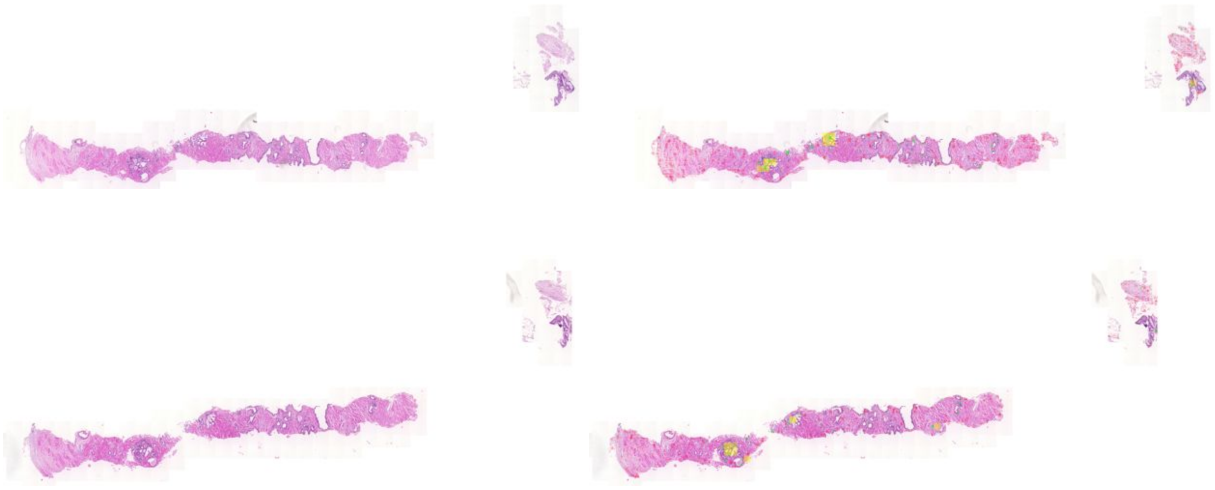
